# Elucidation of the biosynthetic pathway of reserpine

**DOI:** 10.1101/2025.01.23.634628

**Authors:** Jiaqing Cao, Jingxiao Zhong, Feng Li, Yindi Jiang

## Abstract

Reserpine is a landmark natural product that has profoundly influenced our understanding of neurotransmitter biology and cardiovascular medicine. Its complex structure, featuring a unique C3 β-configuration and five consecutive chiral centers, has inspired generations of synthetic chemists to develop innovative strategies for its construction. However, the biosynthetic logic underlying nature’s assembly of this intricate molecule has remained elusive. Here, we decipher the biosynthetic pathway of reserpine in *Rauvolfia verticillata*, revealing that α-configured strictosidine serves as the biosynthetic precursor. The crucial C3 β-configuration is established through a two-step enzymatic epimerization, orchestrated by a flavin-dependent oxidase and a NADPH-dependent reductase. The consecutive chiral centers are constructed through coordinated action of distinct enzyme families. Through the identification of eight biosynthetic enzymes, including those catalyzing late-stage methoxylation, we successfully reconstituted the biosynthesis of rauvomitorine G, a key intermediate in reserpine formation. This work unveils nature’s elegant approach to stereoselective synthesis of complex alkaloids and provides valuable biocatalytic tools for molecular functionalization.

## Introduction

Reserpine is a structurally complex monoterpene indole alkaloid (MIA) that has profoundly influenced our fundamental understanding of biomedicine and chemistry. Since its isolation from *Rauvolfia serpentina* in 1952, reserpine has been instrumental for deciphering the complex mechanisms of neurotransmitter storage and release, particularly regarding monoamine neurotransmitters such as serotonin and norepinephrine^1,2,3^. Its ability to deplete these neurotransmitters has provided groundbreaking insights into the pathophysiology of hypertension and various psychiatric disorders^4,5^. Furthermore, reserpine’s clinical application as antihypertensive and antipsychotic agents paved the way for the development of more targeted therapeutics, revolutionizing the treatment paradigms in cardiovascular and mental health medicine^6^.

Beyond its biomedical significance, the complex structure of reserpine has challenged and inspired generations of synthetic chemists, leading to numerous total syntheses and innovative synthetic methodologies^7^. The molecule’s pentacyclic skeleton features a unique C3β stereochemistry and five consecutive chiral centers densely packed in the E ring (Fig. 1a). Most total syntheses of reserpine begin with the construction of a pentasubstituted E-ring, typically achieved through various synthetic strategies including Diels-Alder reaction, double Michael addition, and radical cyclization (Fig. 1b)^8,9,10,11,12^. This elaborated E-ring subsequently undergoes Pictet-Spengler condensation with 6-methoxytryptamine to generate a pentacyclic intermediate bearing an α-configuration at C3. Access to the desired β-configuration at C3 has largely relied on R. B. Woodward’s landmark conformational control strategy, which involves locking the molecule in an unfavorable conformation through lactonization, followed by acid treatment to achieve late-stage C3 epimerization^8^. Despite these sophisticated strategies developed by synthetic chemists, the biosynthetic pathway by which nature constructs such complex natural products remains elusive.

**Fig. 1:**
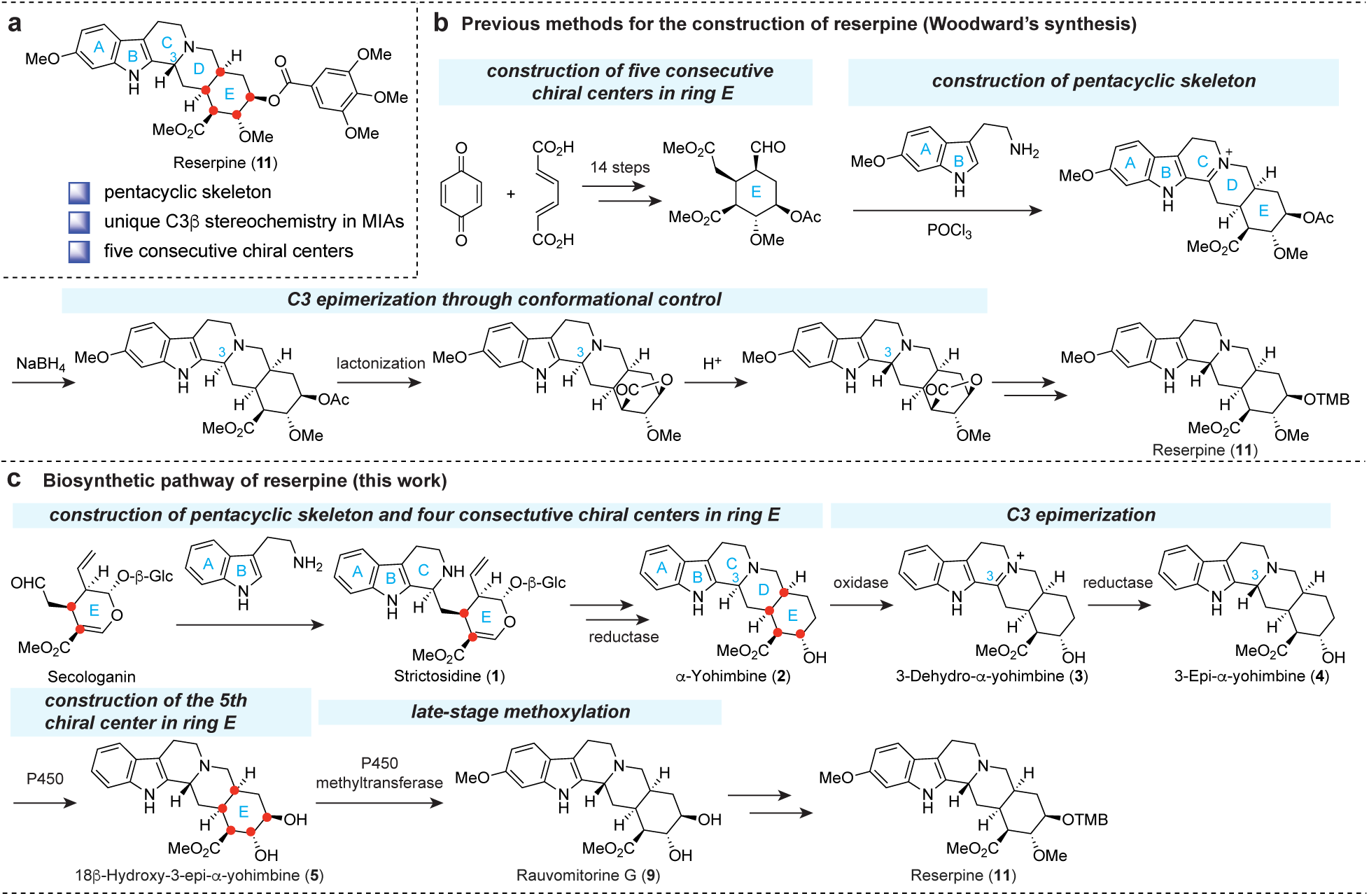
Structural features, synthetic history, and biosynthetic pathway of reserpine. **a**, Chemical structure of reserpine (**11**) highlighting its key structural features. **b**, Previous synthetic approaches exemplified by Woodward’s synthesis. **c**, The biosynthetic pathway of reserpine identified in this work. TMB represents 3,4,5-trimethoxybenzoyl group.

Isotope feeding studies have established tryptamine as the biosynthetic precursor of reserpine^13^. Its condensation with monoterpene-derived secologanin yields two diastereomers, vincoside and strictosidine^14^. The shared C3 configuration between vincoside and reserpine has historically suggested vincoside as the biosynthetic precursor of reserpine^7^. However, this proposal contradicts established MIA biosynthetic pathways, where strictosidine, not vincoside, functions as the common precursor^15^. This discrepancy raises an intriguing possibility that strictosidine might serve as the reserpine precursor, necessitating subsequent stereochemical inversion to establish the characteristic C3β configuration of reserpine.

Here we report the elucidation of the reserpine biosynthetic pathway in *R. verticillata* through an integrated approach combining transcriptome mining, heterologous expression, and biochemical characterization (Fig. 1c and 2). Our findings reveal that strictosidine, not vincoside, serves as the authentic precursor in reserpine biosynthesis. We uncovered the enzymatic basis for C3 epimerization in *Rauvolfia*, which proceeds through a two-step process: an FAD-dependent oxidase *Rv*YOO generates an imine intermediate, which is subsequently reduced by a medium-chain dehydrogenase/reductase (MDR) *Rv*DYR1 to yield the 3-epi product. Furthermore, we revealed nature’s elegant strategy for constructing consecutive chiral centers in ring E, where two stereocenters are established through stereoselective reduction by an MDR *Rv*YOS, and the final stereocenter is installed via cytochrome P450 monooxygenase *Rv*CYP71D820. Through the identification of eight biosynthetic enzymes, including P450 monooxygenase *Rv*CYP72A270 and *O*-methyltransferase *Rv*11OMT for late-stage methoxylation, we successfully reconstituted rauvomitorine G, a crucial intermediate in reserpine biosynthesis. This work advances our understanding of complex alkaloid biosynthesis, revealing both the enzymatic logic underlying reserpine biosynthesis and providing valuable biocatalytic tools for selective functionalization of complex molecular scaffolds.

**Fig. 2:**
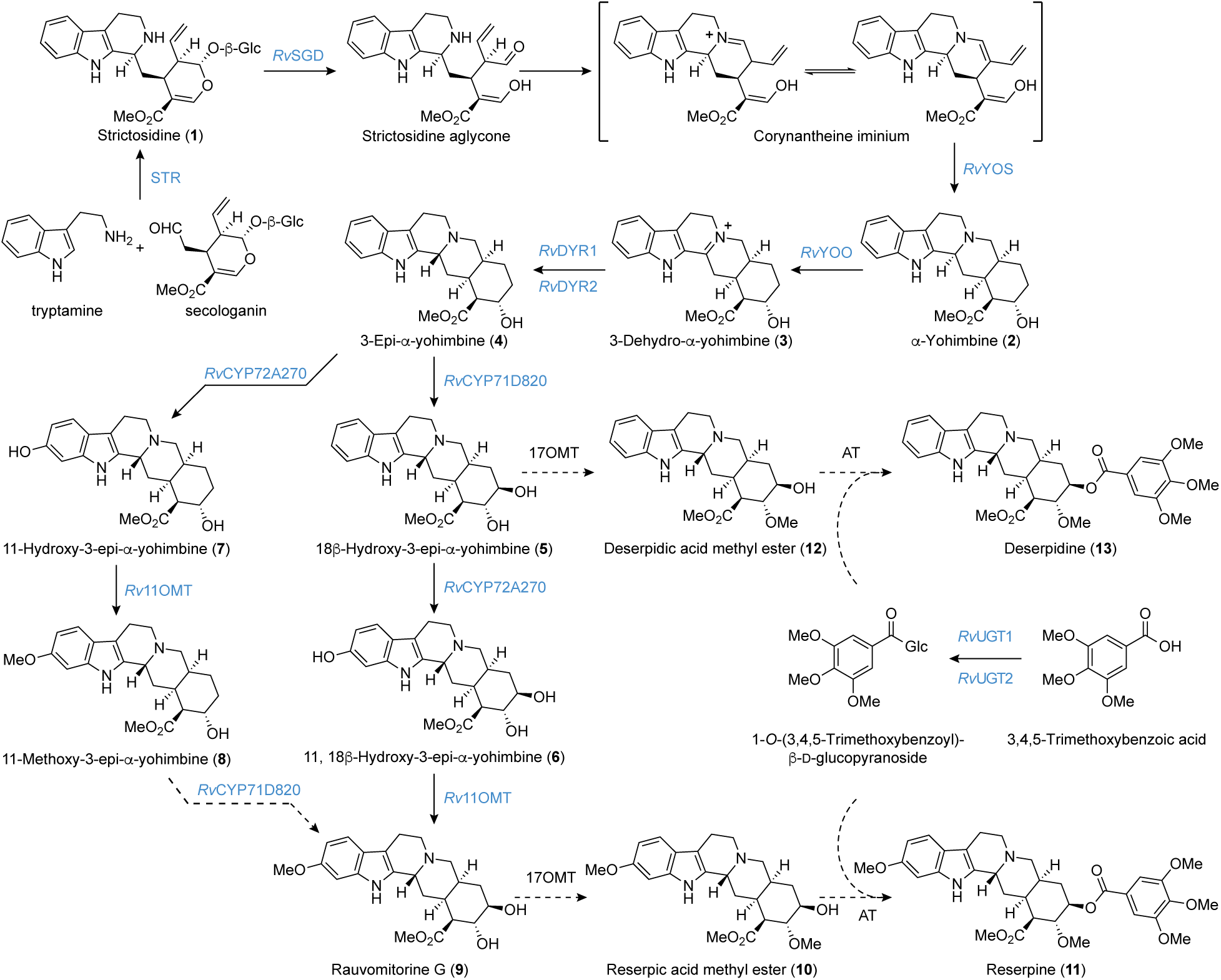
The biosynthetic pathway of reserpine in *R. verticillata*. Solid arrows indicate characterized steps; dashed arrows indicate proposed steps. Names of characterized enzymes are shown in blue. AT represents acyltransferase.

## Results

To elucidate the biosynthesis of reserpine, we first examined the distribution of monoterpene indole alkaloids (MIAs) in *Rauvolfia verticillata*, a medicinal plant used in China as a source of reserpine. Liquid chromatography-mass spectrometry (LC-MS) analysis of tissue extracts revealed several key MIAs, including reserpine (**11**), deserpidine (**13**), α-yohimbine (**2**), and 3-epi-α-yohimbine (**4**) (Supplementary Fig. 1). Having confirmed the presence of these MIAs across different tissues, we sought to identify the genes responsible for their biosynthesis. To facilitate biosynthetic gene discovery, we generated tissue-specific transcriptome datasets in triplicate.

### Formation of α-yohimbine through strictosidine pathway

A distinct structural feature of reserpine is its β-configuration at C3, which contrasts with the α-configuration predominant in most MIAs. The isolation of vincoside and vincoside lactam, which share this β-configuration at C3, from *Rauvolfia* species has long suggested vincoside as the biosynthetic precursor of reserpine (Supplementary Fig. 2)^7,16^. However, the stepwise conversion from vincoside to reserpine remains unsolved.

Strictosidine (**1**), a stereoisomer of vincoside and the common MIA precursor, is stereospecifically synthesized from tryptamine and secologanin by strictosidine synthase (STR). The subsequent deglycosylation by strictosidine glucosidase (SGD) generates a reactive iminium intermediate, which various reductases can transform into diverse indole alkaloids^15^. While *Rauvolfia* SGD showed specificity for **1** deglycosylation, it exhibits no activity towards vincoside^17^. Notably, *Catharanthus roseus* SGD (*Cr*SGD) can deglycosylate both **1** and vincoside, suggesting the possible existence of an unidentified vincoside-specific SGD in *Rauvolfia*^18^.

The prevalence of C3 β-configured intermediates in *Rauvolfia* is evidenced by numerous isolated compounds, including vincoside lactam, 3-epi-α-yohimbine (**4**), rauvomitorine G (**9**), deserpidic acid methyl ester (**12**), and rescinnamine, with **4** representing the structurally simplest intermediate (Supplementary Fig. 2)^19,20,21^. Recent studies have elucidated the biosynthesis of α-yohimbine (**2**), the C3 epimer of **4**, in *R. tetraphylla*^22^. This pathway proceeds via strictosidine (**1**), involving SGD-catalyzed formation of a reactive imine intermediate and subsequent reduction by yohimbane synthase (YOS), a medium-chain dehydrogenase/reductase (MDR).

These findings led us to propose two potential biosynthetic pathways for 3-epi-α-yohimbine (**4**) (Supplementary Fig. 3). Pathway A involves condensation of tryptamine and secologanin to form vincoside (**1a**), followed by conversion to **4** through an unidentified SGD and an MDR. Alternatively, pathway B begins with STR-mediated condensation of tryptamine and secologanin to produce strictosidine (**1**)^23^, which is converted to α-yohimbine (**2**) by characterized SGD and YOS enzymes^22,24^. Subsequently, stereochemical inversion of C-3 of **2** could be catalyzed by an unidentified epimerase to yield **4**.

To investigate pathway A, we chemically synthesized a mixture of strictosidine (**1**) and vincoside (**1a**). Five putative candidate genes potentially capable of vincoside deglycosylation were identified in the *R. verticillata* transcriptome using *Cr*SGD as a query. Each gene was cloned from *R. verticillata* cDNA and expressed in *Nicotiana benthamiana*. LC-MS analysis revealed that while *Rv*SGD1 effectively consumed strictosidine (**1)**, vincoside remained unchanged in all SGD candidate assays (Supplementary Fig. 4), suggesting that vincoside is not the source of 3-*epi*-α-yohimbine (**4**).

We then explored pathway B through reconstituting α-yohimbine (**2**) in planta (Fig. 3a). Previous work in *R. tetraphylla* showed that YOS reduces strictosidine aglycone to form a mixture of yohimbine-type alkaloids, including α-yohimbine (**2**), yohimbine (**2a**), and corynanthine (**2c**)^22^. BLAST analysis of the *R. verticillata* transcriptome using *Rt*YOS as a query revealed a candidate gene, *Rv*YOS, sharing 94.4% sequence identity. To assess functional conservation, we coexpressed *Rv*YOS with *Catharanthus roseus* STR (*Cr*STR) and *Rv*SGD in *N. benthamiana*. Following infiltration of tryptamine and secologanin, this system successfully produced both **2** and **2a** (Supplementary Fig. 5). The formation of these products was further confirmed *in vitro* using purified *Cr*STR, *Rv*SGD, and *Rv*YOS enzymes (Supplementary Fig. 5). Although *Rv*YOS generated **2**, its lack of product specificity led us to screen all MDRs sharing over 30% sequence identity for enzymes that could selectively produce **2** (Fig. 3b). As none exhibited the desired specificity, we proceeded with a chemical standard of **2** for subsequent experiments.

**Fig. 3:**
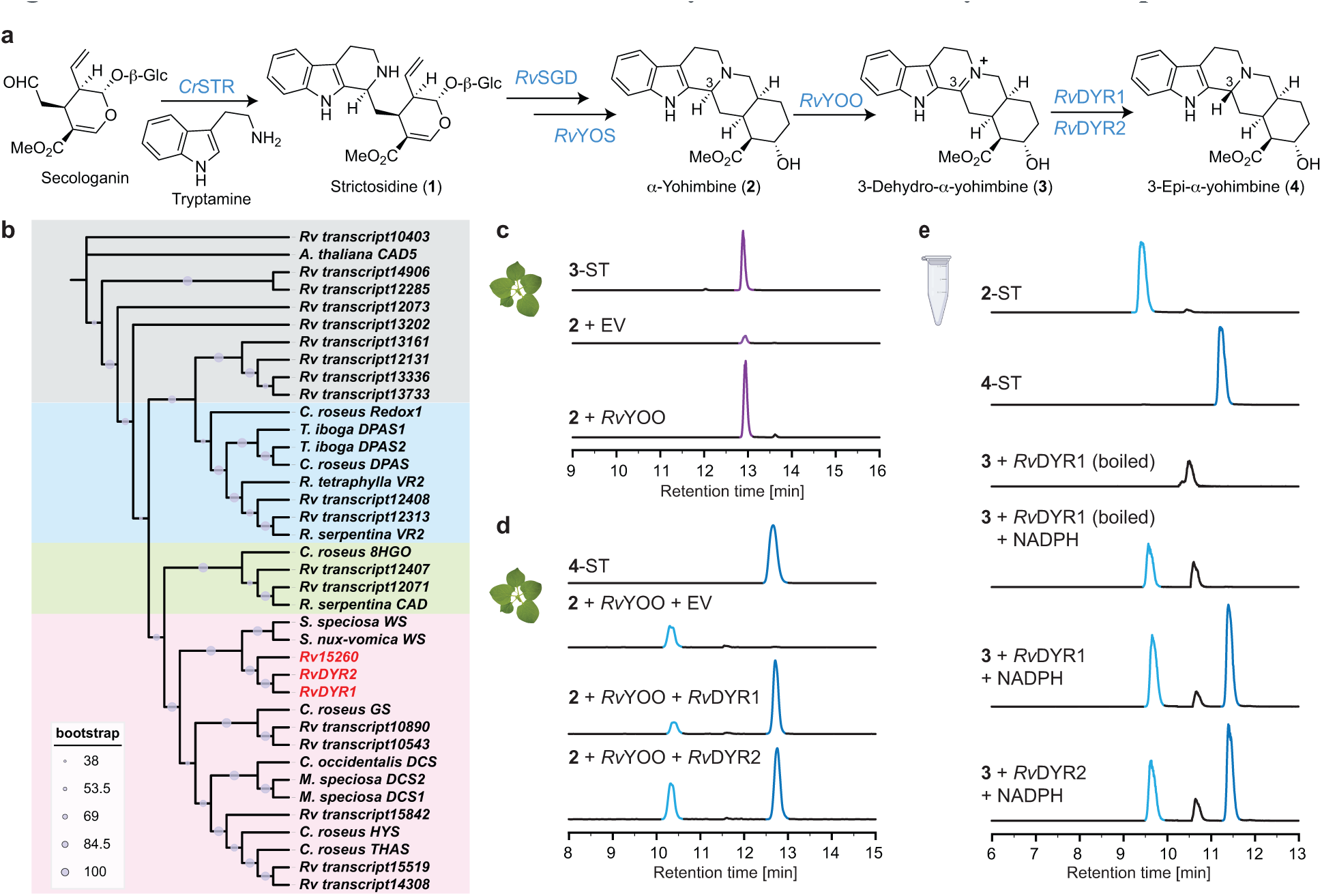
Identification and characterization of enzymes involved in α-yohimbine epimerization. **a**, Proposed biosynthetic pathway from tryptamine and secologanin to 3-epi-α-yohimbine (**4**) via α-yohimbine (**2**) and 3-dehydro-α-yohimbine (**3**) intermediates. **b**, Phylogenetic analysis of medium-chain dehydrogenase/reductase (MDR) enzymes, with colored boxes indicating different reaction types catalyzed by characterized enzymes. Bootstrap values are shown as circles of varying sizes. **c**, LC-MS analysis showing oxidation of α-yohimbine (**2**) by *Rv*YOO expressed in *N. benthamiana* leaves compared to empty vector (EV) control and standard (ST) of **3**. **d**, LC-MS analysis demonstrating conversion of α-yohimbine (**2**) to 3-epi-α-yohimbine (**4**) by co-expression of *Rv*YOO with *Rv*DYR1 or *Rv*DYR2 in *N. benthamiana*. **e**, In vitro characterization of purified *Rv*DYR1 and *Rv*DYR2 showing stereoselective reduction of synthetic 3-dehydro-α-yohimbine (**3**) to 3-epi-α-yohimbine (**4**) in the presence of NADPH.

### Enzymatic epimerization of α-yohimbine (2)

We next investigated the enzymatic basis of α-yohimbine (**2**) epimerization to 3-epi-α-yohimbine (**4**). Epimerization could be catalyzed by epimerases, which are crucial in the primary metabolism of carbohydrates and amino acids, as well as in secondary metabolite biosynthesis^25,26^. During morphine biosynthesis in opium poppy, a reticuline epimerase (REPI) catalyzed the stereoconversion of *S*-reticuline to *R*-reticuline through sequential oxidation and reduction with its two domains^27^. The N-terminal cytochrome P450 domain oxidizes *S*-reticuline to an iminium intermediate, which was stereo-specifically reduced by the C-terminal aldo-keto reductase (AKR) domain to *R*-reticuline in a NADPH-dependent manner (Supplementary Fig. 6). Thus, we speculated that epimerization of α-yohimbine (**2**) happened in a similar manner that **2** was oxidized to iminium intermediate 3-dehydro-α-yohimbine (**3**), followed by reduction from the convex face to give 3-epi-α-yohimbine (**4**) (Fig. 3a).

Our search for REPI homologs in the *R. verticillata* transcriptome failed to identify candidates containing both P450 and AKR domains. Although the CYP450 domain in REPI belongs to the CYP82 family, expression of seven CYP82 gene candidates from *R. verticillata* in *N. benthamiana* leaves co-infiltrated with **2** yielded no detectable oxidized products. We then considered alternative oxidases other than cytochrome P450s. In vinblastine biosynthesis in *C. roseus*, the flavin-dependent precondylocarpine acetate synthase (*Cr*PAS) oxidizes the tertiary amine in stemmadenine acetate to an iminium intermediate^28^. This led us to investigate PAS homologs as potential oxidants for **2**. Among eight *Cr*PAS homologs identified in the *R. verticillata* transcriptome, one candidate sharing 58.0% identity with *Cr*PAS, which we named α-yohimbine oxidase (*Rv*YOO), catalyzed the conversion of **2** to a product with an exact mass of [M+H]^+^ 353.1857, consistent with an oxidized product of **2** (calculated for C21H25N2O3^+^: 353.1860) (Fig. 3c). The structure was confirmed using synthetic 3-dehydro-α-yohimbine (**3**), which showed identical retention time, exact mass, and MS/MS fragmentation patterns (Supplementary Fig. 7). Further investigation of *Rv*YOO’s substrate scope through in planta expression and infiltration with α-yohimbine stereoisomers revealed that α-yohimbine (**2**), yohimbine (**2a**), and *β*-yohimbine (**2b**) were oxidized, while corynanthine (**2c**) and 3-epi-α-yohimbine (**4**) were not substrates (Supplementary Fig. 7). Notably, the inability of *Rv*YOO to oxidize **4** suggests that once formed, **4** cannot be coverted back to **3** through the same enzymatic mechanism, indicating the irreversible nature of this transformation.

For the subsequent reduction of 3-dehydro-α-yohimbine (**3**), we focused on MDR enzymes, which are known to catalyze iminium reduction in MIA biosynthesis^29,30,31,32^. A well-characterized example is tetrahydroalstonine synthase (THAS) in *C. roseus* that performs 1,2-reduction of iminium strictosidine aglycone to yield tetrahydroalstonine^33^. Co-expression of either of the two 3-dehydro-α-yohimbine reductases (*Rv*DYR1 and *Rv*DYR2) with *Rv*YOO in *N. benthamiana* infiltrated with α-yohimbine (**2**) resulted in the formation of a product that co-eluted with authentic 3-epi-α-yohimbine (**4**) standard (Fig. 3d). However, due to incomplete substrate conversion by *Rv*YOO, we observed a mixture of **2** and **4** in *N. benthamiana* leaves expressing active MDRs, which prevented definitive conclusions about the stereoselectivity of *Rv*DYR1 and *Rv*DYR2 toward **3**.

To unambiguously determine stereoselectivity, we expressed and purified N-terminal His6-tagged *Rv*DYR1 and *Rv*DYR2 for *in vitro* characterization (Supplementary Fig. 8). Incubation of synthetic **3** with purified enzymes and NADPH predominantly yielded 3-epi-α-yohimbine (**4**) over α-yohimbine (**2**) (Fig. 3e). Notably, **2** could be generated by NADPH alone, while the formation of **4** required both NADPH and active *Rv*DYR enzymes, indicating *Rv*DYR enzymes’ preference for C3β stereoselectivity. Interestingly, despite high sequence similarity among *Rv*DYR1, *Rv*DYR2, and *Rv*DYR15260, *Rv*DYR15260 showed no activity toward the substrate (Supplementary Fig. 9). This lack of activity might be attributed to certain non-conserved residues among these three enzymes. While we were preparing this manuscript, two independent studies reported similar epimerization mechanisms for MIAs in plants^34,35^. To further understand the molecular basis of substrate recognition and and stereochemical control, structural studies including X-ray crystallography and site-directed mutagenesis are currently underway.

### Stereospecific C18 hydroxylation establishing the fifth chiral center in ring E

The next stage of reserpine biosynthesis involves two hydroxylation events at C11 and C18 as well as two methylation events at O11 and O17. The isolation of 18β-hydroxy-3-epi-α-yohimbine (**5**) and rauvomitorine G (**9**) from *Rauvolfia* suggested that stereospecific hydroxylation at C18 of **4** occurs first, followed by methoxylation at C11 (Fig. 4a)^19,20^. Given that no C18 hydroxylase has been previously reported, sequence homology-based approaches were not feasible for identifying the responsible enzyme. Therefore, we employed a co-expression analysis strategy using *Rv*DYR1 as a bait, as the expression of upstream biosynthetic genes showed similar tissue-specific expression patterns and high correlation (Fig. 4b and 4c). We selected 30 P450 enzymes (Pearson’s r>0.7) for functional characterization (Supplementary Fig 10). Transient expression of *Rv*CYP71D820 in *N. benthamiana* leaves infiltrated with **4** resulted in the formation of a new product with an exact mass (*m/z* [M+H]^+^ = 371.1963) matching hydroxylated **4** (calculated for C21H27N2O4^+^: 371.1965) (Fig. 4d). We validated this finding *in vitro* using *Rv*CYP71D820 in yeast microsomal fractions, which efficiently converted **4** to the same product observed in planta (Fig. 4d). MS/MS analysis of this product revealed an intact indole moiety (*m/z* [M+H]^+^ = 144.0807) (Supplementary Fig. 11), indicating hydroxylation occurred outside the indole ring. The product’s identity was confirmed using a chemically synthesized standard of **5**, prepared from deserpidine (**13**) in two steps, which showed identical retention time, exact mass, and MS/MS fragmentation patterns (Supplementary Fig. 11). These results establish *Rv*CYP71D820 as the enzyme responsible for stereospecific C18 hydroxylation of **4**, completing the construction of five consecutive chiral centers in the E ring of reserpine.

**Fig. 4:**
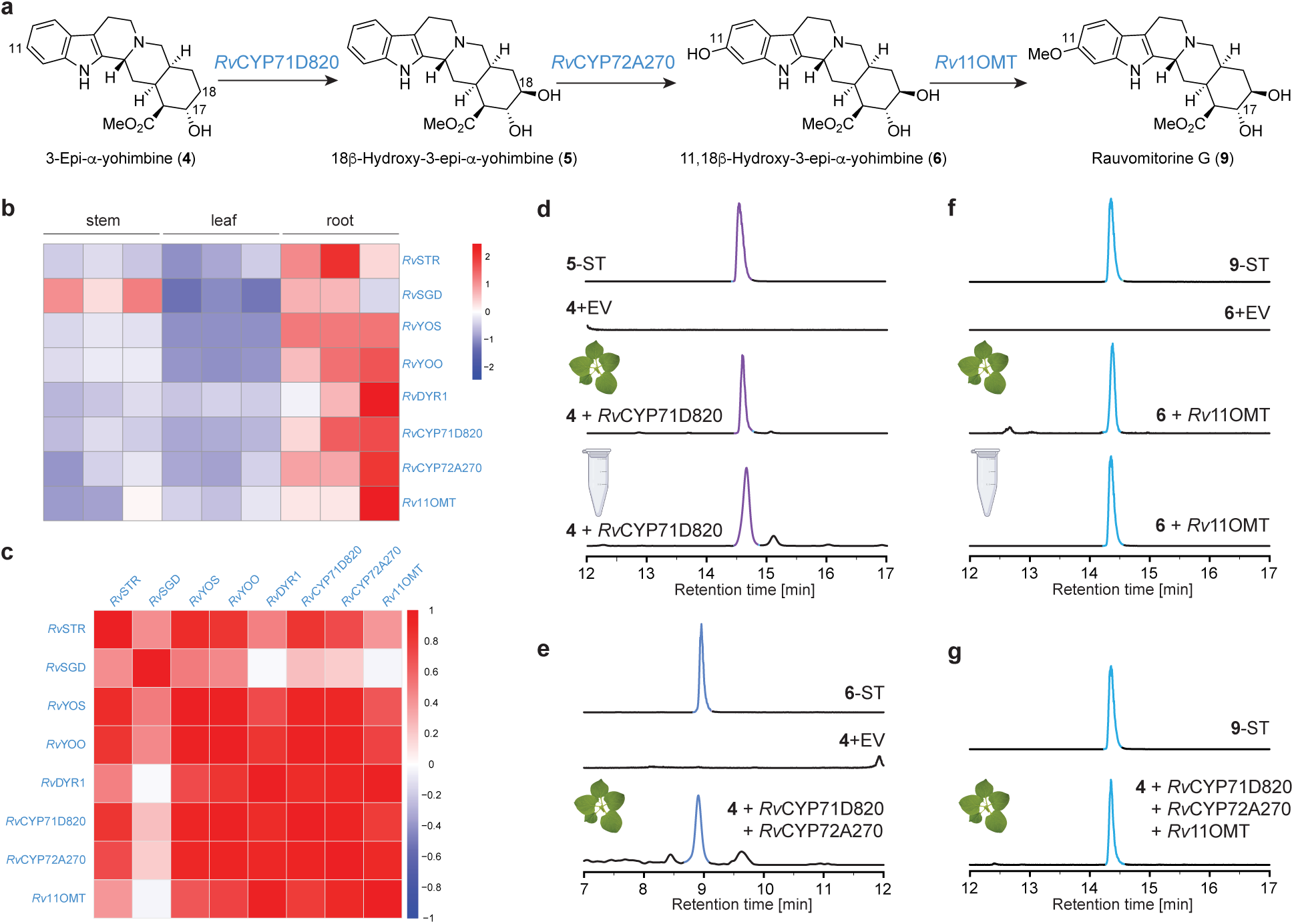
Discovery and characterization of enzymes responsible for C11 and C18 hydroxylation and *O*-methylation. **a**, Proposed biosynthetic pathway from 3-epi-α-yohimbine (**4**) to rauvomitorine G (**9**) via sequential hydroxylation and *O*-methylation reactions. **b**, Tissue-specific expression patterns of biosynthetic genes in *R. verticillata* stem, leaf, and root tissues; **c**, Correlation matrix showing Pearson correlation coefficients between expression levels of identified biosynthetic genes. **d**, LC-MS analysis of *Rv*CYP71D820-catalyzed C18 hydroxylation of **4** in *N. benthamiana* (leaf icon) and *in vitro* using yeast microsomes (tube icon), compared with synthetic standard of **5**. **e**, LC-MS analysis showing formation of 11,18β-dihydroxy-3-epi-α-yohimbine (**6**) through co-expression of *Rv*CYP71D820 and *Rv*CYP72A270 *in N. benthamiana*, compared with synthetic standard of **6**. **f**, LC-MS analysis of *Rv*11OMT-catalyzed *O*-methylation of **6** in *N. benthamiana* and *in vitro* using purified enzyme, compared with synthetic standard of **9**. **g**, LC-MS analysis of the reconstitution of **9** from **4** through co-expression of *Rv*CYP71D820, *Rv*CYP72A270, and *Rv*11OMT in *N. benthamiana*. EV, empty vector control; ST, synthetic standard.

### Enzymatic late-stage methoxylation

We next sought to identify the oxidase and methyltransferase responsible for C11 methoxylation of **5**. During vindoline biosynthesis in *C. roseus*, tabersonine 16-hydroxylases (T16Hs) of the CYP71D family catalyze C16 hydroxylation of tabersonine to form 16-hydroxytabersonine, followed by *O*-methylation catalyzed by 16-hydroxytabersonine 16-*O*-methyltransferase (16OMT) to complete methoxy group installation^36,37,38^. Given that the C16 position of tabersonine is structurally equivalent to the C11 position in reserpine, we searched for T16H and 16OMT homologs in the *R. verticillata* transcriptome. When T16H homologs proved inactive toward **5**, we rescreened P450 candidate genes co-expressed with *Rv*DYR1. Among these candidates, coexpression of *Rv*CYP72A270 with *Rv*CYP71D820 in *N. benthamiana* leaves infiltrated with **4** resulted in a product that co-eluted with synthetic 11,18β-dihydroxy-3-epi-α-yohimbine (**6**) (Fig. 4e and Supplementary Fig. 12). Next, screening of 16OMT homologs in *N. benthamiana* leaves infiltrated with **6** revealed that only *Rv*11OMT could catalyze the formation of rauvomitorine G (**9**) (Fig. 4d and Supplementary Fig. 13). We further validated this activity using purified N-terminal His6-tagged *Rv*11OMT *in vitro* (Fig. 4f). Finally, co-expression of *Rv*CYP71D820, *Rv*CYP72A270, and *Rv*11OMT in *N. benthamiana* leaves treated with **4** successfully yielded **9** (Fig. 4g), confirming the sequential activities of these enzymes in the pathway.

### Branched pathway

During the co-expression of *Rv*CYP72A270 and *Rv*CYP71D820 in *N. benthamiana* infiltrated with **4**, we observed a minor peak sharing the exact mass with **5** but exhibiting a different retention time (Supplementary Fig. 14). MS/MS analysis of the peak indicated hydroxylation on the indole ring (Supplementary Fig. 14). Given *Rv*CYP72A270’s C11 hydroxylation activity on **5**, we proposed that *Rv*CYP72A270 could also oxidize **4** to form the 11-hydroxy-3-epi-α-yohimbine (**7**). This hypothesis was confirmed when we detected **7** in *N. benthamiana* expressing *Rv*CYP72A270 and treated with **4** (Supplementary Fig. 15). Based on *Rv*11OMT’s known ability to methylate the C11 hydroxyl group of **6**, we investigated its activity toward **7**. Co-expression of *Rv*CYP72A270 and *Rv*11OMT in *N. benthamiana* infiltrated with **4** yielded a product peak (**8**) corresponding to the methoxylated derivative (Supplementary Fig. 15). The MS/MS fragmentation pattern of **8** confirmed the presence of a methoxy group on the indole ring (Supplementary Fig. 15). However, without authentic standards for **7** and **8**, their exact structures could not be unambiguously confirmed. Furthermore, whether **8** could serve as a substrate for *Rv*CYP71D820 to generate **9** remain to be determined. These findings suggest a potential branched pathway from **4** to **9**, adding another layer of complexity to the biosynthetic route (Supplementary Fig. 16).

### Remaining biosynthetic steps

According to our biosynthetic hypothesis, the next step involves the *O*-methylation of the C17 hydroxyl group of rauvomitorine G (**9**) to yield reserpic acid methyl ester (**10**) (Fig. 2). Guided by the robust co-expression patterns observed among our newly identified biosynthetic genes (Fig. 4b and 4c), we initially employed co-expression analysis to identify the putative C17 *O*-methyltransferase. We heterologously expressed 30 methyltransferases that showed strong co-expression correlation (r>0.5) with *Rv*11OMT in *N. benthamiana*, using **9** as the substrate. However, none of these candidates exhibited the desired methyltransferase activity. Further exploration of the *R. verticillata* transcriptome for homologs of well-characterized methyltransferases from the MIA biosynthetic pathway, including *Cr*16OMT and loganic acid *O*-methyltransferase (LAMT), also proved unsuccessful (Supplementary Fig. 17)^38,39^. No activity was detected toward either **9** or 18β-hydroxy-3-epi-α-yohimbine (**5**), the latter being a potential substrate for C17 *O*-methylation in the biosynthesis of deserpidine (**13**), a demethoxylated reserpine derivative with significant biological activities^40^. This challenging identification might be attributed to the unique nature of the C17 hydroxyl group of **9**. Typical plant *O*-methyltransferases target hydroxyl groups with relatively lower pKa values, such as phenolic hydroxyl or carboxyl groups^41^. However, the C17 hydroxyl group in **9** has a significantly higher pKa, making it less prone to form oxyanions for nucleophilic attack on *S*-adenosylmethionine. This distinctive property suggest that an unusual methyltransferase might be required to facilitate the deprotonation and subsequent methylation of this challenging hydroxyl group.

The final step of reserpine biosynthesis involves the transfer of a 3,4,5-trimethoxybenzoyl (TMB) group to reserpic acid methyl ester (**10**) (Fig. 2). Such acylation steps are typically catalyzed by two enzyme families: BAHD-acyltransferases (BAHD-ATs) and serine carboxypeptidase-like acyltransferases (SCPL-ATs)^42^. Recent studies of leonurine biosynthesis in two *Leonurus* species have demonstrated that 4-hydroxy-3,5-dimethoxybenzoic acid (syringic acid) is first glycosylated by a UDP-glucosyltransferase (UGT) to form the glucose ester of syringic acid, which then undergoes SCPL-catalyzed acylation with 4-guanidinobutanol to yield leonurine (Supplementary Fig. 18)^43^. Given the structural similarity between syringic acid and TMB, we hypothesized that a similar pathway operates in reserpine (**11**) biosynthesis, where a UGT glycosylates 3,4,5-trimethoxybenzoic acid (TMBA) to form 1-*O*-TMB-β-D-glucopyranoside (TMB-Glc), followed by SCPL-mediated acylation of **10** (Supplementary Fig. 18).

To identify the UGT responsible for TMBA glycosylation, we performed transcriptome analysis using the *Leonurus* UGT sequence as a query and identified seven UGT candidate genes (Supplementary Fig. 19). These candidates were cloned from *R. verticillata* cDNA and heterologously expressed in *E. coli* BL21. In vitro assays with purified candidate enzymes, TMBA, and uridine diphosphate glucose (UDP-Glc) revealed that *Rv*UGT1 and *Rv*UGT2 catalyzed TMBA glycosylation, yielding a product identical to the synthetic TMB-Glc standard in retention time, exact mass, and MS/MS fragmentation pattern (Supplementary Fig. 20).

For the final acylation step, we identified 15 SCPL candidate genes from the transcriptome using the *Leonurus* SCPL sequence as a query (Supplementary Fig. 21). These candidates possessed the characteristic N-terminal signal peptides typical of SCPLs, which require precise post-translational processing for activation (Supplementary Fig. 22). Despite coexpressing full-length versions of each SCPL candidate with *Rv*UGT1 in both *S. cerevisiae* and *N. benthamiana*, reserpine (**11**) formation was not detected when supplementing with reserpic acid methyl ester (**10**) and TMBA. This lack of activity likely stems from the complex post-translational modifications required for SCPL maturation, including signal peptide cleavage, protein folding in the endoplasmic reticulum, and potential glycosylation^42,44^. These plant-specific processing events may not be adequately reproduced in heterologous hosts, suggesting the need for optimized expression conditions or alternative expression systems that better mimic the native cellular environment.

### Reconstitution of rauvomitorine G (9)

Having identified all biosynthetic enzymes responsible for the production of reserpine biosynthetic intermediate **9**, we aimed to reconstitute **9** in tobacco. We overexpressed eight genes starting from STR to *Rv*11OMT in tobacco and supplemented tryptamine and secologanin as substrates. LC-MS analysis revealed only trace amounts of **3**, **4, 5** and **9** (Fig. 5a). The production efficiency of **9** was significantly compromised likely by the extensive length of the biosynthetic pathway and low efficiency of certain enzymes, particularly *Rv*YOS, which exhibits low catalytic activity and generates multiple yohimbine isomers that can serve as competing substrates for downstream enzymes. To better validate the entire pathway, we divided it into two segments for separate verification: the first segment from tryptamine and secologanin to **2**, and the second segment from **2** to **9**. When tested separately, both segments showed markedly improved performance. The first segment, comprising *Cr*STR, *Rv*SGD, and *Rv*YOS produced **2** (Supplementary Fig. 5). Similarly, the second segment, containing *Rv*YOO, *Rv*DYR1, *Rv*CYP72A270, *Rv*CYP71D820, and *Rv*11OMT, efficiently converted externally supplied **2** to **9** (Fig. 5b and Supplementary Fig. 23). These results confirmed the functionality of all biosynthetic steps and suggested that pathway optimization strategies, such as protein engineering of *Rv*YOS and balanced expression of pathway genes, could enhance the overall production of **9**.

**Fig. 5:**
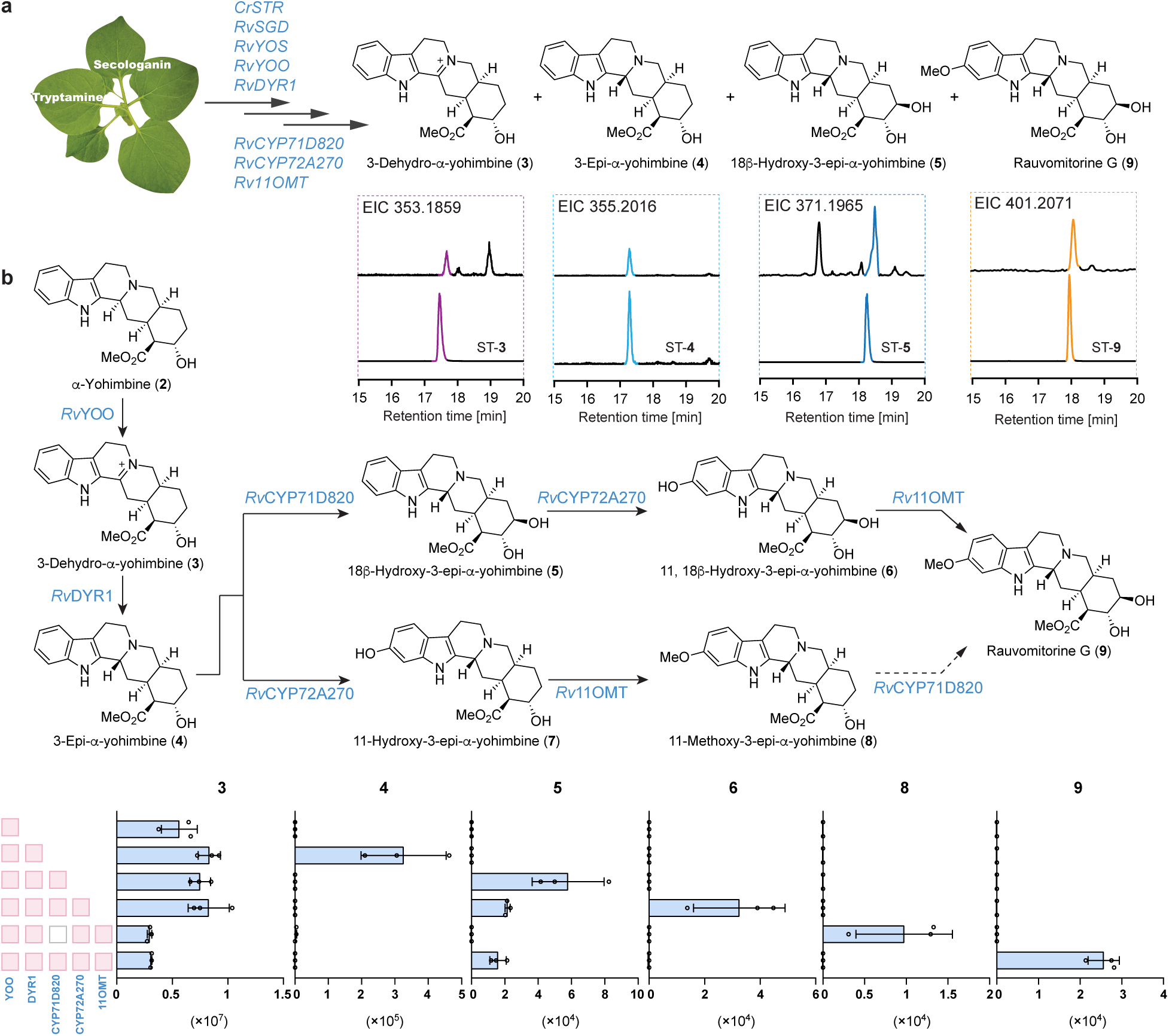
Reconstitution of rauvomitorine G biosynthesis. **a**, Reconstitution of the complete biosynthetic pathway in tobacco using eight enzymes with tryptamine and secologanin as substrates. Extracted ion chromatograms (EIC) show the detection of intermediates **3**, **4**, **5** and final product **9**. **b**, Stepwise reconstitution and validation of the biosynthetic pathway from α-yohimbine (**2**) to rauvomitorine G (**9**), showing the sequential enzymatic transformations and corresponding intermediates. Bar graphs indicate peak areas of pathway intermediates (n = 3 biological replicates, error bars represent standard deviation).

## Conclusion

In conclusion, we have deciphered the biosynthetic pathway of reserpine in *Rauvolfia verticillata*, resolving a long-standing question in natural product chemistry. The molecule’s complex architecture, featuring a unique C3β stereochemistry and five consecutive chiral centers, exemplifies sophisticated natural product biosynthesis. Our investigation revealed the enzymatic basis for C3 epimerization, which proceeds through an oxidation-reduction sequence catalyzed by an FAD-dependent oxidase (*Rv*YOO) and a medium-chain dehydrogenase/reductase (*Rv*DYR1) to establish the crucial C3β stereochemistry. The pathway systematically constructs the five consecutive chiral centers: the first two derive from secologanin, the third and fourth are installed via stereoselective reduction by *Rv*YOS, and the final stereocenter is established through cytochrome P450-catalyzed hydroxylation. Notably, while chemical synthesis typically introduces the methoxy group at an early stage, nature employs a distinct late-stage strategy using a P450-dependent methyltransferase, highlighting the contrasting approaches between biosynthetic and synthetic chemistry. This work not only provides fundamental insights into the biosynthesis of complex alkaloids but also establishes a foundation for metabolic engineering efforts aimed at producing reserpine and related pharmaceutically important compounds. The enzymatic tools uncovered here may enable the development of chemoenzymatic approaches for accessing structurally diverse alkaloids with potential therapeutic applications.

## Methods

### Candidate gene cloning

The full-length sequences of candidate genes were amplified from cDNA using the primers listed in Supplementary Table 2 with 2 × Phanta Max Master Mix according to the manufacturer’s instructions. PCR products were purified and ligated into different linearized expression vectors: 3Ω1 vector (digested with BsaI), and pOPINF and pOPINM vectors (both digested with HindIII-HF and KpnI-HF) using the ClonExpress^®^ Ⅱ one step cloning kit. The ligation reactions were transformed into chemically competent *E. coli* DH5*α* cells (AlpaLifeBio, Shenzhen, China). Transformed cells were plated on LB agar plates containing either 50 μg/mL spectinomycin for 3Ω1 constructs or 50 μg/mL ampicillin for pOPINF/pOPINM constructs. Single colonies were picked and cultured overnight in 3 mL selective LB medium at 37 °C with shaking at 220 rpm. Plasmids were isolated and verified by Sanger sequencing. Confirmed positive strains were mixed with an equal volume of 50% glycerol and stored at −80 °C as frozen stocks.

### Transformation of Agrobacterium tumefaciens

The 3Ω1 plasmids containing the full-length candidate gene were transformed into chemically competent *A. tumefaciens* strain GV3101 (AngYuBio., Shanghai, China) following the manufacturer’s instructions. The transformed cells were plated onto LB agar plates containing 50 μg/mL rifampicin, 50 μg/mL gentamycin, and 25 μg/mL spectinomycin, and incubated at 28 °C for 2-3 days. Single colonies were picked and grown overnight at 28 °C in 5 mL of LB medium containing 50 μg/mL rifampicin, 50 μg/mL gentamycin, and 25 μg/mL spectinomycin. The overnight cultures were mixed with an equal volume of 50% glycerol and stored at −80 °C as frozen stocks.

### Transient expression of candidate genes in *N. benthamiana*

*Agrobacterium* GV3101 strains harboring the constructs of candidate genes were cultivated in 3 mL LB medium supplemented with 25 µg/mL rifampicin, 25 µg/ml gentamycin, and 50 µg/ml spectinomycin overnight at 28 °C with a shaking of 220 rpm. The overnight cultures were centrifuged at 4,000 g for 10 min at room temperature. The cell pellet was resuspended in 3 mL infiltration buffer (50 mM MES, 22.2 mM glucose, 10 mM MgCl_2_ and 100 μM acetosyringone). After centrifugation at 4,000 g for 10 min, the pellet was resuspended in 5 mL infiltration buffer and adjusted to an OD_600_ of 0.6. The bacterial suspension was incubated at room temperature for 2 h in the dark with gentle shaking before being infiltrated into the abaxial (bottom) side of 4-week-old *N. benthamiana* leaves using a needleless 1 mL syringe. Four days post-infiltration, substrate solutions (200 µM in water or in 1 % DMSO) were infiltrated into the same areas of Agrobacterium-infiltrated leaves using a needleless 1 mL syringe. Substrate-infiltrated leaves were harvested 2 days after substrate introduction for metabolite analysis. Each biological replicate consisted of leaves from independent *N. benthamiana* plants (n≥3).

### Heterologous expression of candidate genes in *E. coli* BL21 (DE3)

Chemically competent *E. coli* BL21 (DE3) cells (AlpaLifeBio) were transformed with pOPINF or pOPINM plasmids containing the coding sequences of interest. Transformants were selected on LB agar plates supplemented with 100 μg/mL ampicillin and verified by colony PCR using vector-specific sequencing primers (Supplementary Table 3). For each strain, a single colony was used to inoculate 10 mL of LB medium containing 100 μg/mL ampicillin, which was cultured overnight at 37 °C in a temperature-controlled shaker incubator. The overnight culture was then used to inoculate 400 mL of TB media with 100 μg/mL ampicillin. Cultures were grown at 37 °C with a shaking of 220 rpm until reaching an OD_600_ of 0.6-0.8. At this point, the temperature was lowered to 18 °C, and protein expression was induced by adding isopropyl *β*-D-1-thiogalactopyranoside (IPTG) to a final concentration of 0.5 mM. After approximately 16 hours of induction, the cultures were centrifuged at 8,000 rpm at 4 °C for 10 min to harvest the cells.

### Protein purification

Bacterial cells were resuspended in lysis buffer (100 mM Tris-HCl, 100 mM NaCl, 5% glycerol (*v/v*), pH 8.0) and disrupted using a high-pressure cell disruption equipment (ATS Engineering, Suzhou, China). The remaining steps were performed at 4 °C or on ice. The cell lysates were clarified by centrifugation 12,000 rpm for 30 min, and the supernatant (crude enzyme) was collected. The clarified lysate was loaded onto a column packed with Ni-NTA Beads 6FF agarose resin (Smart-Lifesciences). The column was washed with Tris-HCl buffer before stepwise elution using increasing concentrations of imidazole (30−300 mM) in Tris-HCl buffer. Elution fractions were collected separately and analyzed by sodium dodecyl sulfate polyacrylamide gel electrophoresis (SDS-PAGE) with Coomassie Blue staining. Fractions containing the purified protein were combined and concentrated using Amicon Ultra-15 Centrifugal Filter Units (10 kDa molecular weight cutoff) via centrifugation at 3,500 rpm for 1 h at 4 °C. The concentrated protein solution (<0.5 mL) underwent two rounds of buffer exchange with ice-cold HEPES buffer (50 mM HEPES, pH 7.5, 100 mM NaCl) to remove imidazole. The final protein preparation was diluted to approximately 2 mL with ice-cold HEPES buffer. Protein concentrations were determined using a NanoDrop spectrophotometer. Aliquots of the purified protein were flash-frozen in liquid nitrogen and stored at −80 °C until further use.

### Enzymatic in vitro assays

#### Enzymatic assays for reductase activity

The reductase activity of recombinant *Rv*YOS was assessed using tryptamine and secologanin as substrate. Reaction mixtures (100 µL total volume) contained 1 µM *CrS*TR, 1 µM *Rv*SGD, 2 µM *Rv*YOS, 100 µM NADPH, 25 µM tryptamine, and 25 µM secologanin in 50 mM HEPES (pH 7.4). The reactions were incubated at 30 °C with shaking at 500 rpm for 8 h. The reactions were quenched by adding an equal volume of methanol, followed by vacuum concentration to dryness. The residues were reconstituted in 100 µL methanol and centrifugation at 15,000 × g for 10 min. The supernatants were filtered through 0.22 µm PTFE syringe filters for LC-MS analysis (LC-MS Method 1).

The reductase activity of recombinant *Rv*DYR1 and *Rv*DYR2 was assessed using 3-dehydro-α-yohimbine (**3**) as substrate. Reaction mixtures (50 µL total volume) contained 1 µM purified reductase, 100 µM NADPH, and 40 µM **3** in 50 mM HEPES (pH 7.4). Reactions were incubated at 30°C with shaking at 500 rpm for 12 h. Boiled enzymes (90 °C, 10 min) served as negative controls. Reactions were quenched by adding an equal volume of methanol, followed by centrifugation at 15,000 × g for 10 min. The supernatants were filtered through 0.22 µm PTFE syringe filters for LC-MS analysis (LC-MS Method 2).

#### Enzymatic assays for *O*-methyltransferase activity

The *O*-methyltransferase activity of recombinant *Rv*11OMT candidate was assessed using 50 µM 11,18β-hydroxy-3-epi-α-yohimbine (**6**) as substrate. Reaction mixtures (50 µL total volume) contained 1 µM *Rv*11OMT, 250 µM *S*-adenosyl methionine (SAM), and 200 µM ascorbate in 50 mM HEPES (pH 7.4). Reactions were incubated at 30 °C with shaking at 500 rpm for 12 h and quenched by adding an equal volume of methanol, followed by centrifugation at 15,000 × g for 10 min. Sample preparation for LC-MS analysis was performed as described above for reductase activity assays.

#### Enzymatic assays for UGTs activity

The UGT activity of recombinant *Rv*UGT1 and *Rv*UGT2 was assessed using trimethylgallic acid and UDP-glucose as substrate. Reaction mixtures (100 µL total volume) contained 2 µM purified enzymes, 50 µM trimethylgallic acid, and 25 µM UDP-glucose in 50 mM HEPES (pH 7.4). The reactions were treated as for reductase activity assays, and analysis was performed as described in the LC-MS analysis (LC-MS Method 3).

### Heterologous expression of *Rv*CYP71D820 in *Saccharomyces cerevisiae*

The full-length coding sequence of *Rv*CYP71D820 and *Arabidopsis thaliana* NADPH-cytochrome P450 reductase (*At*CPR) were cloned into the yeast expression vector pESC-Ura under the control of GAL1 and GAL10, respectively. The resulting construct was transformed into *S. cerevisiae* WAT11 chemically competent cells (Coolaber, Beijing, China) using the lithium acetate/single-stranded carrier DNA/polyethylene glycol (LiAc/SS carrier DNA/PEG) method as described previously^45^. Transformants were selected on synthetic defined medium lacking uracil (SD-Ura). Positive clones were identified by colony PCR using vector-specific primers listed in Supplementary Table 4.

Single colonies of recombinant *S. cerevisiae* strains were cultured in 10 ml of SD-Ura liquid medium containing 2% glucose and grown at 30 °C for 24 h with a shaking of 220 rpm. To induce protein expression, the cells were harvested by centrifugation (1,000 rpm, 5 min) at room temperature and resuspended in 500 mL of SD-Ura medium supplemented with 2% glucose and 1.8% galactose. The cultures were then incubated at 30 °C with a shaking of 220 rpm for approximately 3-4 days. Yeast cells were harvested by centrifugation (1,000 g, 5 min) for microsomal protein preparation.

### Preparation of *Rv*CYP71D820-containing microsome

Isolation of yeast microsomal protein followed a previously described protocol^46^. The cell pellet was resuspended in TEK buffer (50 mM Tris-HCl, 1 mM EDTA, 100 mM potassium chloride, pH 7.4) at a ratio of 1 mL buffer per 0.5 g wet cell pellet mass and incubated at room temperature for 5 min. Cells were harvested by centrifugation (1,000 g, 5 min) and then resuspended in 50 mL ice-cold TES B buffer (50 mM Tris-HCl, 1 mM EDTA, 600 mM sorbitol, pH 7.4). The cell suspensions were lysed using an ultra-high-pressure homogenizer (1100 bar, 5 min, 4 °C). Cell debris was removed by centrifugation (10,000 g, 15 min, 4 °C). The supernatant was further centrifuged at 87,000 × g for 60 min at 4 °C. The resulting microsomal pellet was resuspended in 2 mL of ice-cold TEG buffer (50 mM Tris-HCl, 1 mM EDTA, 20% [*v/v*] glycerol, pH 7.4). The protein concentration in the microsomal fraction was measured using the Bio-Rad Protein Assay Kit. Aliquots (100 μL) were snap-frozen in liquid nitrogen and stored at −80 °C until further use.

### *In vitro* assays of *Rv*CYP71D820

The enzymatic activity of *Rv*CYP71D820 was assessed in a reaction mixture (100 μL final volume) containing 40 μM substrate, 2 mM NADPH, 50 mM phosphate buffer (pH 8.0), and 30 μL microsomal fraction. The reaction mixture was incubated at 30 °C for 16 h. Reactions were terminated by the addition of 200 μL methanol. The reaction mixture was concentrated to dryness using a vacuum concentrator (Eppendorf, Hamburg, Germany) at 30 °C. The residue was redissolved in 100 μL methanol and subjected to analysis.

### Metabolite extraction

For *R. serpentina* tissue analysis, freeze-dried tissue was ground to a fine powder, and 10 mg was accurately weighed into a microcentrifuge tube. The tissue was extracted with 300 μL methanol in an ultrasonic bath (SB-3200DT, Scientz, Ningbo, China) at room temperature for 30 min. The extracts were centrifuged at 15,000 rpm for 5 min at room temperature, and the supernatant was filtered through a 0.22 μm PES syringe filter into HPLC vials for analysis.

For *Nicotiana benthamiana* samples, leaves were collected with a 1.0-cm diameter cork borer. The leaf discs were transferred to 2 mL microcentrifuge tubes and lyophilized overnight in a freeze dryer (SCIENTZ-10N, Ningbo Scientz Biotechnology Co., Ltd). The dried tissue was extracted with 600 μL methanol and homogenized using a homogenizer (SWE-D6, Servicebio, Hubei, China). The homogenate was centrifuged at 15,000 rpm for 10 min, and the supernatant was filtered through a 0.22 μm PES syringe filter into HPLC vials for analysis.

### HPLC analysis and purification

Analytical high performance liquid chromatography (HPLC) was performed on an Agilent 1260 Series liquid chromatography system equipped with a multiple wavelength detector and a COSMOSIL 5C18-MS-II column (250 mm × 4.6 mm, 5 μm particle size). The injection volume was 10 μL and the flow rate was 1.0 mL/min using 0.1% formic acid in water as mobile phase A and acetonitrile as mobile phase B. Samples were analyzed with the following liquid chromatography gradient: 0-5 min, 10% B isocratic; 5-25 min, 10% to 50% B; 25-30 min, 50% to 95% B;30-40 min, 10% B isocratic.

Semi-preparative HPLC was carried out using the same Agilent 1260 Series system equipped with either a COSMOSIL 5C18-MS-II column (250×10 mm, 5 μm particle size) or an XBridge BEH C18 OBD Prep column (250 mm ×10 mm, 5 µm particle size, 130Å pore size). The injection volume was 50 μL and the follow rate was 3 mL/min using 0.1% formic acid in water as mobile phase A and acetonitrile as mobile phase B. For compound isolation, the following liquid chromatography gradient was applied: 0-10 min, 10% B isocratic; 10-40 min, 10% to 50% B; 40-50 min, 50% to 95% B; 50-60 min, 10% B isocratic. The fractions were collected based on UV absorption at 254 nm.

### LC-MS analysis

#### Method 1

Liquid chromatography-mass spectrometry (LCMS) analysis was performed using a SIL-30AC HPLC system (SHIMADZU, Kyoto, Japan) coupled to a ZenoTOF 7600 System (Ultra-High-Resolution Quadrupole-Time-of-Flight) mass spectrometer (AB SCIEX, Framingham, MA, USA). Chromatographic separation was achieved using a Phenomenex Kinetex XB-C18 column (100 mm × 2.1 mm, 1.7 µm particle size; 100 Å pore size) maintained at 40 °C. The injection volume was 3 µL and the flow rate was 0.3 mL/min using 0.1 % formic acid in water as mobile phase A and acetonitrile as mobile phase B. Authentic standards were prepared in methanol at concentrations ranging from 20-100 µM. The liquid chromatography gradient elution program was as follows: 0-5 min, 5 % B isocratic; 5-15 min, 5 % B to 50 % B; 15-20 min, 50 % B to 95 % B; 20-22 min, 95 % B isocratic; 22-25 min, 5 % B isocratic. Mass spectrometry was performed in positive electrospray ionization mode using data-independent acquisition (DIA). Operating parameters were as follows: capillary voltage, 3500 V; end plate offset, 500 V; nebulizer pressure, 2.5 bar; drying gas (N_2_) temperature, 250 °C; drying gas flow rate, 11 L/min. Mass spectra were acquired over the range *m/z* 50 to 1000. For MS/MS analysis, fragmentation was triggered at an absolute threshold of 100 with a total cycle time of 0.3-0.5 s and collision energy of 35 eV.

#### Method 2

Additional LC-MS analysis was performed using a Vanquish liquid chromatography system (Thermo Fischer Scientific) connected to an IMPACT Ⅱ System (High Resolution Quadrupole-Time-of-Flight) mass spectrometer (Bruker). Separation was achieved using a Waters Acquity BEH C18 column (100 mm × 2.1 mm, 2.5 µm particle size) operated at 40 °C with a flow rate of 0.3 mL/min and injection volume of 3 µL. The mass spectrometry was operated in positive electrospray ionization mode (ESI^+^) with the following parameters: capillary voltage, 3500 V; end plate offset, 500 V; nebulizer pressure, 2.5 bar; drying gas (N_2_) temperature, 250 °C; drying gas flow rate, 8 L/min. Data acquisition was performed in autoMS/MS mode at 12 Hz over *m/z* 50-1000. Fragmentation was triggered on an absolute threshold of 500 with a cycle time of 0.5 s. Collision energy was applied using a stepping gradient either from 11.0 to 44.0 eV or from 35.0 to 52.5 eV.

#### Method 3

Additional LC-MS analysis was performed using a Vanquish liquid chromatography system (Thermo Fischer Scientific) connected to an IMPACT Ⅱ System (High Resolution Quadrupole-Time-of-Flight) mass spectrometer (Bruker). Separation was achieved using a Waters Acquity BEH C18 column (100 mm × 2.1 mm, 2.5 µm particle size) operated at 40 °C with a flow rate of 0.3 mL/min and injection volume of 3 µL. The mass spectrometry was operated in positive electrospray ionization mode (ESI^-^) with the following parameters: capillary voltage, 4000 V; end plate offset, 500 V; nebulizer pressure, 2.5 bar; drying gas (N_2_) temperature, 250 °C; drying gas flow rate, 8 L/min. Data acquisition was performed in autoMS/MS mode at 12 Hz over *m/z* 50-1000. Fragmentation was triggered on an absolute threshold of 500 with a cycle time of 0.5 s. Collision energy was applied using a stepping gradient from 17.5 to 35 eV.

## Data availability

The gene sequences of all characterized enzymes are provided in Supplementary Table 1. The transcriptome data generated in this study have been deposited in the National Genomics Data Center BioProject databased with the accession number PRJCA035431.

## Acknowledgements

We thank Dr. David Nelson (University of Tennessee) for his assistance with naming *Rv*CYP72A270 and *Rv*CYP71D820 identified in this study. We are grateful to the Shenzhen Synthetic Biology Infrastructure for instrument support and technical assistance. This research was supported by the National Key R&D Program of China (2023YFA0915500) and Shenzhen Science and Technology Program (KJZD20230923115906013) to Y. J, National Natural Science Foundation of China (82104023) to J. C.

## Author contributions

J. C., J. Z, and F. L. performed the experiments and analyzed the data. J. C. and Y. J. designed the experiments, wrote and revised the manuscript. Y. J. supervised the project. All authors discussed the results and commented on the manuscript.

