## Supplementary for "Elucidation of the biosynthetic pathway of reserpine"

### 1. Supplementary Materials and Methods

#### Plant material

*Rauvolfia verticillata* plants were collected from Guigang city, Guangxi province, China, in February 2023. Species authentication was confirmed through DNA barcoding[1]. Tissue samples were collected from half-year-old seedlings, and immediately snap-frozen in liquid nitrogen and stored at −80 °C until further analysis. These samples were subsequently used for both RNA extraction and metabolomic studies.

*Nicotiana benthamiana* plants were grown in an intelligent artificial climate chamber (RXZ-500C-4, Ningbo Jiangnan Instrument Factory, Ningbo, China) using a standard soil mixture (Miracle-Gro, Marysville, OH, US). Growth conditions were maintained at 25 °C with 60% relative humidity under a 16 h light/8 h dark photoperiod. Plants were cultivated 4-6 weeks before *Agrobacterium tumefaciens* GV3101 infiltration. Plants were watered periodically as needed.

#### Chemicals

Yohimbine (**2a**) was purchased from MedChemExpress (Shanghai, China). *α*-Yohimbine (**2**), corynantheine (**2c**), and 3-epi-*α*-yohimbine (**4**) were obtained from BioBioPha (Kunming, China). 1-*O*-(3,4,5-Trimethoxybenzoyl)-*β*-D-glucopyranoside was custom synthesized by WuXi AppTec (Shanghai, China). The following compounds were not commercially available and were synthesized according to procedures detailed in the "Synthetic procedures" section: 3-dehydro-*α*-yohimbine (**3**), 18*β*-hydroxy-3-epi-*α*-yohimbine (**5**), 11,18*β*-hydroxy-3-epi-*α*-yohimbine (**6**), rauvomitorine G (**9**), reserpic acid methyl ester (**10**), and deserpidic acid methyl ester (**12**). The identity and purity (≥95%) of all synthesized compounds were confirmed by nuclear magnetic resonance (NMR) spectroscopy and HPLC analysis. NMR spectra are provided in the "NMR Spectra" section.

All chemical reagents used in reactions were purchased from Aladdin Reagents (Shanghai, China) and Sigma-Aldrich (Darmstadt, Germany) and used without further purification. HPLC-grade solvents were used for HPLC and preparative HPLC analyses, while LC/MS-grade solvents (Merck, Darmstadt, Germany) were used for LC-MS analysis. Deuterated reagents were purchased from Cambridge Isotope Laboratories (Tewksbury, MA, US).

#### Molecular biology kits and reagents

All primers were synthesized by Sangon Biotech (Shanghai, China). Gene and fragment amplifications were performed using an Applied Biosystems ProFlex™ PCR System from Thermo Fisher Scientific (Waltham, MA, US) with one of the following reagents from Vazyme (Nanjing, China): 2 × Phanta Flash Master Mix, 2 × Phanta Max Master Mix, or 2 × Rapid Taq Master Mix. PCR products were purified using the E.Z.N.A.® Gel Extraction Kit from Omega Bio-Tek (Norcross, GA, US). All restriction enzymes were purchased from New England BioLabs (Beverly, MA, US). Gene recombination was carried out using the ClonExpress® II One Step Cloning Kit from Vazyme. Plasmid DNA was isolated using the E.Z.N.A.® Plasmid Mini Kit I from Omega-BioTek. DNA concentrations were measured using a Nanodrop One Spectrophotometer from Thermo Fisher Scientific. DNA sequencing was performed by Sangon Biotech.

**RNA extraction**

Fresh tissues (leaves, stems, and roots) of *R. verticillata* were collected (three biological replicates per tissue type). Total RNA was extracted using the RNeasy Plant Mini Kit (QIAGEN, Hilden, Germany) following the manufacturer's instructions. The quality and quantity of the isolated RNA was assessed using a NanoDrop™ One spectrophotometer, and RNA integrity was verified by 1% agarose gel electrophoresis. First-strand cDNA synthesis was performed using the PrimeScript™ II 1st Strand cDNA Synthesis Kit (TaKaRa, Kusatsu, Japan) with random hexamer primers. The reaction was carried out at 30 °C for 10 min, followed by heat inactivation at 70 °C for 15 min. RNA templates were removed by RNase H treatment at 50 °C for 20 min. The resulting cDNA library was used as the template for candidate gene cloning.

**cDNA library preparation and transcriptome sequencing**

Fresh tissues (leaf, stem, and root) of *R. verticillata* were processed for RNA sequencing at Novogene (https://en.novogene.com), where sequencing library preparation and RNA-seq analysis were performed according to the service provider's standard protocol. RNA integrity was assessed using the Fragment Analyzer 5400 (Agilent Technologies, CA, USA). Sequencing libraries were prepared using the NEBNext® UltraTM RNA Library Prep Kit for Illumina® (NEB, USA) following manufacturer's protocols. Index-coded samples were clustered using the TruSeq PE Cluster Kit v3-cBot-HS (Illumia) on a cBot Cluster Generation System according to the manufacturer's instructions. After cluster generation, the libraries were sequenced on an Illumina Novaseq 6000 platform to generate 150 bp paired-end reads.

**Multiple sequence alignment**

Multiple sequence alignment of amino acid sequences was performed using MUSCLE (Multiple Sequence Comparison by Log-Expectation) algorithm implemented in MEGA 11 software[2]. The aligned sequences were visualized using ESPript 3.0 (https://espript.ibcp.fr)[3].

**Phylogenetic analysis**

The phylogenetic tree was constructed using IQ-TREE (version 2)[4] with the Maximum Likelihood method in a bootstrap test of 1,000 replicates. The resulting tree was visualized using iTOL v6[5].

**NMR analysis**

Nuclear magnetic resonance (NMR) spectra were recorded on a Bruker Avance III HD 400 MHz spectrometer (Bruker Biospin GmbH, Rheinstetten, Germany). Spectrometer control and data processing were performed using Bruker TopSpin version 3.6.1 and MestReNova version 14.0.0 software, respectively. Samples were dissolved in $CD_3OD$, and chemical shifts ($\delta$) are reported in parts per million (ppm) downfield from tetramethylsilane using the solvent resonance as an internal standard for $^1H$ ($CD_3OD$ = 4.870 ppm) and $^{13}C$ ($CD_3OD$ = 49.000 ppm). Data are reported as follows:

chemical shift, multiplicity (s = singlet, d = doublet, t = triplet, q = quartet, m = multiplet, dd = doublet of doublet), coupling constants (J) in hertz (Hz), and integration.

**Synthetic procedures**

<u>Preparation of 3-dehydro-*α*-yohimbine (**3**)</u>

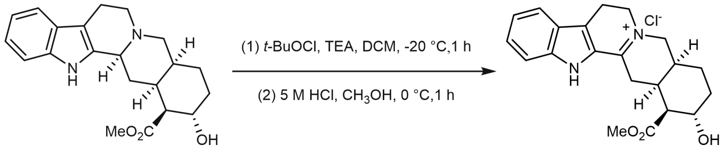

*α*-Yohimbine (**2**) 3-Dehydro-*α*-yohimbine (**3**)

3-Dehydro-*α*-yohimbine (**3**) was synthesized according to a previously reported method[6]. To a stirred solution of *α*-yohimbine (**2**) (35.5 mg, 0.10 mmol) and dry $Et_3N$ (23 μL, 0.165 mmol) in ultra-dry $CH_2Cl_2$ (2 mL) at 0 °C under argon atmosphere was added dropwise *t*-BuOCl (25 μL,0.22 mmol) in ultra-dry $CH_2Cl_2$ (0.2 mL). After stirring for 1 h at 0 °C, the reaction was quenched with water and extracted with $CH_2Cl_2$ (3 × 6 mL). The combined organic layers were evaporated under reduced pressure to afford crude chloroindolenine. The crude product was then treated with 5 M methanolic hydrogen chloride (2.5 mL) at 0 °C under argon atmosphere and stirred for 1 h at the same temperature. The solvent was removed under reduced pressure to obtain crude iminium, which was purified by semi-preparative HPLC using the method described in the "HPLC analysis and purification" section**.** The structure of **3** was confirmed by 1D and 2D-NMR.

**3-Dehydro-*α*-yohimbine** (**3**) : **$^1$H NMR** (400 MHz, $CD_3OD$) *δ* 7.68 (d, *J* = 8.3 Hz, 1H), 7.49 (d, *J* = 8.5 Hz, 1H), 7.42 (dd, *J* = 8.3, 7.2 Hz, 1H), 7.16 (dd, *J* = 8.5, 7.2 Hz, 1H), 4.16 (dd, *J* = 15.7, 3.8 Hz, 1H), 4.01-4.12 (m, 3H), 3.78 (s, 3H), 3.74 (d, *J* = 15.7 Hz, 1H), 3.25-3.32 (m, 4H), 2.62-2.81 (m, 1H), 2.64 (dd, *J* = 10.5, 4.2 Hz, 1H), 2.23-2.29 (m, 1H), 2.06-2.12 (m, 1H), 1.60-1.75 (m, 2H), 1.44 (ddd, *J* = 23.8, 11.8, 5.1 Hz, 1H).

**$^{13}$C NMR** (100 MHz, $CD_3OD$) *δ* 174.6, 166.1, 142.8, 129.7, 127.7, 125.4, 123.9, 122.8, 122.5, 114.2, 66.4, 58.5, 54.3, 54.1, 52.5, 34.7, 34.3, 31.6, 26.5, 24.9, 20.2.

**HRMS** $[M]^+$ calculated for $C_{21}H_{25}N_2O_3^+$: 353.1859, found: 353.1857.

<u>Preparation of 18*β*-hydroxy-3-epi-*α*-yohimbine (**5**)</u>

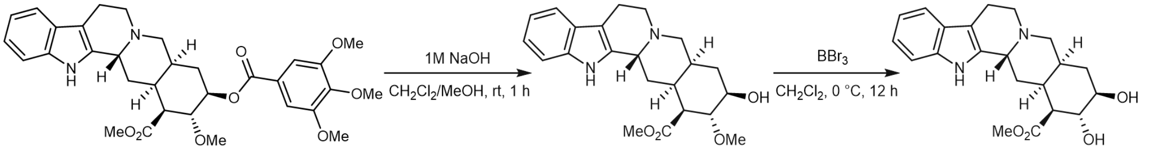

Deserpidine (**13**) Deserpidic acid methy lester (**12**) 18*β*-Hydroxy-3-epi-*α*-yohimbine (**5**)

18*β*-Hydroxy-3-epi-*a*-yohimbine (**5**) was synthesized according to a previously reported method[7]. To a stirred solution of deserpidine (**13**) (50 mg, 0.086 mmol) dissolved in $CH_2Cl_2$/MeOH (*v*/*v* =1:1, 2.0 mL) was added NaOH (2.0 M, 1.0 mL, 2.0 mmol). The mixture was stirred at room temperature for 1

h. The reaction was quenched with HCl (1.0 M), and the solvent was removed under vacuum. The residue was purified by semi-preparative HPLC to yield deserpidic acid methyl ester (**12**) (20 mg). The structure of **12** was confirmed by 1D-NMR and found to be consistent with the literature[7].

To a stirred solution of **12** (20 mg, 0.052 mmol) in dry $CH_2Cl_2$ (1.0 mL) under argon at 0 °C, $BBr_3$ (0.53 mL, 0.52 mmol, 1.0 M solution in $CH_2Cl_2$) was added. After 8 h, the reaction was quenched with saturated solution of $NaHCO_3$ (1.0 mL) and extracted with $CH_2Cl_2$ (3 × 2mL) followed by EtOAc (3 × 2 mL). The combined organic layers were evaporated under reduced pressure to afford the crude product. The crude product was dissolved in MeOH and purified by semi-preparative HPLC to afford **5**. The structure of **5** was confirmed by 1D-NMR and reference[8].

**Deserpidic acid methyl ester** (**12**): **$^1$H NMR** (400 MHz, $CD_3OD$) $\delta$ 7.44 (d, $J$ = 7.7 Hz, 1H ), 7.33 (d, $J$ = 7.8 Hz, 1H ), 7.11 (dd, $J$ = 8.6, 7.4 Hz, 1H ), 7.02 (dd, $J$ = 8.6, 7.4 Hz, 1H ), 5.01 (br s, 1H), 3.81 (s, 3H), 3.57 (m, 1H), 3.54-3.60 (m, 2H), 3.54 (s, 3H), 3.39-3.46 (m, 3H), 3.07-3.15 (m, 1H), 3.03 (d, $J$ = 12.8 Hz, 1H), 2.95 (dd, $J$ = 17.0, 4.4 Hz, 1H), 2.51 (dd, $J$ = 10.5, 4.4 Hz, 1H), 2.34 (ddd, $J$ = 16.1, 13.9, 4.4 Hz, 1H), 2.21 ($J$ = 13.9, 4.4 Hz, 1H), 1.94-1.99 (m, 3H), 1.75-1.79 (m, 1H).
**$^{13}$C NMR** (100 MHz, $CD_3OD$) $\delta$ 173.7, 138.5, 127.8, 127.4, 123.5, 120.8, 119.1, 112.5, 106.9, 82.1, 75.4, 61.5, 56.8, 52.5, 52.4, 52.2, 49.9, 34.0, 33.2, 32.0, 24.3, 16.8.
**HRMS** $[M + H]^+$ calculated for $C_{22}H_{29}N_2O_5^+$: 385.2121, found: 385.2127.

**18*β*-Hydroxy-3-epi-*α*-yohimbine** (**5**): $^1$H NMR (400 MHz, $CD_3OD$) $\delta$ 7.45 (d, $J$ = 7.2 Hz, 1H ), 7.34 (d, $J$ = 7.8 Hz, 1H ), 7.12 (dd, $J$ = 8.6, 7.4 Hz, 1H ), 7.03 (dd, $J$ = 8.6, 7.4 Hz, 1H ), 4.96 (s, 1H), 3.81 (s, 3H), 3.79 (m, 1H), 3.38-3.55 (m, 4H), 2.91-3.16 (m, 3H), 2.53-2.58 (m, 1H), 2.21-2.38 (m, 2H), 2.01 (m ,3H), 1.81 (m, 1H).
**$^{13}$C NMR** (100 MHz, $CD_3OD$) $\delta$ 173.8, 138.4, 128.1, 127.9, 123.3, 120.7, 119.0, 112.5, 107.0, 75.0, 72.1, 56.6, 53.2, 52.5, 52.3, 50.1, 34.3, 32.6, 32.3, 24.5, 16.9.
**HRMS** $[M+H]^+$ calculated for $C_{21}H_{27}N_2O_4^+$: 371.1965, found: 371.1962.

<u>Preparation of reserpic acid methyl ester (**10**), 11, 18*β*-hydroxy-3-epi-*α*-yohimbine (**6**), and rauvomitorine G (**9**)</u>

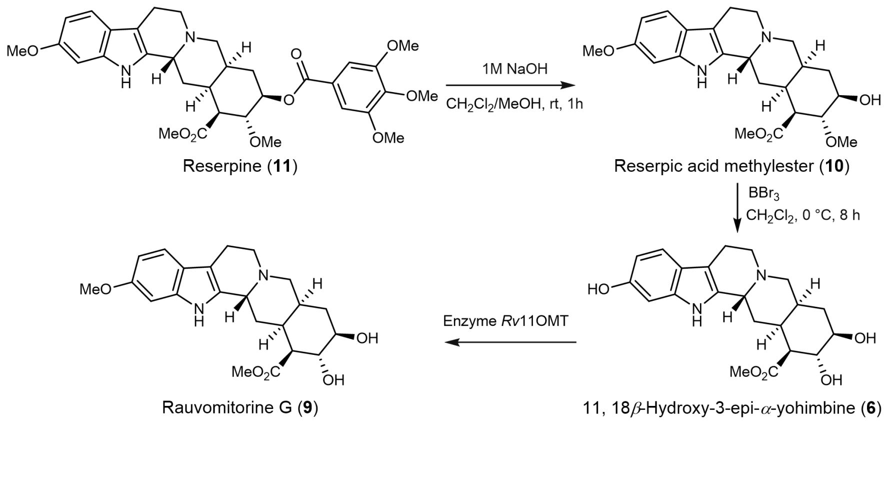

To a stirred solution of reserpine (**11**) (100 mg, 0.164 mmol) in $CH_2Cl_2$/MeOH (*v/v* =1:1, 6.0 mL) was added NaOH (2.0 M, 2.0 mL, 4.0 mmol). The mixture was stirred at room temperature for 1 h. The reaction was quenched with HCl (1.0 M), and the solvent was removed under vacuum. The residue was redissolved in $H_2O$ (5 mL) and extracted with $CH_2Cl_2$ (3 × 5 mL). The organic layers were combined, dried over anhydrous $Na_2SO_4$, filtered, and concentrated *in vacuo* to yield reserpic acid methyl ester (**10**). The structure of **10** was confirmed by 1D and 2D-NMR. Compound **6** was synthesized following the same procedure as for **5**, and its structure was confirmed by 1D and 2D-NMR (Supplementary Table 6). Compound **9** was obtained through enzymatic reaction following the method described in "Enzymatic assays for *O*-methyltransferase activity". The structures of these compounds were confirmed by 1D-NMR and were consistent with literature data[9].

**Reserpic acid methyl ester (10):** $^1$**H NMR** (400 MHz, $CD_3OD$) $\delta$ 7.30 (d, *J* = 8.6 Hz, 1H ), 6.86 (d, *J* = 2.0 Hz, 1H ), 6.69 (dd, *J* = 8.6, 2.0 Hz, 1H ), 3.82 (s, 3H), 3.78 (s, 3H), 3.51 (s, 3H), 3.36-3.52 (m, 4H), 3.02-3.11 (m, 1H), 2.96 (d, *J* = 12.5 Hz, 1H), 2.86 (dd, *J* = 16.6, 3.2 Hz, 1H), 2.52 (dd, *J* = 10.2, 4.2 Hz, 1H), 2.27-2.34 (m, 1H), 2.16 (m, 1H),1.93-2.03 (m, 2H), 1.75( br d, *J* = 10.5 Hz, 1H).
$^{13}$**C NMR** (100 MHz, $CD_3OD$) $\delta$ 173.7, 158.2, 139.2, 126.6, 122.3, 119.6, 110.7, 107.0, 95.9, 82.2, 75.5, 61.4, 56.7, 56.0, 52.5, 52.4, 52.3, 49.9, 34.2, 33.3, 32.2, 24.5, 17.0.
**HRMS** $[M+H]^+$ calculated for $C_{23}H_{31}N_2O_5^+$: 415.2227, found: 415.2229.

**11, 18*β*-hydroxy-3-epi-*α*-yohimbine (6):** $^1$**H NMR** (400 MHz, $CD_3OD$) $\delta$ 7.25 (d, *J* = 8.6 Hz, 1H ), 6.75 (d, *J* = 2.0 Hz, 1H ), 6.61 (dd, *J* = 8.6, 2.0 Hz, 1H ), 4.99 (br s, 1H), 3.81 (s, 3H), 3.75 (dd, *J* = 10.5, 9.5 Hz, 1H), 3.49-3.64 ( (m, 3H), 3.39 (m, 1H), 3.11 (d, *J* = 12.8 Hz, 1H), 3.05 (m, 1H), 2.94 (dd, *J* = 16.7, 5.0 Hz, 1H), 2.56 (dd, *J* = 11.2, 4.5 Hz, 1H ), 2.30 (ddd, *J* = 24.5, 14.4, 4.5Hz, 1H ), 2.23 (m, 1H), 2.04-2.11 (m, 2H), 1.92 (dd, *J* = 24.5, 12.8 Hz, 1H ), 1.80-1.86 (m, 1H).
$^{13}$**C NMR**(100 MHz, $CD_3OD$) $\delta$ 173.7, 155.2, 139.7, 125.1, 121.6, 119.6, 111.0, 106.8, 98.0, 74.8, 72.0, 57.1, 53.0, 52.5, 50.0, 34.0, 32.5, 31.9, 24.3, 16.9.
**HRMS** $[M+H]^+$ calculated for $C_{21}H_{27}N_2O_5^+$: 387.1914, found: 387.1912.

**Rauvomitorine G (9):** $^1$**H NMR** (400 MHz, $CD_3OD$) $\delta$ 7.29 (d, *J* = 8.6 Hz, 1H ), 6.85 (br s, 1H ), 6.66 (br d, *J* = 8.6 Hz, 1H ), 5.01 (br s, 1H), 3.80 (s, 3H), 3.75 (s, 3H), 3.71 (m, 1H), 3.50-3.60 (m, 2H), 3.44 (d, *J* = 11.4 Hz, 1H), 3.32 (m, 1H), 3.05-3.08 (2H, m), 2.92 (d, *J* = 16.5, 1H), 2.49 (dd, *J* = 10.4, 3.1 Hz, 1H ), 2.30-2.38 (m, 1H), 2.19 (d, *J* = 10.4, 13.5, 1H), 1.96-2.00 (m, 1H), 1.78 (m, 1H).
$^{13}$**C NMR** (100 MHz, $CD_3OD$) $\delta$ 173.7, 158.4, 139.4, 125.5, 122.1, 119.8, 110.9, 106.8, 95.9, 74.8, 72.0, 56.8, 56.0, 53.0, 52.5, 52.3, 49.8, 34.0, 32.5, 31.9, 24.2, 16.8.
**HRMS** $[M+H]^+$ calculated for $C_{22}H_{29}N_2O_5^+$: 401.2071, found: 401.2070.

### 2. Supplementary Figures

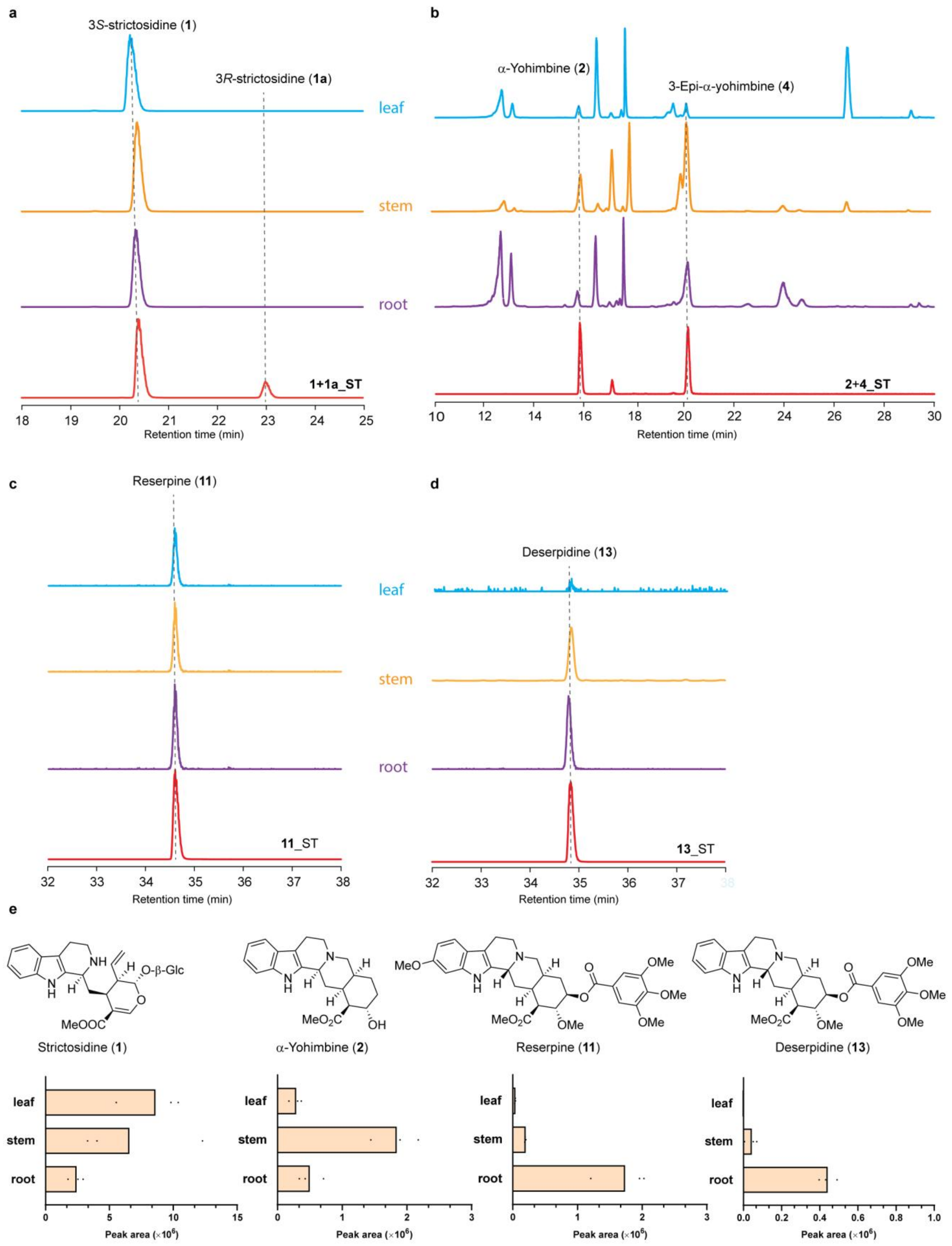

**Supplementary Fig. 1 Distribution of monoterpene indole alkaloids in *R. verticillata*. a-d.** Extracted ion chromatograms (EICs) of metabolites from leaf, stem, and root tissues showing: (**a**)

strictosidine (**1**) and vincoside (**1a**) (*m/z* 531.2337 ± 0.005); (**b**) *α*-yohimbine (**2**) and 3-epi-*α*-yohimbine (**4**) (*m/z* 355.2016 ± 0.005); (**c**) reserpine (**11**) (*m/z* 609.2810± 0.005); and (**d**) deserpidine (**13**) (*m/z* 579.2706 ± 0.005). Standard (ST) compounds are shown in red traces. **e.** Quantitative comparison of metabolite abundance across different tissues, shown as average EIC peak areas (*n* = 3 biological replicates), with corresponding chemical structures shown above. LC-MS analysis was performed using LC-MS Method 2.

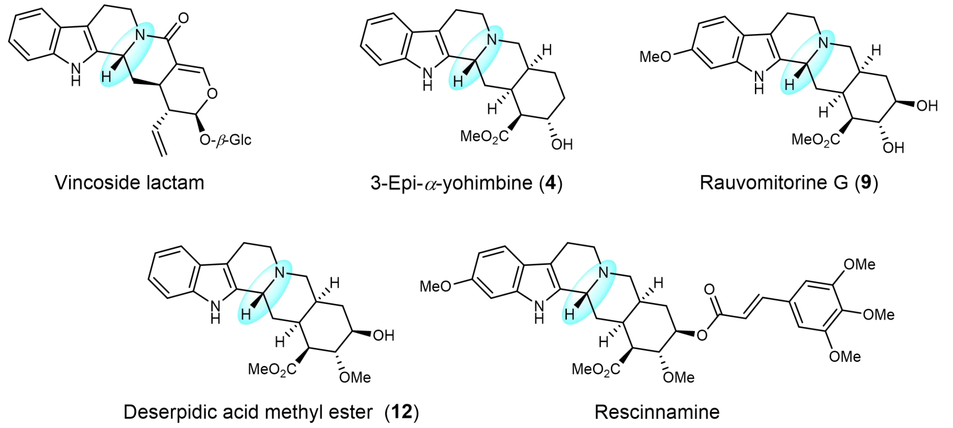

**Supplementary Fig. 2 Chemical structures of C3 *β*-configured intermediates involved in reserpine biosynthesis in *Rauvolfia* species**.

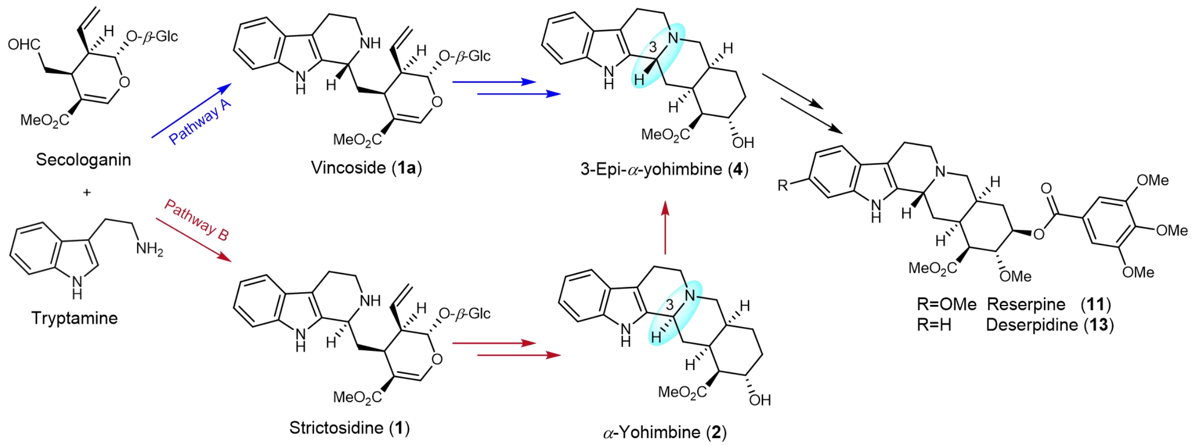

**Supplementary Fig. 3 Two proposed biosynthetic pathways for 3-epi-*α*-yohimbine (4).**

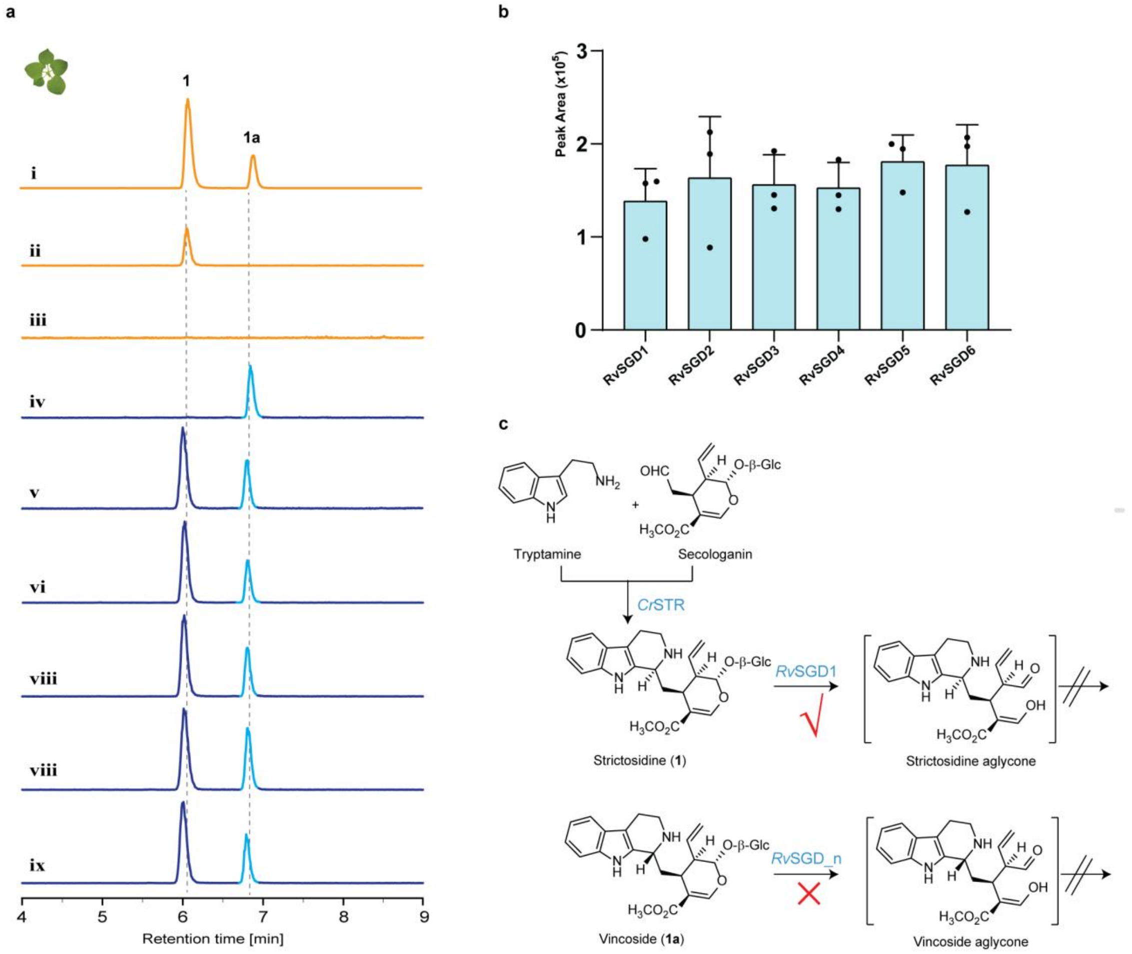

**Supplementary Fig. 4 Functional characterization of *Rv*SGD candidates using strictosidine (1) and vincoside (1a) in *N. benthamiana*.** **a**. Extracted ion chromatograms (EICs) showing *m/z* $[M + H]^+$ = 531.2337 ± 0.005 for: **i**: **1** + **1a** + empty vector; **ii**: tryptamine + secologanin + *Cr*STR; **iii**: tryptamine + secologanin + *Cr*STR + *Rv*SGD1; **iv-ix**: **1** + **1a** + *Rv*SGD1-6, respectively. **b.** Integrated peak area of EICs for the residual substrate **1a** (light blue peak in traces iv-ix, $n$ = 3 biological replicates). **c.** Schematic representation of the pathway showing that STR catalyzes the condensation of tryptamine and secologanin to produce **1**. *Rv*SGD1 exhibits selective deglycosylation activity towards **1** but not **1a**, while *Rv*SGD2-6 show no detectable activity towards either substrate.

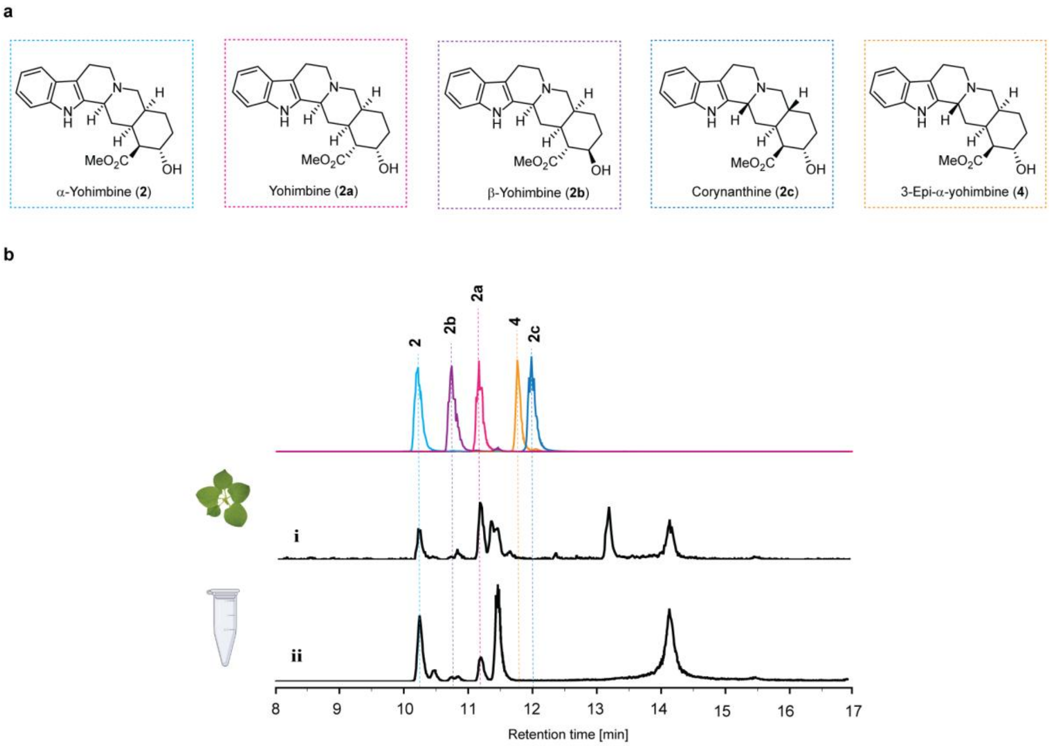

**Supplementary Fig. 5 Characterization of *Rv*YOS in *N. benthamiana* and *in vitro*. a**. Chemical structures of yohimbine stereoisomers. **b**. (**i**): Extracted ion chromatogram (EIC *m/z* $[M + H]^+$ = 355.2016 ± 0.005 ) of multiple yohimbine isomers produced through transient expression of *Cr*STR, *Rv*SGD, and *Rv*YOS in *N. benthamiana*, with co-infiltration of tryptamine and secologanin; (**ii**): Extracted ion chromatogram (EIC *m/z* $[M + H]^+$ = 355.2016 ± 0.005 ) of *in vitro* enzymatic reaction using purified recombinant *Cr*STR, *Rv*SGD, *Rv*YOS enzymes with tryptamine and secologanin as substrates.

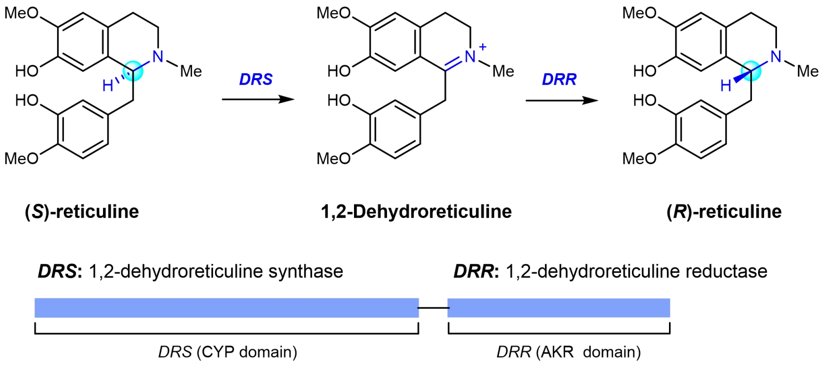

**Supplementary Fig. 6 Enzymatic epimerization of *S*-reticuline to *R*-reticuline.**

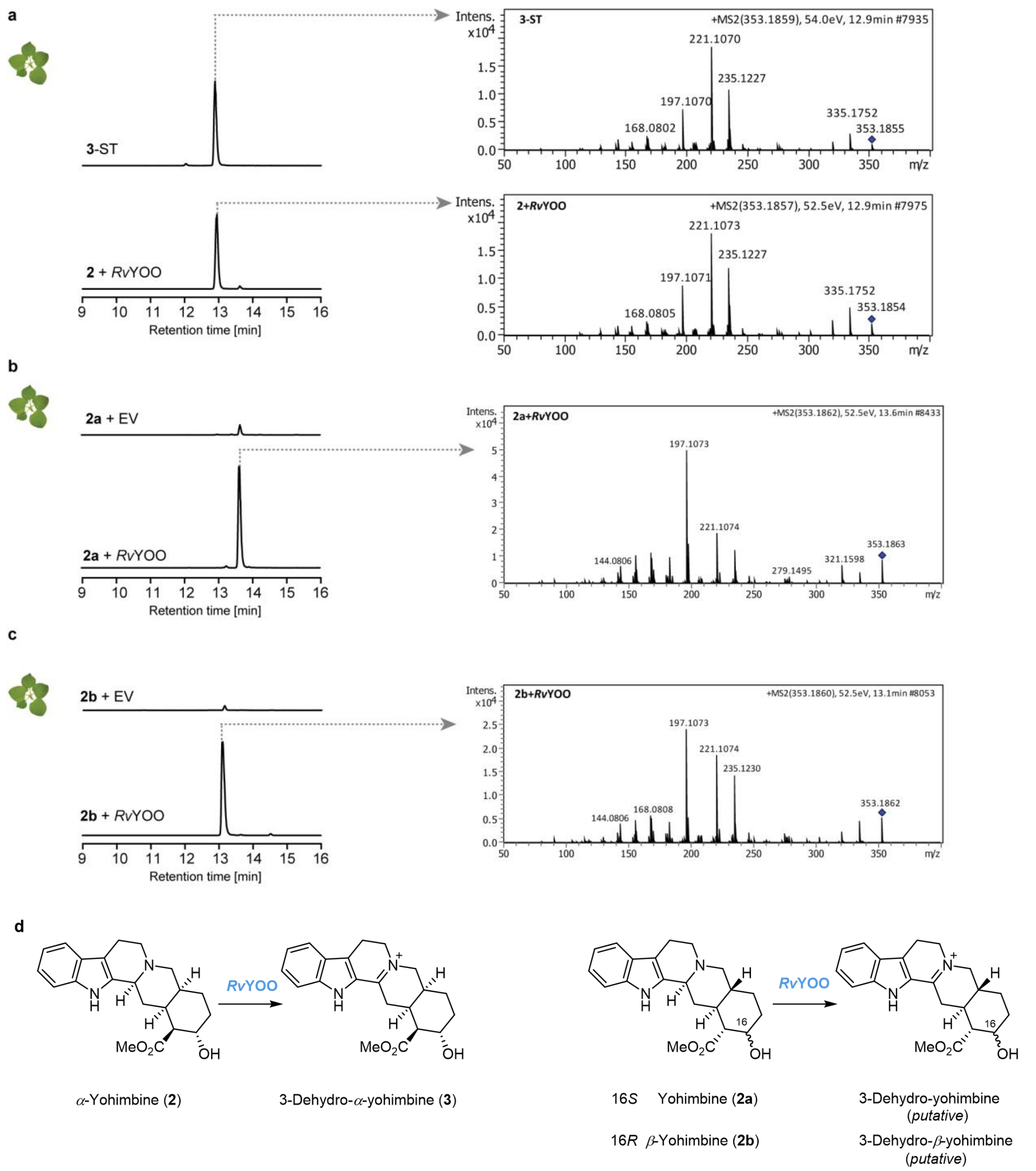

**Supplementary Fig. 7 Functional characterization of *Rv*YOO in *N. benthamiana*.** **a.** LC-MS analysis of *Rv*YOO activity with *α*-yohimbine (**2**): extracted ion chromatogram (EIC) for *m/z* $[M + H]^+ = 353.1859 \pm 0.005$ and comparative MS/MS fragmentation patterns between synthetic standard **3** (**3**-ST, 54 eV) and in planta generated **3** (**2** + *Rv*YOO, 52.5 eV). **b.** LC-MS analysis of *Rv*YOO with yohimbine (**2a**): EIC for *m/z* $[M + H]^+ = 353.1859 \pm 0.005$ and MS/MS fragmentation pattern of planta generated 3-dehydro-yohimbine (**2a** + *Rv*YOO, 52.5 eV). **c.** LC-MS analysis of *Rv*YOO with *β*-yohimbine (**2b**): EIC for *m/z* $[M + H]^+ = 353.1859 \pm 0.005$ and MS/MS fragmentation patterns of planta generated 3-dehydro-*β*-yohimbine (**2b** + *Rv*YOO, 52.5 eV). **d**. Schematic representation of *Rv*YOO-catalyzed oxidation of *α*-yohimbine (**2**), yohimbine (**2a**), and *β*-yohimbine (**2b**) to their corresponding 3-dehydro products. EV: empty vector control.

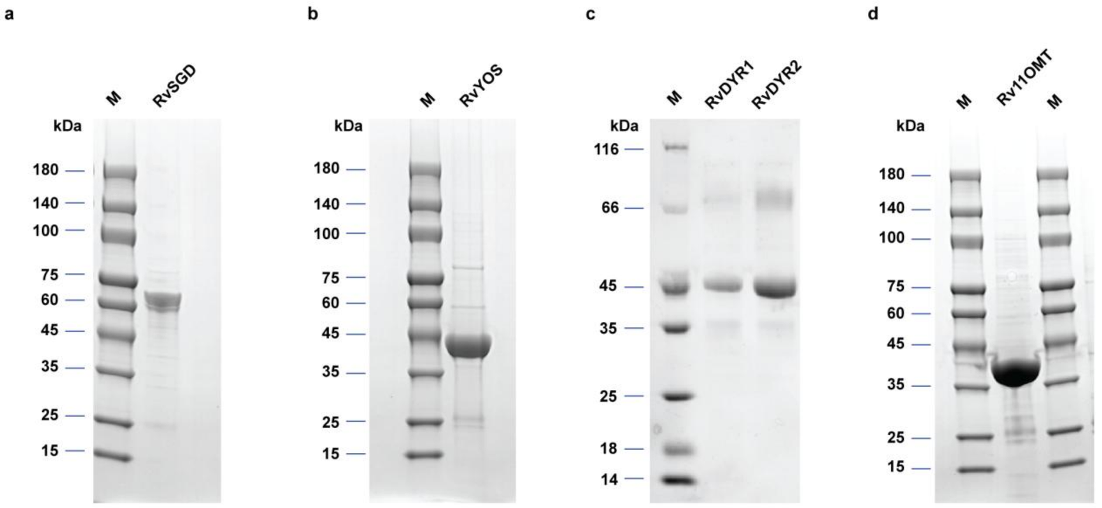

**Supplementary Fig. 8 SDS-PAGE analysis of the purified recombinant proteins. a.** Purified *Rv*SGD protein (theoretical molecular weight: 61.1 kDa). **b.** Purified *Rv*YOS protein (theoretical molecular weight: 38.5 kDa). **c.** Purified *Rv*DYR1 protein (theoretical molecular weight: 40.7 kDa) and *Rv*DYR2 protein (theoretical molecular weight: 40.8 kDa). **d.** Purified *Rv*11OMT protein (theoretical molecular weight: 39.1 kDa). **M**: molecular weight marker.

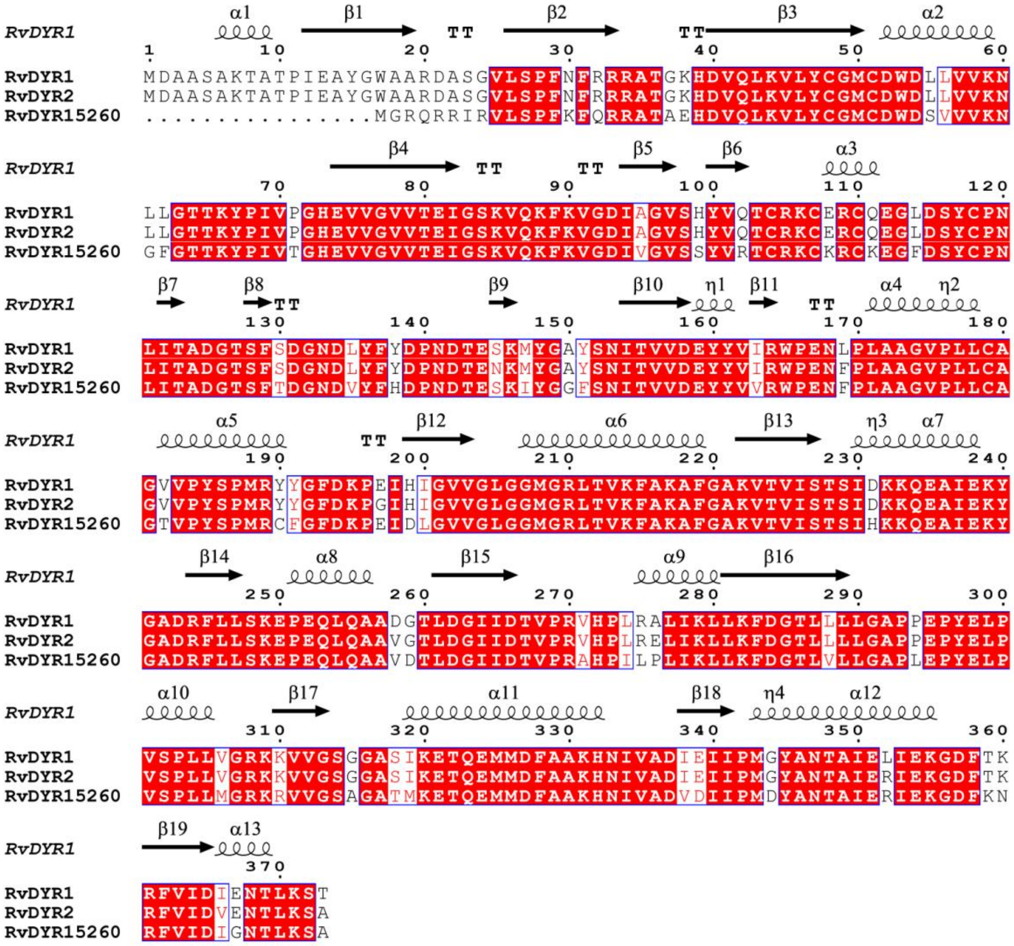

**Supplementary Fig. 9 Amino acid alignment of imine reductases used in this study**. Protein sequence alignment of *Rv*DYR1, *Rv*DYR2, and *Rv*DYR15260.

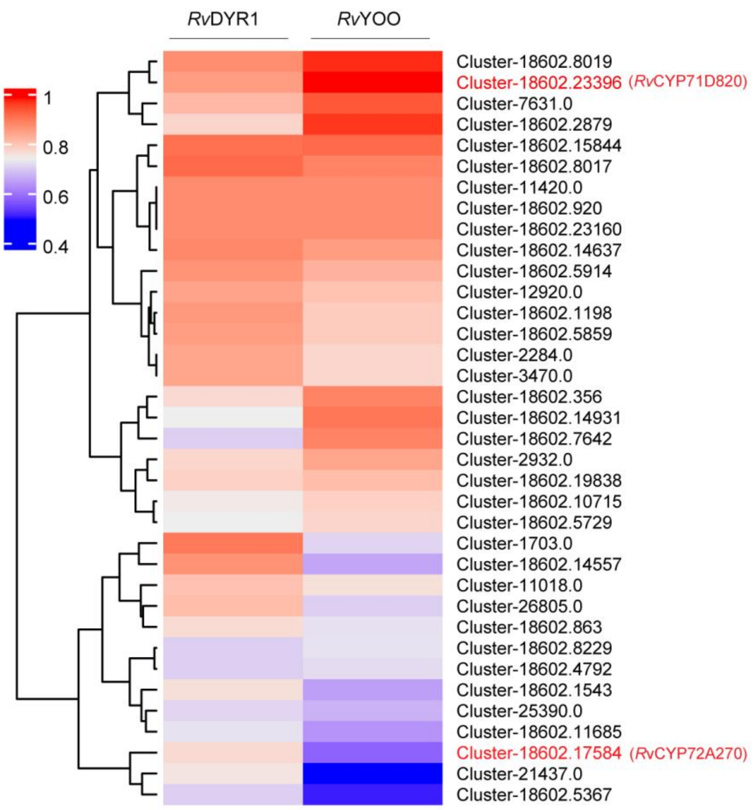

**Supplementary Fig. 10 Coexpression analysis of candidate cytochrome P450 genes with *Rv*DYR1 and *Rv*YOO** (Pearson's r>0.4).

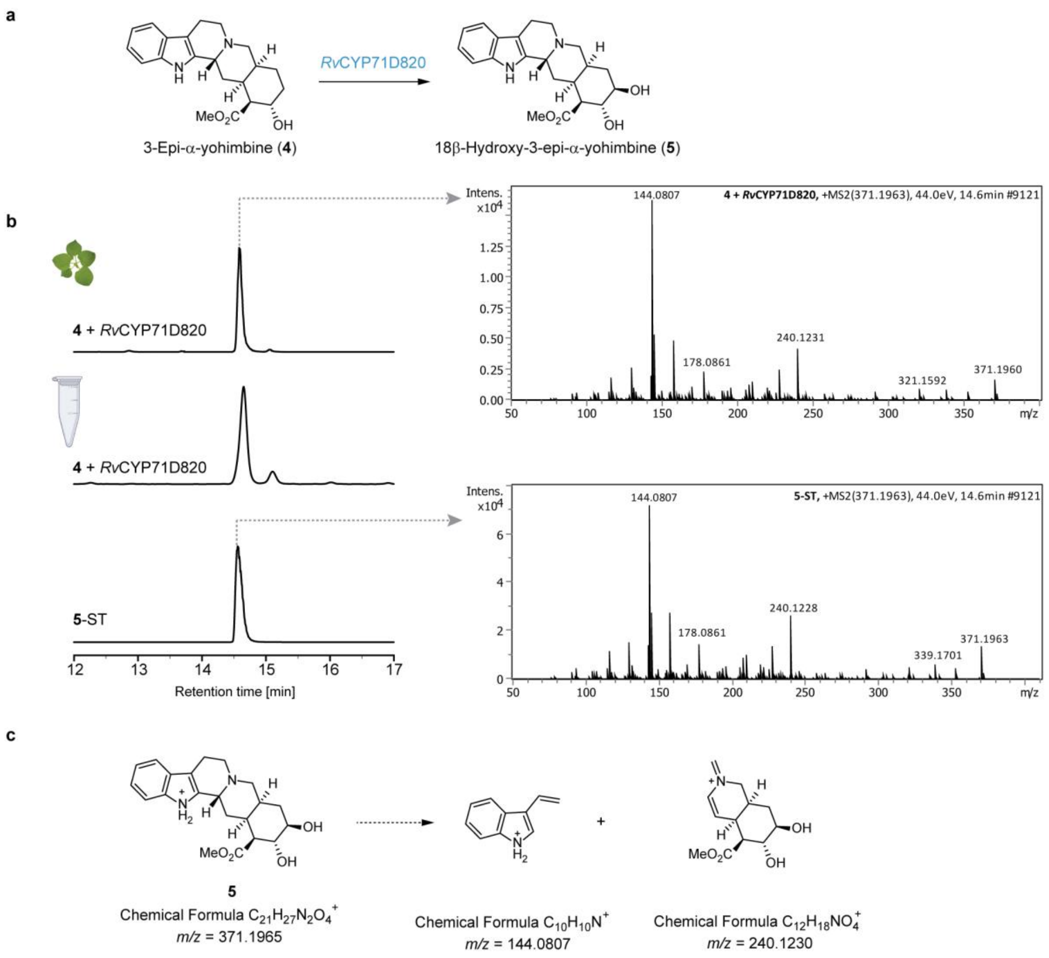

**Supplementary Fig. 11 Functional characterization of *Rv*CYP71D820 in *N. benthamiana* and yeast microsomes. a.** The biosynthetic route from 3-epi-*α*-yohimbine (**4**) to 18*β*-hydroxy-3-epi-*α*-yohimbine (**5**). **b.** Extracted ion chromatograms for **5** (*m/z* $[M + H]^+$ = 371.1965 ± 0.005) generated by transient expression of *Rv*CYP71D820 with co-infiltration of 3-epi-*α*-yohimbine (**4**) in *N. benthamiana*, *Rv*CYP71D820 yeast microsomes *in vitro*, and standard-**5**. MS/MS fragmentation patterns are shown for both in planta generated **5** (4 + *Rv*YOO in *N. benthamiana*, 44.0 eV) and standard **5** (5-ST, 44.0 eV). **c.** Theoretical MS/MS fragmentation patterns of **5**.

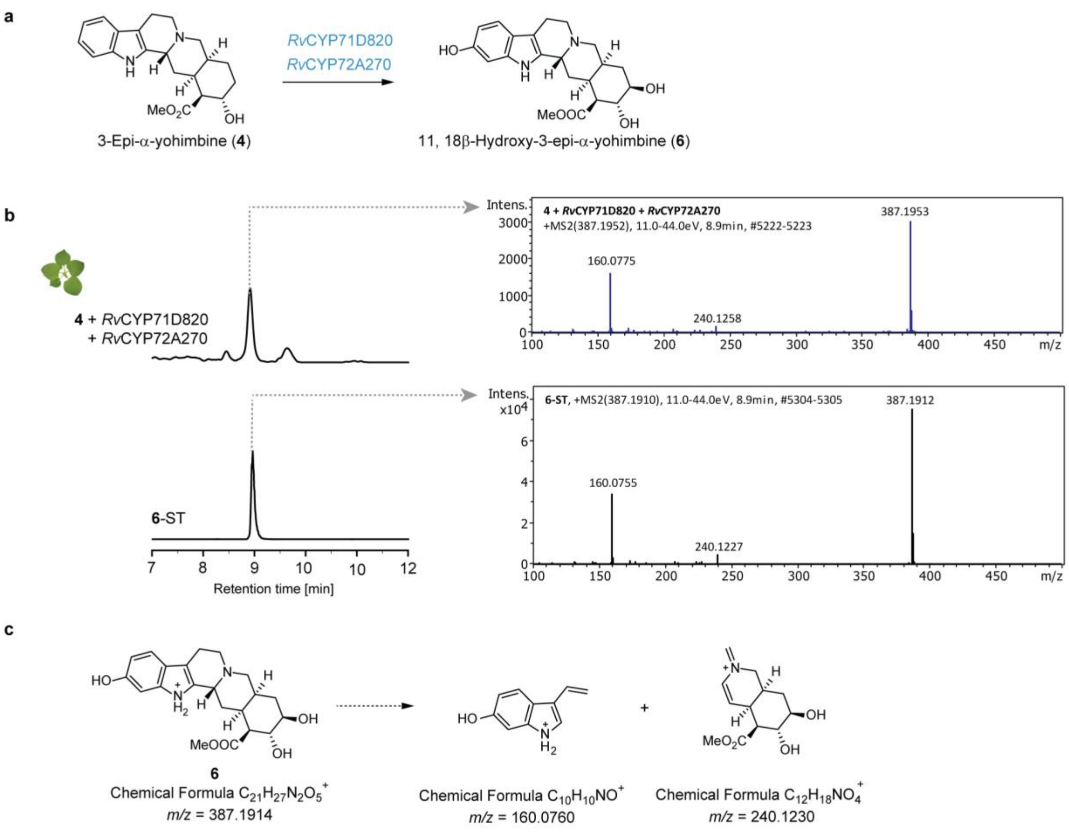

**Supplementary Fig. 12 Functional characterization of *Rv*CYP72A270 in *N. benthamiana.* a.** Biosynthetic pathway from 3-epi-*α*-yohimbine (**4**) to 11, 18*β*-hydroxy-3-epi-*α*-yohimbine (**6**). **b.** Extracted ion chromatograms (EICs) for **6** (*m/z* $[M + H]^+ = 371.19 \pm 0.005$) generated by transient expression of *Rv*CYP71D820 and *Rv*CYP72A270 in *N. benthamiana* with co-infiltration of **4**. MS/MS fragmentation patterns of planta generated **6** (**4** + *Rv*CYP71D820 + *Rv*CYP72A270, 11.0-44.0 eV) compared to synthetic standard **6** (**6-**ST, 11.0-44.0 eV). **c.** Theoretical MS/MS fragmentation patterns of **6**. This experiment was repeated three times with consistent results.

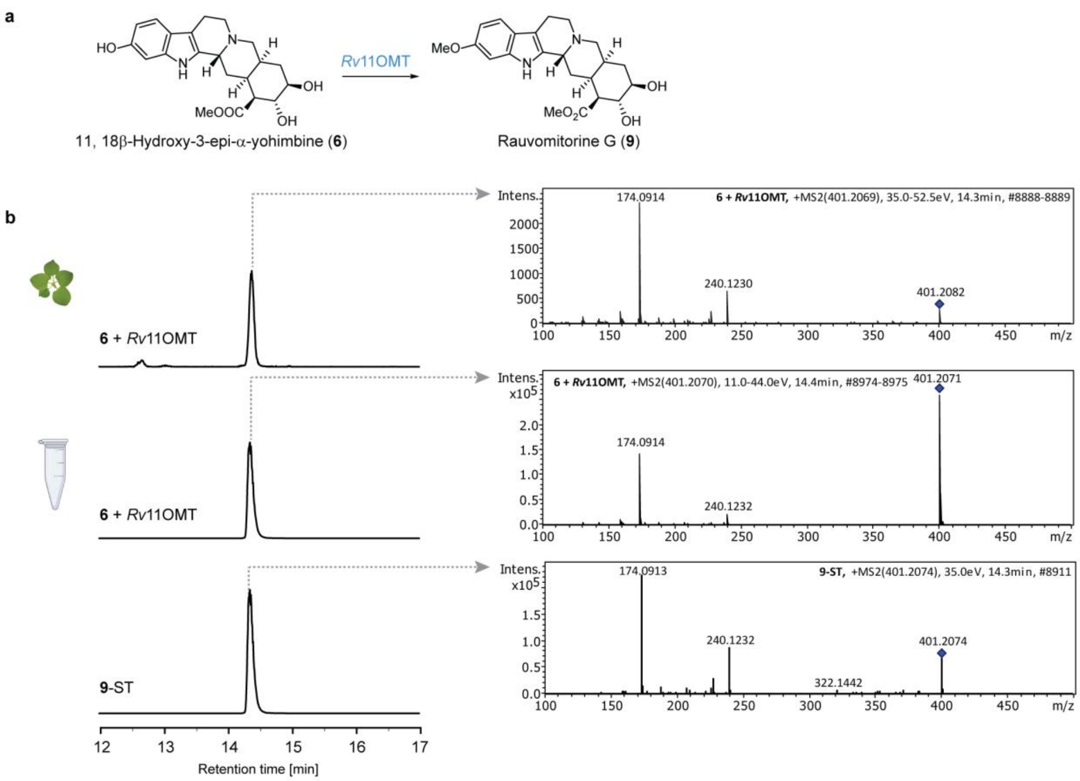

**Supplementary Fig. 13 Characterization of *Rv*11OMT in *N. benthamiana* and *in vitro*. a.** Biosynthetic pathway from 11, 18*β*-hydroxy-3-epi-*α*-yohimbine (**6**) to rauvomitorine G (**9**). **b.** Left: Extracted ion chromatograms (EICs) of **9** (*m/z* $[M + H]^+$ = 401.2071 ± 0.005) from *N. benthamiana* expressing *Rv*11OMT and treated with **6** (top), *in vitro* enzymatic assay with **6** (middle), and synthetic standard (**9**-ST, bottom). Right: Corresponding MS/MS fragmentation patterns of **9** produced in planta (**6** + *Rv*11OMT in *N. benthamiana*, 35.0-52.5 eV), *in vitro* (**6** + *Rv*11OMT *in vitro*, 11.0-44.0 eV), and synthetic standard (**9**-ST, 35.0 eV).

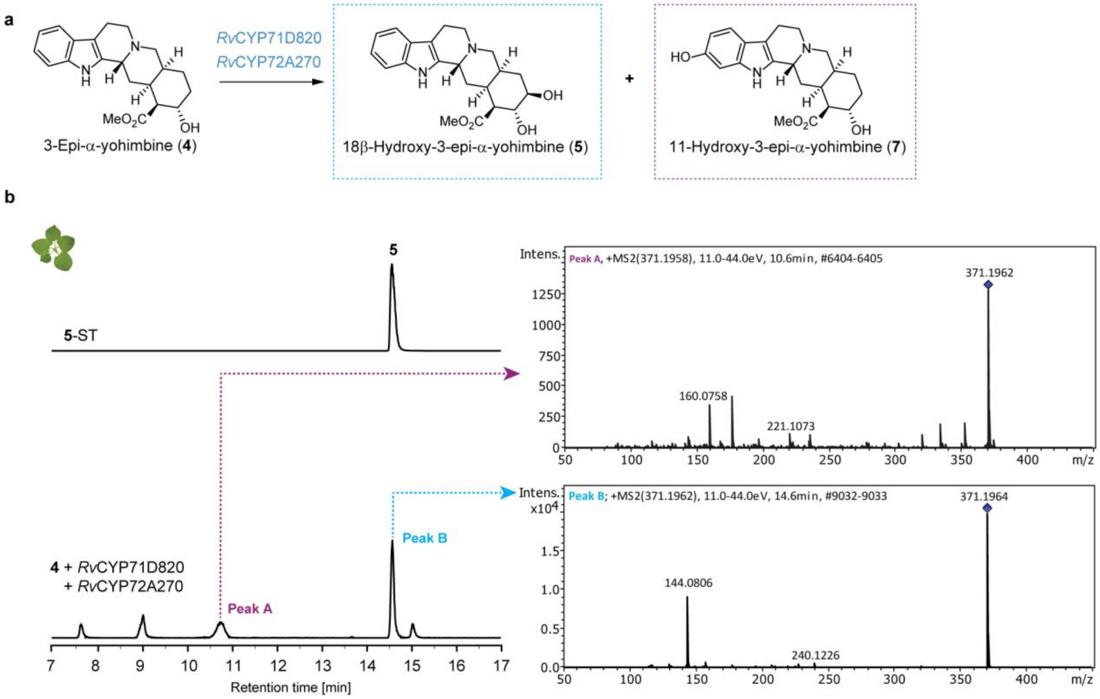

**Supplementary Fig. 14 Functional characterization of *Rv*CYP71D820 and *Rv*CYP72A270 in *N. benthamiana*.** **a.** Schematic representation of the conversion of 3-epi-*α*-yohimbine (**4**) to 18*β*-hydroxy-3-epi-*α*-yohimbine (**5**) and 11-hydroxy-3-epi-*α*-yohimbine (**7**) catalyzed by *Rv*CYP71D820 and *Rv*CYP72A270. **b**. Extracted ion chromatograms (EICs) for *m/z* $[M + H]^+$ = 371.1965 ± 0.005, and MS/MS fragmentation patterns of the products: Peak A, (**7**, 11.0-44.0 eV) and Peak B (11.0-44.0 eV).

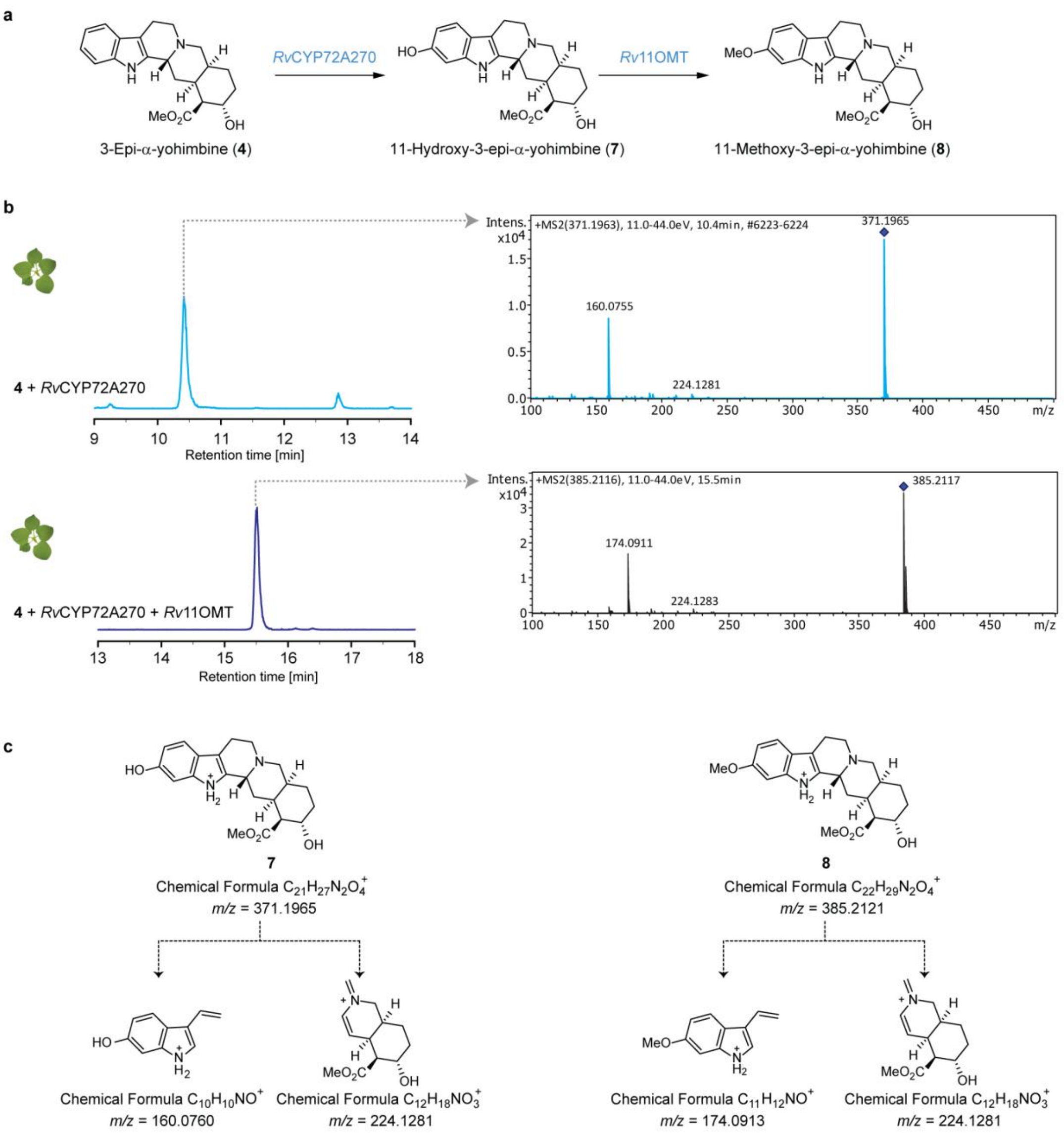

**Supplementary Fig. 15 Identification of 11-hydroxy-3-epi-*α*-yohimbine (7) and 11-methoxy-3-epi-*α*-yohimbine (8) in *N. benthamiana*.** **a**. Biosynthetic pathway for the conversion of **4** to **7** by *Rv*CYP72A270, and subsequent conversion to **8** by *Rv*11OMT. **b.** Extracted ion chromatograms for *m/z* ($[M + H]^+$ = 371.1965 ± 0.005) and MS/MS fragmentation patterns of generated **7** (**4** + *Rv*CYP72A270, 11.0-44.0 eV), extracted ion chromatograms for *m/z* ($[M + H]^+$ = 385.2121 ± 0.005) and MS/MS fragmentation patterns of generated **8** (**4** + *Rv*CYP72A270 + *Rv*OMT, 11.0-44.0 eV). **c**. Theoretical MS/MS fragmentation patterns of **7** and **8**.

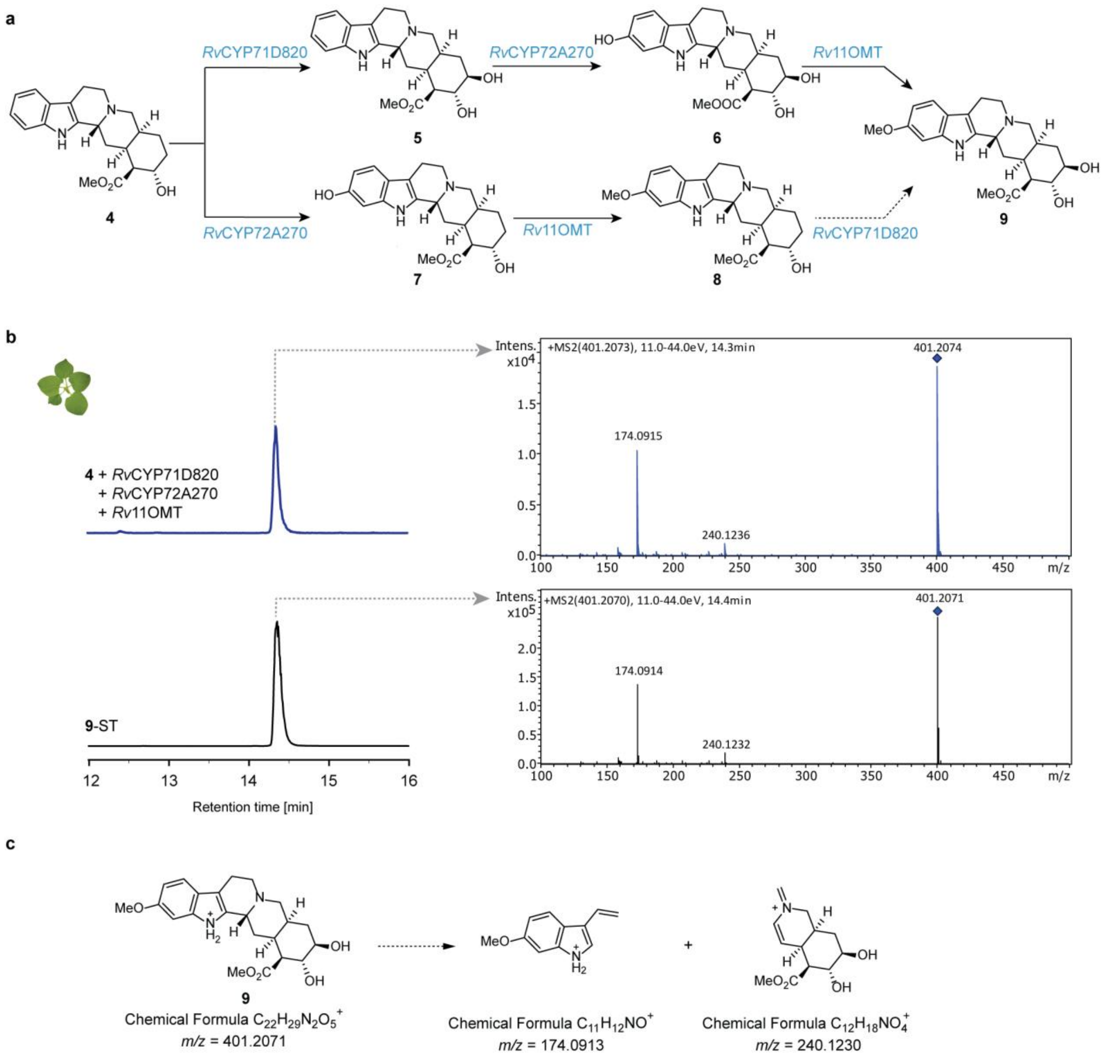

**Supplementary Fig. 16 Identification of rauvomitorine G (9) in *N. benthamiana*. a.** Biosynthetic pathway from **4** to **9** upon transient coexpression of *Rv*CYP71D820, *Rv*CYP72A270, and *Rv*OMT. **b.** Extracted ion chromatograms for *m/z* ($[M + H]^+$ = 401.2071 ± 0.005) and MS/MS fragmentation patterns of generated **9** (**4** + *Rv*CYP71D820 + *Rv*CYP72A270 + *Rv*OMT, 11.0-44.0 eV) compared to synthetic standard **9** (**9**-ST, 11.0-44.0 eV). **c**. Theoretical MS/MS fragmentation patterns of **9**.

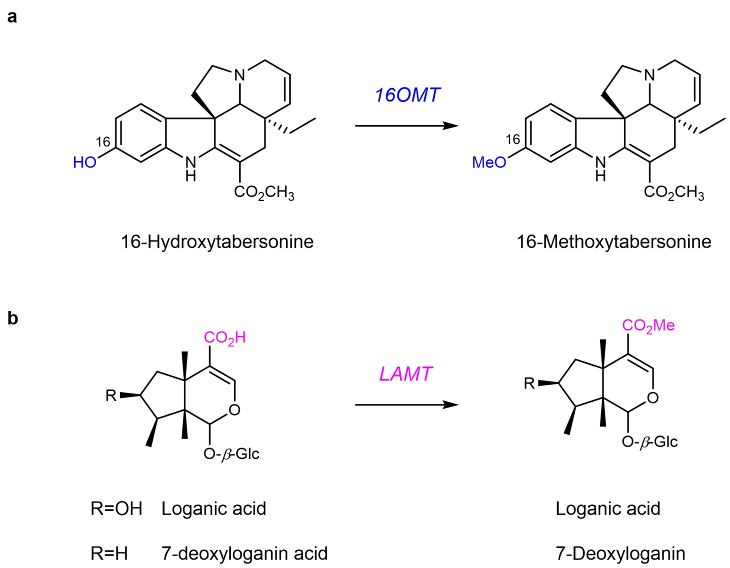

**Supplementary Fig. 17 *O*-methyltransferase-mediated biosynthetic pathways. a**. Conversion of 16-hydroxytabersonine to 16-methoxytabersonine catalyzed by 16-hydroxytabersonine-16-*O*-methyltransferase (16OMT). **b**. *O*-methylation of loganic acid and 7-deoxyloganin to their respective methyl ester products by loganic acid *O*-methyltransferase (LAMT). The -Glc represents the glucose moiety.

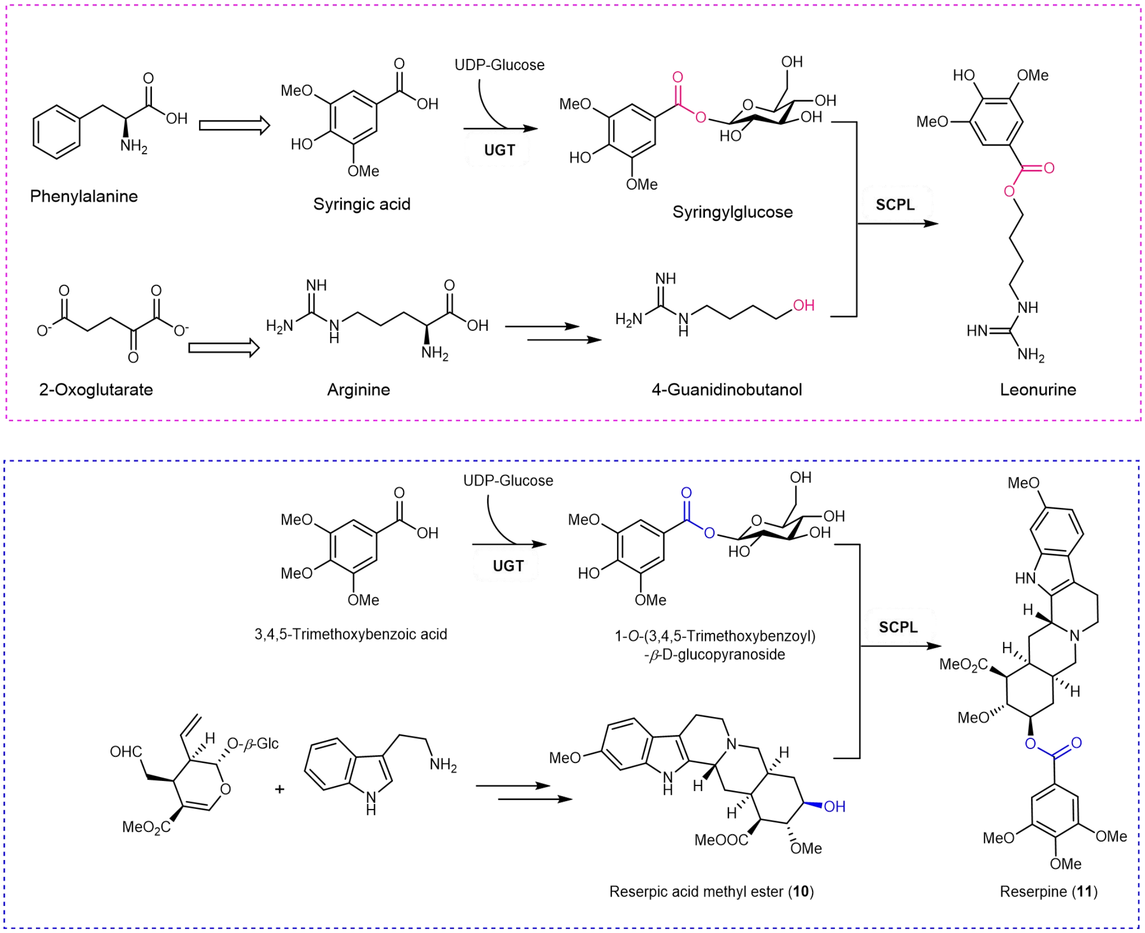

**Supplementary Fig. 18 Parallel biosynthetic pathways for leonurine and reserpine production. a-b.** In both pathways, UDP-glucosyltransferase (UGT) first activates syringic acid or trimethoxybenzoic acid to form their respective glucose esters. These intermediates then undergo SCPL-catalyzed acylation to produce leonurine and reserpine, respectively.

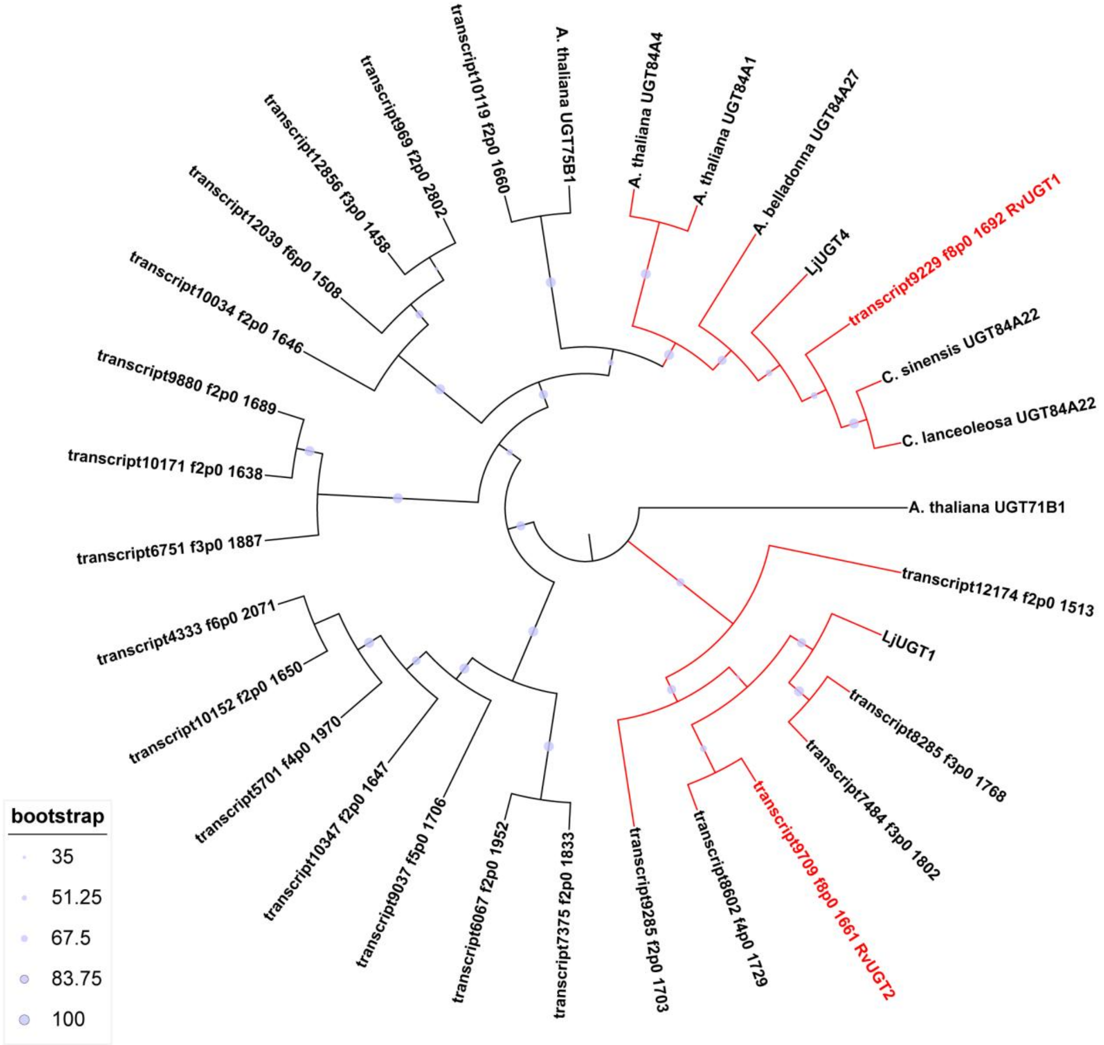

**Supplementary Fig. 19 Phylogenetic analysis of UGTs.** *Lj*UGT1 and *Lj*UGT4 sequences were obtained from the *Leonurus japonicus* genome assembly (GWHBQCV00000000) deposited in the Genome Warehouse in the National Genomics Data Center. Additional reference proteins and their UniProt accession numbers included *Atropa belladonna* UGT84A27 (ATG80135), *Arabidopsis thaliana* UGT71B1 (NP_188812), UGT75B1 (ANM58449), UGT84A1 (Q5XF20), UGT84A4 (O23402), *Camellia lanceoleosa* UGT84A22 (KAI8028241), and *Camellia sinensis* UGT84A22 (ALO19890). Clades highlighted in red indicate tested sequences, with red-labeled UGTs representing experimentally validated active enzymes. Bootstrap values are indicated by circles of varying sizes.

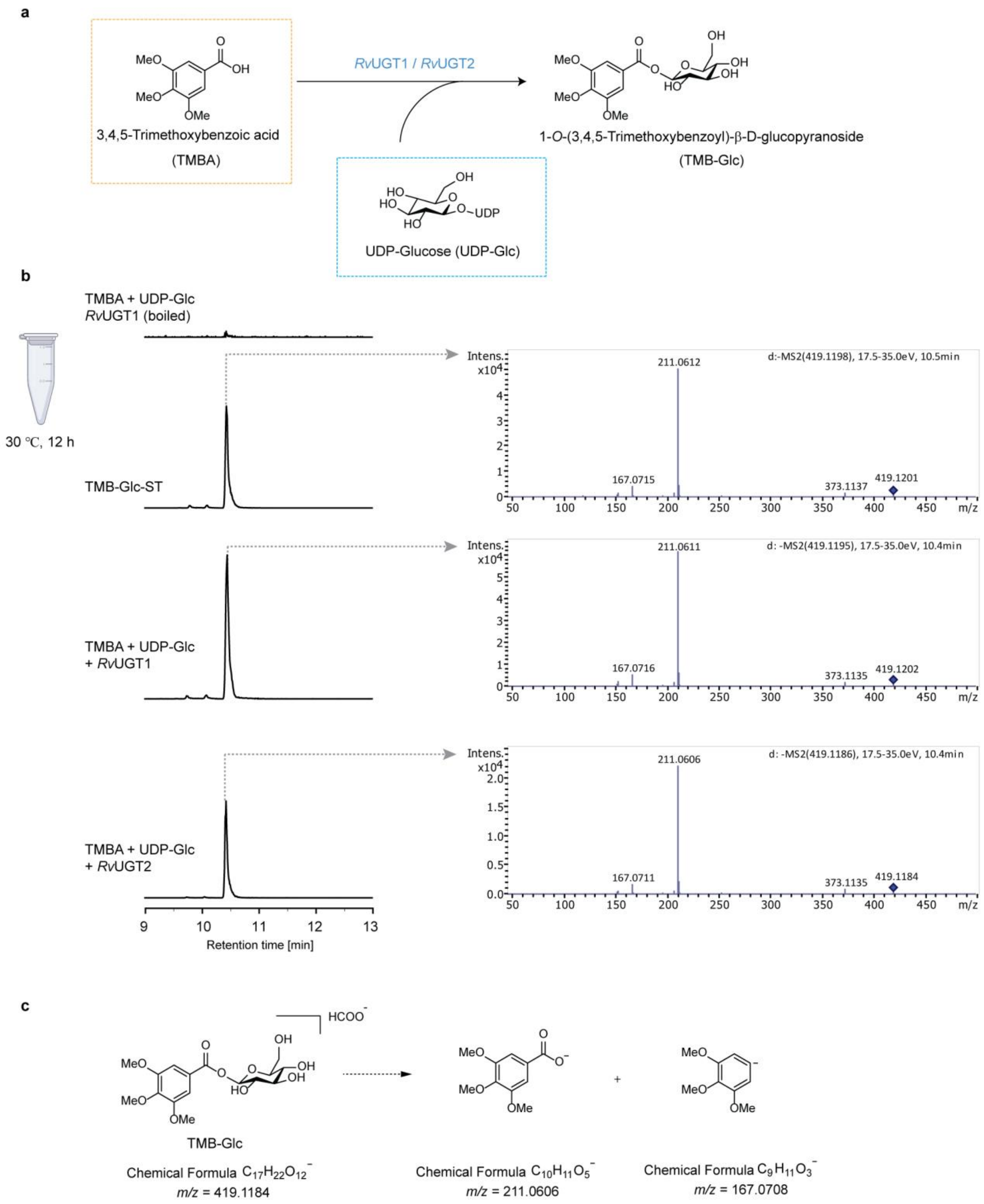

**Supplementary Fig. 20 *In vitro* characterization of *Rv*UGT1 and *Rv*UGT2. a.** Schematic representation of the glycosylation of 3,4,5-trimethoxybenzoic acid (TMBA) catalyzed by *Rv*UGT1/*Rv*UGT2. **b.** LC-MS analysis of enzymatic assays. Extracted ion chromatograms for [M + COO]$^-$ = 419.1184 and MSMS fragmentation patterns of 1-*O*-(3,4,5-Trimethoxybenzoyl)-*β*-D-glucopyranoside (TMB-Glc, 17.5-35.0eV) are shown for: control reactions with boiled *Rv*UGT1 (top), TMB-Glc standard (middle), and reactions with TMBA,

UDP-Glucose (UDP-Glc), and active *Rv*UGT1 or *Rv*UGT2 (bottom). **c.** Theoretical MS/MS fragmentation patterns of TMB-Glc in negative ionization mode.

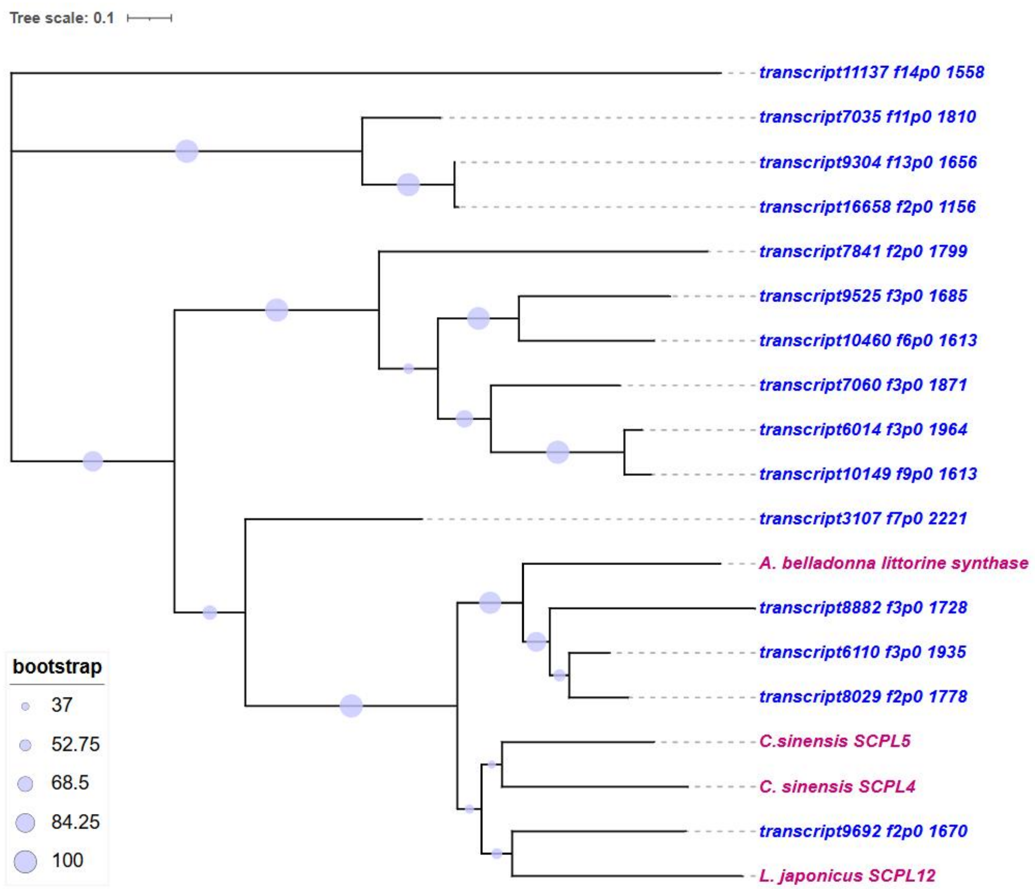

**Supplementary Fig. 21 Phylogenetic analysis of SCPLs.** Candidate genes tested in this study are highlighted in blue. The *Leonurus japonicus* SCPL12 sequence was obtained from the *L. japonicus* genome assembly (GWHBQCV00000000) deposited in the Genome Warehouse in the National Genomics Data Center. Additional reference proteins and their UniProt accession numbers were *Atropa belladonna* littorine synthase (QGI57842), *Camellia sinensis* SCPL4 (UPO25246), and *SCPL5* (UPO25247). Bootstrap values are indicated by circles of varying sizes.

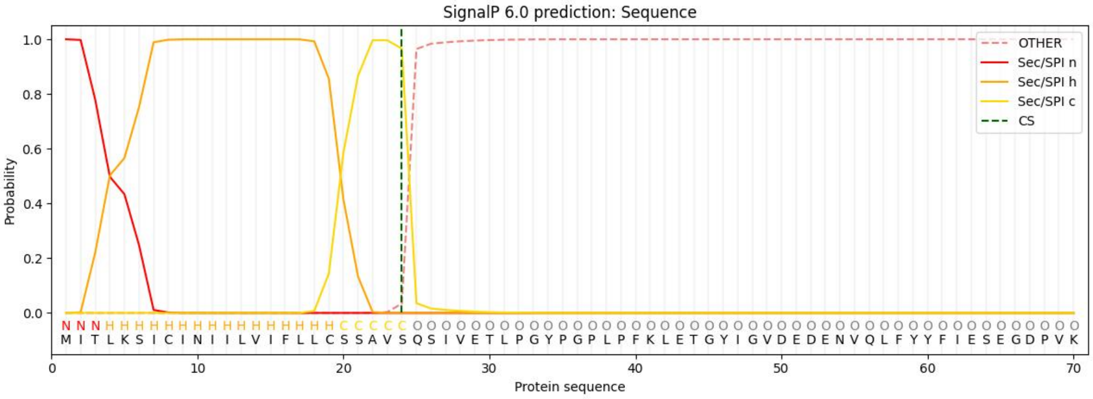

| Protein type | Other | Signal Peptide (Sec/SPI) | Lipoprotein signal peptide (Sec/SPII) | TAT signal peptide (Tat/SPI) | TAT Lipoprotein signal peptide (Tat/SPII) | Pilin-like signal peptide (Sec/SPIII) |
|---|---|---|---|---|---|---|
| **Likelihood** | 0.0003 | 0.9991 | 0.0002 | 0.0001 | 0.0001 | 0.0001 |

**Supplementary Fig. 22 Signal peptide prediction and cleavage site analysis of SCPL proteins.** SignalP 6.0 analysis results for a representative SCPL protein (*Rv*transcript9692/f2p0/1670). The predicted cleavage site is between positions 24 and 25 with a probability of 0.966. The protein exhibits a classical Sec/SPI signal peptide with a likelihood of 0.9991, as determined by the SignalP 6.0 server (https://services.healthtech.dtu.dk/services/SignalP-6.0/).

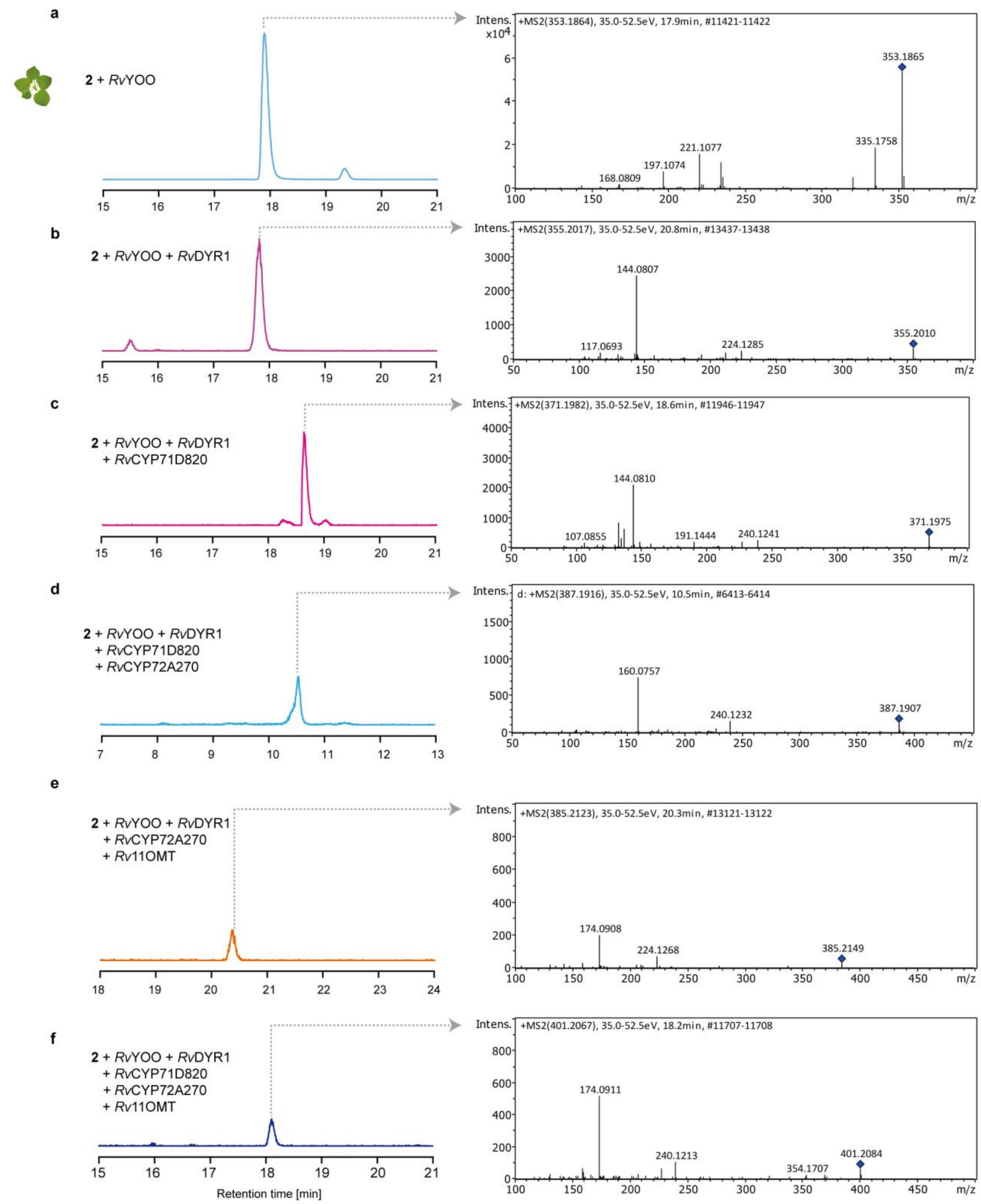

**Supplementary Fig. 23 Reconstitution of rauvomitorine G (9) pathway in *N. benthamiana* from *α*-yohimbine (2). a-f.** Extracted ion chromatograms (EICs) and corresponding mass spectra for the following combinations: (a) *Rv*YOO alone, (b) *Rv*YOO + *Rv*DYR1, (c) *Rv*YOO + *Rv*DYR1 + *Rv*CYP71D820, (d) *Rv*YOO + *Rv*DYR1 + *Rv*CYP71D820 + *Rv*CYP72A270, (e) *Rv*YOO + *Rv*DYR1 + *Rv*CYP72A270 + *Rv*11OMT, (f) *Rv*YOO + *Rv*DYR1 + *Rv*CYP71D820 + *Rv*CYP72A270 + *Rv*11OMT. EICs and MS/MS fragmentation patterns are shown for the following compounds:

3-dehydro-$\alpha$-yohimbine (**3**) (*m/z* $[M+H]^+ = 353.1859 \pm 0.005$), 3-epi-$\alpha$-yohimbine (**4**) (*m/z* $[M + H]^+ = 355.2016 \pm 0.005$), 18$\beta$-hydroxy-epi-$\alpha$-yohimbine (**5**) (*m/z* $[M + H]^+ = 371.1965 \pm 0.005$), 11, 18$\beta$-hydroxy-epi-$\alpha$-yohimbine (**6**) (*m/z* $[M + H]^+ = 387.1914 \pm 0.005$) and rauvomitorine G (**9**) (*m/z* $[M + H]^+ = 401.2071 \pm 0.005$).

### 3. Supplementary Tables

**Supplementary Table 1** Nucleotide sequences for genes cloned and described in this study; start codons are highlighted in bold; stop codons are underlined

| Gene Name | Nucleotide sequence |
|---|---|
| ***Rv*SGD** | **ATG**GAAAGTAATCAAGGTGAGCCTCTTGTTGTTGCCATTGTCCCGAAGCCAAAT<br>GCATCAACAGAGCACAAGAATTCTCACCTCATTCCCGCAACACGAAGTAAGATC<br>GTTGTTCATCGTCGAGATTTCCCCCAAGATTTCATCTTCGGTGCCGGAGGATCTG<br>CTTATCAGTGTGAGGGTGCATATAACGAAGGCAATCGAGGTCCAAGTATATGGG<br>ATACTTTCACTCAGCGAACCCCAGCCAAAATATCTGATGGATCTAATGGAAACC<br>AAGCCATCAATTGTTACCATATGTACAAGGAAGATATCAAAATTATGAAGCAGG<br>CAGGGTTAGAAGCATATAGGTTCTCAATTTCATGGTCAAGAGTATTACCAGGTG<br>GAAGACTAGCTGCTGGAGTGAATAAAGATGGTGTCAAGTTCTATCATGACTTTA<br>TAGATGAGCTTCTAGCCAATGGCATCAAACCTTTTGCAACTCTCTTCCACTGGGA<br>TCTTCCCCAAGCTCTTGAAGACGAATATGGTGGCTTCTTGAGTCACAGAATCGT<br>GGATGATTTTTGTGAGTACGCAGAATTTTGCTTTTGGGAATTTGGTGACAAAATC<br>AAATATTGGACGACATTCAATGAACCGCATACCTTTACTGCAAACGGCTATGCC<br>CTTGGTGAATTTGCACCGGGGAGGGGTAAAGATGGCAAAGGGGATCCAGCCAC<br>AGAACCCTATTTGGTCACACATAATATACTTCTTGCTCACAAAGCTGCTGTGGA<br>AGCATATAGGAACAAATTTCAGAAATGTCAGGAAGGTGAAATTGGAATTGTGCT<br>TAATTCAACGTGGATGGAGCCTCTCAATGATGTCCAGGCTGATATTGATGCTCA<br>TAAAAGAGCTCTTGATTTCATGCTTGGATGGTTTATCGAGCCATTAACAACGGG<br>TGACTACCCAAAATCCATGAGAGAAATCGTAAAAGGACGCCTTCCGAGATTTTC<br>ACCCGAAGATTCCGAAAAATTAAAAGGCTGCTATGATTTTGTCGGAATGAATTA<br>CTATACTGCTACTTATGTGACTAATGCAGCTAAATCCAACTCTGAAAAATTAAG<br>TTACGAAACTGATGATCACGTTGATAAGACTTTTGACCGTGTAGTTGACGGGAA<br>ATCCGTACCAATTGGTGCTGTGTTGTATGGAGAGTGGCAGCATGTCGTTCCTTGG<br>GGACTTTACAAGCTCTTGGTTTACACAAAAGAGACATACCATGTTCCGGTGCTG<br>TATGTCACAGAAAGTGGGATGGTTGAAGAAAACAAAACCAAGATACTGCTTTC<br>AGAAGCTCGTCGCGATCCAGAGAGGACAGATTATCACCAAAAGCATCTTGCAA<br>GTGTACGAGACGCAATCGATGATGGGGTGAATGTAAAAGGTTACTTTGTTTGGT<br>CATTCTTCGACAACTTCGAGTGGAATTTGGGTTTTATATGTCGTTATGGAATTAT<br>TCATGTTGATTACAATAGTTTTGAAAGATGTCCAAAGGAATCAGCCATATGGTA<br>CAAGAATTTCATTGCTGGAGTATCTACTACTTCTCCAGCTAAGAGACGCCGTGA<br>AGAAGCTGAAGGAGTTGAGTTAGTCAAGAGGCAAAAAACCTAA |
| ***Rv*YOS** | **ATG**GCTTCAGAGTCACCGGAAGATGTATATCCAGTGAAGACCGTTGGCTTGGCT |

| | |
|---|---|
| | GCTAAGGATTCATCTGGGGTTTTCTCTCCATTCAAGTTCTCGAGAAGGGCCACA<br>GGTGAACATGATGTGCAGTTCAAAATATTGTATTGTGGGATATGCCAATATGAT<br>AGAGAAATGAGCAGAAACAAATTTGTGATGACCAACTATCCTTATGTTTTTGGG<br>CATGAAATTGTGGGGGTGGTAACTGAGGTTGGACTCAAGGTGCAGAAATTCAA<br>AGTCGGGGACAAAGTTGGCATAGGGAGCATCATTGATACGTGTGGCGAATGCG<br>AAATGTGTACCAATGAAGTGGAAAATTACTGTCCAAAAGCAGTATCAATAGAC<br>AGCAATTTCGGACACAATTACGGAGGTTGTTCAAATATAATGGTGGCCAACGAG<br>AACTTCATCATCCATTGGCCTGAAAACCTTCCTCTGGATTCCGGTGTTCCTCTCC<br>TGTGTGCAGGCATCAATGCTTACAGCCCCATGAGACGTTATGGACTCGATAAAC<br>CAGGGATGCGTGTCGGCATAGCTGGTCTAGGAGGACTTGGACATGTTGCTGTGA<br>GATTTGCCAAGGCATTTGGTGCTAAGGTTACAGTAATCAGTTCATCCCTCAAGA<br>AAAAACATGAAGCCCTTGAGAAATTTGGTGCAGATTCTTTCTTAGTTGGCAGCG<br>ATGCCGAGGAAATGGAGGCTGCAGCCGGAACACTGGATGGCGTCATAGACACT<br>ATACCAGCGGATCACTCTATTGAGCCACTCCTTGCTTTATTGAAGCCTCTTGGGA<br>AGCTTATCATTTTAGGTGCACCAGAAAGGCCGTTTGAGGTGCCCGCTCCGGCCC<br>TACTCCAGGGTGGGAAGTTAATGGCCGCGAGTACCAGTGGAAGCGTGAAGGAA<br>TTACAAGAAATGGTTGATTTTGCAGCAAAACACAACATAGTAGCAGATGTTGAG<br>GTTATCCCCATTGATTATGTGAACGCTGCCATGGAGCGCTTCGATAAATCTGATG<br>TGAAATACCGCTTCGTGATTGACATGGCAAACACTGTCAAATCGGCA<u>TAA</u> |
| ***Rv*YOO** | **ATG**GAAACAAAAGTCCATATGGTTCTTTCATTTGTTATTTCCTTTCTCCTCCTCCT<br>TTCATCTACCCATTCTAGTTCGATTCCTCAAGGCTTCATCGATTGCCTTTCCAATG<br>AATTTTCATCAGACATATATCTACTAGATGTTCTTTATGAGCCCAGCAATTCTTC<br>TTTTGAACCTCTCCTGGAATCTACCATTCAAAATCTTAGATTCTTATCGTCTTCCA<br>CATTAAAGCCACTAGCTATAATCACCCCTATAGCTTTCTCCCATGTCCAAGCTAC<br>AGTAGTCTGCTGCAAACAAAGTGGACTGCAAATTAGAATACGAAGTGGCGGCC<br>ATGATTATGAGGGGCTATCTTACCGATCCCGCCTTCCTTTTGTAATTCTTGACCT<br>CCGAAATCTTACATCAGTAAGCGTGGACGTCAAAGATAACAGCGCTTGGGTCGA<br>GTCTGGAGCGACACTTGGTGATTTGTATTATGGGATAGCTGAGAAAAGTCCTGT<br>GCTTGCCTTCCCTGCTGGGCTCTGCCCAACTGTTGGTGTTGGCGGGCACTTGAGT<br>GGTGGTGGAACTGGAAACTTGGTCAGAAAATACGGACTAGCTGCTGATAATGTC<br>ATCGATGCCCTCATAGTTGTTGCCGACGGTCGAATTCTTGACCGGAAAACTATG<br>GGAACAGATCTGTTTTGGGCCATCAGGGGAGGTGGAGCTGCAAGTTTCGGGGTC<br>GTGGTAGCATGGAAGATCAGGCTCGTACGTGTTCCTCCTACGGTCACTGTTTTCA<br>AATTGAATAAGACTTTAGAGCAAGGAGCTGTAAATCTTCTTCACAAATGGCAGT<br>ATATAGCGCACAAACTCCACGAAGACCTACTCATTACTGCAGAGATTTTCCGGA<br>ATGTTGGGAATAGAACACTTCAAGCGCAGTTCAGCTCGTTCTTCCTCGGTAGAG<br>CTCACCAACTACTGAAAACAATGGAGGAAATCTTCCCTGAACTTGGCCTGAGGA<br>AAGAAGACTGCTCAGAAATGAGTTGGATCGAGTCCGTCGACTATTTTGCAGATT<br>TTCCTAGCAGAGAAACTACAGATAGCCTCAAGAAAAGCATACATTCCCCAGAGT<br>TAACCAAAAACTACTTCAAGTCTAAGTCAGATTTCATTGTGGAACCACTTCCCCT<br>TAGTTCATTACAAAAATTATGGAACCTGTGCCTCGAGGAGGAAAATCTTTCATT |

| | |
|---|---|
| | GCTAATGCATCCTTCTGGCGGGAAAATGGAGATGATAGCGGAATCAGAAACTCC<br>ATTTCCGTTCAGGCAAGGTTTGCTGTATGACATCCAATATGAAGTGGACTGGTA<br>TAGCGAAAATCAATCATCAGAAAAACATATTGACTGGGTAAGAAAGATGTATG<br>ATTACATGACTCCTTACGCATCGAAACGACCAAGGGGTGCTTATCTTAATGCCA<br>GAGATCTTGATTTGGGTACAAATGACAATCCTTTCACAACGTATTCAGAGGCTC<br>AAAGATGGGGGTTCAAATATTTCAAGAAAAACTTTAAGAGATTGGCTACTGTTA<br>AGGGTGTCGTTGATCCAGAAAACTTCTTTTTCTTCGAGCAAAGCATTCCGCCTCT<br>ATGTAACAAGAATAACTTTAAGAGATTGGCTACTGCATAG |
| ***Rv*DYR1** | **ATG**GATGCAGCATCTGCAAAAACAGCAACGCCAATTGAGGCCTACGGATGGGC<br>AGCCAGAGACGCATCTGGAGTTCTCTCCATTCAACTTCCGAAGAAGGGCTAC<br>AGGGAAGCACGATGTGCAGCTCAAAGTGTTGTATTGTGGTATGTGCGATTGGGA<br>TCTACTTGTAGTCAAGAATTTGCTTGGCACTACTAAATATCCCATTGTACCTGGG<br>CATGAGGTGGTGGGTGTGGTGACTGAGATCGGTAGCAAGGTGCAAAAGTTCAA<br>GGTTGGGGACATAGCAGGTGTTAGCCACTACGTTCAGACATGTCGTAAATGTGA<br>GAGATGCCAAGAAGGTCTTGACAGTTATTGTCCAAACTTGATAACAGCAGATGG<br>AACTTCTTTTAGTGACGGAAACGACCTATATTTCTACGATCCAAATGACACAGA<br>GAGCAAGATGTACGGTGCCTATTCCAACATCACGGTTGTCGATGAGTACTACGT<br>AATCCGTTGGCCGGAAAACCTTCCTTTGGCTGCCGGCGTACCTCTTTTATGTGCT<br>GGTGTAGTTCCCTACAGCCCCATGAGATACTACGGATTTGATAAACCCGAAATT<br>CATATTGGTGTCGTTGGACTTGGTGGGATGGGCAGATTAACCGTGAAATTTGCC<br>AAGGCTTTCGGAGCAAAAGTTACAGTAATCAGTACATCCATTGACAAGAAGCA<br>AGAAGCTATTGAGAAATACGGTGCAGATAGATTTTTACTCAGCAAAGAACCTGA<br>GCAGCTGCAGGCCGCGGATGGGACGCTCGATGGCATCATTGACACAGTCCCTAG<br>AGTTCACCCCCTTCGCGCATTGATCAAATTGTTGAAATTCGACGGCACTCTTCTT<br>TTGCTTGGAGCACCCCCGGAGCCATATGAGTTGCCAGTCTCTCCACTGCTCGTAG<br>GTAGGAAGAAGGTGGTTGGAAGTGGTGGTGCGAGTATAAAAGAAACACAAGAG<br>ATGATGGATTTTGCAGCAAAGCACAATATAGTCGCAGATATAGAGATCATTCCA<br>ATGGGTTATGCAAACACTGCAATCGAGCTTATAGAGAAGGGTGATTTCACAAAA<br>CGTTTCGTGATTGATATAGAGAATACATTGAAATCTACTTAG |
| ***Rv*DYR2** | **ATG**GACGCAGCATCTGCAAAAACAGCAACGCCAATTGAGGCCTACGGATGGGC<br>AGCCAGAGACGCATCTGGAGTTCTCTCCATTCAACTTCCGAAGAAGGGCTAC<br>AGGAAAGCACGATGTGCAGCTCAAAGTGTTGTATTGTGGGATGTGCGACTGGGA<br>TCTACTTGTAGTCAAGAATTTGCTTGGCACTACTAAATATCCCATTGTACCTGGG<br>CATGAGGTGGTGGGTGTGGTGACTGAGATCGGTAGCAAGGTGCAAAAGTTCAA<br>GGTTGGGGACATAGCAGGTGTTAGCCACTACGTTCAGACATGTCGTAAATGTGA<br>GAGATGCCAAGAAGGTCTTGACAGTTATTGTCCAAACTTGATAACAGCGGATGG<br>AACTTCTTTTAGTGATGGAAACGATCTATATTTCTACGATCCAAATGACACAGA<br>GAACAAGATGTACGGTGCCTATTCCAACATCACGGTTGTCGATGAGTACTACGT<br>GATCCGTTGGCCGGAAAACTTTCCTTTGGCTGCCGGCGTACCTCTTTTATGTGCT<br>GGTGTAGTTCCCTACAGCCCCATGAGATACTACGGATTTGATAAACCCGGAATT<br>CATATTGGTGTCGTTGGACTTGGTGGGATGGGCAGATTAACCGTGAAATTTGCT |

| | |
|---|---|
| | AAGGCTTTCGGAGCAAAAGTTACAGTAATCAGTACATCCATTGACAAGAAGCA AGAAGCTATTGAGAAATACGGTGCAGATAGATTTTTACTCAGCAAAGAACCTGA GCAGCTGCAGGCGGCGGTTGGGACGCTCGATGGCATCATTGACACAGTCCCTAG AGTTCACCCCCTTCGCGAATTGATCAAATTGTTGAAATTCGACGGCACTCTTCTT TTGCTTGGAGCACCCCCGGAGCCATATGAGTTGCCAGTCTCTCCACTGCTCGTAG GTAGGAAGAAGGTGGTTGGAAGTGGTGGTGCGAGTATAAAAGAAACACAAGAG ATGATGGATTTTGCAGCAAAGCACAATATAGTCGCAGATATAGAGATCATTCCA ATGGGTTATGCAAACACTGCAATCGAGCGGATAGAGAAGGGTGATTTCACAAA ACGATTCGTGATTGATGTAGAGAATACATTGAAATCTGCTTAG |
| ***Rv*CYP71D820** | **ATG**GACCTTCAGCACTTGCCCTTTAACCTGGTTTCTTTCTTCCTCTTCATATTCTT CCTTTCAATAGTTATAAATGGCTGGATAAAGTGCAAAACAAAGACAAATCTGCC ACCAGGGCCATGGAAGCTACCTTTCTTAGGAAACCTTCACAATCTGATGATGGG TTCTCATCCGCATCGTACCCTCAGAGATTTAGCTCGAAAACACGGAGATCTGAT GCTTCTAAAGCTCGGAGAAGTTAATGCCATCGTTGTTTCATCACCTCGTACGGCT AAAGAGGTGCTGAAAACTCATGATCTTGCATTTGCAGACAGGCCAGTTATTCAA GCACCCAAGATCCTCTGTTACGATAATTCAGATGTAGTCTTCAGTCCATACGGTG AGGCCTGGAGACAAATGCGTAAAATATGCGTGTTAGAACTTTTAAGCACGAAGC GTGTTCGATCCTTTCAGCCTATCAGGAAGGATGAAGCTTCTCGTCTGATTGCTTC TCTGGAAGCATCTGCCGGGAAACCGATAGATTTATCGGAGAAAATCAACTGGTA CACAACTTCTATGGTTAGCAGAGCAGCATTTGGTAAGGTAAATAGAGCCGGAAA CACATTCTCAAAGTTGGTGAAGGAAGCATCAGAGGTAGCAGGAGGGTATCACC CAGCTGATATATTCCCTTCCTACAAATTTCTTCATGTTCTTTTCGGTTCGATGTCT AATTTGTTGAAGATTCATGGAAAAACTGACATGATTTTGGAGGAGATGCTCGGT GAGCACATAAAGAACCACCAGATGTCCAAGAAAGCCAATGGTGAGGATGGTAA AGAAGATATAATTGATATCCTACTCAGAATCAAAGATAGCGGTGAGCTGCAATT CCCCTTCACGATCAAGAATGTCAAAGCCATCATTTTTGATATGTTTGCTGCAGGA ACCGAAACTTCATCCTCAACTACGGAATGGGCTATGGCAGAGTTGATGAGAAAT CCACATGTGATGGCCAAGGCACAAGAGGAAGTGAGGCAAGCCTTCAAGGGAAA GAAGATAATTGATGAGGCTGATATTCAAGAATTGAGGTACCTCAAGTTAGTGAT CAAAGAAGTTCTAAGGCTCCATCCTTCTTCCCCTTTGTTGATTCCCAGAGAATGC AGAGAGGCTTGTAAAATAGATGGCTACGATATACCGTTGAAGACGAGGGTTTTC GTCAACATTTGGGCAATTGGAAGAGATGAAAAGTACTGGCCAGATCCGGAGAG TTTCATTCCTGAGAGATTTGAGAACAGTTCCTTTGATTTCACAGGAAATAACTAC GAATATTTACCCTTTGGTAGTGGAAGGAGGATGTGCCCAGGATTGATCTTTGGC TTAGCAAATGTTGAACTTGTTTTAGCTCTGCTACTCTACCATTTCAACTGGAAAC TCCCACCGGGTCTGAGTCCCGGTGACGTAGATATGACGGAGACAATTGGGTTAG GAGCTACAAAACAAAATAGCCTCGTTTTGATTCCCACATTGTATGATCCATCCCT AGCTACATCCTGTTAA |
| ***Rv*CYP72A270** | **ATG**GACAACATCTTCAACTTAATTGCAGTTTCGTGTGTCCTTTCTCTTTTAGTTTG GGCATGGAAGGTACTGAATTCAGCATGGTTAAAGCCAAAAAGGCTAGAGAGAT GTCTCCGAGCGCGAGGCTTTAATGGGAACCCCTATAGACTATTGCTCGGAGACT |

| | |
|---|---|
| | TGAAAGAACTGGGGACGATGCTTAAAGAAGCTCAGTCAAAGCCAATCTGTTTCA<br>CCGATGATATCGTGCCACGGGTTGCACCCTTCTTCCTTGAAACCATCAAGAAAT<br>ATGGTAGCAATTCCTTTTTCTGGTTTGGACCAAACCCGTCGGTTTTCATCTTGGA<br>TCCTGAAGTTGTGAAGGAGATCTTCCTTAACCATAATCGCTTCCAGAAGCCTCCT<br>GCTAATGCAGTTACCAAATTGCTGCAGCGAGGTCTCATCAGCTATGAGGGAGAA<br>GAATGGGAGAAACGCCGGAGAATAATCACCCCATCTTTCCACAGAGAGAAGCT<br>GAAGCATATGCTGCCTGCTTTCCACTTGAGCGTGAGTGAGATGGTAGGGAAATG<br>GGAGGCCACTGTTTCACCAGAAGGTTCAGGCGAGTTAGATGTTTGGCCTTACAT<br>TCGGAGACTAACTAGCGATGCACTTTCTCGCACAGCATTTGGTAGCAGTTATGA<br>AGAAGGGAGAAAGATATTTGAACTTCAAAGTGAACAGGTTGAACTTCTTGTAAT<br>GCTTTCGAGAGCGTTATACCTCCCCGGTTTTAGGTTTCTGCCAACCAAGACGAAC<br>AGGAGGATGAAATATATCCGAAAAGCAGTTGAGGATTCAATTAGACAGATCAT<br>CAACAAAAGATTGAAGGCAATGAAAGAAGGAGAAGCCAGTAAAGATGATTTAT<br>TGGGACTGCTATTGGAATCTAATCAGAAAGAGATTCAAGTACACGGGGACAAG<br>AAATCTGGAATGACCATCCAAGAAATAATTGAAGAATGCAAACTCTTCTATGTT<br>GCAGGGCAGGAAACCACCTCGGTGTTGCTTGTTTGGACATTGATCTTACTCGGT<br>GGGCATCAAGAATGGCAATCGCGTGCCAGAGAAGAAGTTCTGCAACAGTTTGG<br>GAGGAATCAACCAGATTTTGAAGGGCTTAATCATCTAAAAGTTGTGACTAGGAT<br>CTTAAACGAGACTTTAAGACTATATCCGCCCCTAGTCTTTCTCGACCGAACAATT<br>CAAGAAGACACAAAGATAGGAGCACTTTCTCTGCCATCAGGAGTGACGGTCAC<br>ATTACCAGTAATCTTATTGCACCATGATACTAAAATATGGGGTGATGATGCAGT<br>GGAGTTCAAACCAGAGAGGTTCGGTGAAGGAGTGTCCAGTGCAACAAAGGGTC<br>AGGCTGTATTTCTGCCTTTTGGTGGGGGGCCTAGAATATGCATTGGACAGAACT<br>TCACTATGGTAGAAGCAAAACTTGCCTTAGCCATGATTCTGCAGAATTTCTCCTT<br>TGAACTTTCCCCATCATATTCTCATGCTCCGCAGATGATATTCACACTTGTTCCT<br>CAGCATGGGGCTCACCTTATCTTGCACAAATTATAA |
| ***Rv*11OMT** | **ATG**GATTTGCCATCTTCTGAGATCCGTAAAGCTCAAGCTCATTTCTTCAGCCAAG<br>CATTCTCCTTCACAAGCGGTGCGTCTTTAAAATGTGCGGTTCAACTGGGTATTCC<br>AGATGCAATACAAAATCACGGCAAAGCCATGGCTCTCTCTGAGCTCATTGATGC<br>TCTCCCAGTCAGCCCTTCTAAAGCTCCATACATACGCCGCCTAATGCGTATATTA<br>GTCAGTGCCGGCTACTTCTCTGAAGAAAAAACCAATGTTTATGCACTCACTCCTT<br>TGAGCCGTCTTCTTCTGAAGGAGGAGGCACTGAGTTTAAGAGGATTTGTGCTTT<br>CGACCCTCGAAATTGCTGAGATGAAGGCTTGGAATGCCTTGAGCGAGTGGTTTC<br>AGAACGACGATCGAACTGCTTTTGAAACAGCTCACGGAAAAAATTACTGGGAG<br>TATTGCGCCGAAGACAGGTATGGCAAAAGTATCGATGAAGTCATGGCTACTGAC<br>TCGCACTTGGTCTCGAAGCTGCTGATCCCAGAGTACAAGTTCTTGTTCGAGGGCT<br>TGACTTGTTTGGTGGATGTTGGAGGAGGCACAGGGACAATCGCCAAAGCCATCG<br>CAAAAAGTTTCCCTAGCTTAAAGTGCACTGTGTTTGATCTTCCCCATGTGGTGGC<br>CAATCTCGAGCCAACGGAGAACTTGGACTTTGTTGCAGGAGACATGTTTGAGAA<br>AATACCCCCTGCTAATGCGATCCTTCTTAAGTGGGTTCTTCATGACTGGAAAGAC<br>GAAGACTGTGTGAAGATACTCAAAAGCTGCAAGAAGGCAATTCCGGAGAAAGA |

| | |
|---|---|
| | CAAAGGTGGGAAGGTGATCATTATAGACACTGTATTAATGGAGGATAGCCAGA<br>AGCATGAAAATGAATCAGTTAAAACTCAGATATGTAGTGACTTGGACATGATGG<br>TATATTTTGGAGCTAAAGAAAGAACTGAGGAGGAATGGGCAACCATCTTTCGAG<br>AAGCTGGTTTCGTCGATTACAAGATTTTTCCCCTGTTAGATTTCAGGAGTCCCAT<br>TGAACTGTATCCTTAA |
| ***Rv*UGT1** | **ATG**GGTTCCGAGTGCAGAGGTCCTCTTCATGTTTTCTTGGTCTCGTTCCCGGGAC<br>AAGGTCATTTTAATCCTCTGCTTAGACTTGGCAGGCTCCTTGCTTCAAGGGGGCT<br>GCTGGCTACCTTATCCGCGCCTGAACTCATAGGCAAAGACATAAAAAAAGCCAA<br>CAATATTGCCGATGATCAACCCATCGCAGTTGGTAGTGGTTTCGTCAGGTTCGA<br>GTTCTTTGATGATGAATCGGAGTCTAAAGCCTTCGATCTTGATACGTACTTGAAT<br>CATCTGGAGCTAGTCGGCAAGCAGAAACTCCCACAGATGCTCAAGAAATACGA<br>GGAACAAGGTCGCCGTGTTTCCTGCGTAATCGTCAATCCTTTCCTTCCCTGGGTT<br>TCTGATGTCGCCGAGAGCCTAAACATCCCAAGTGCCACGCTTTGGGTGCAGTCT<br>TGTGCCAGCTTCACTGCCTATTATCATTATCACCATCGTCTGGTGCCTTTCCCCA<br>GCGAAGCCGAGCCCGAAATCGATGTTCAACTGCCAGGCATGCCTTTGTTGAAGT<br>ATGATGAGGTGCCAAGTTTCTTGCACCCTACAAGTCCGTACCTGTGCCTGGCAA<br>GAGCTATCCTGGGCCAGTTCAAGAACTTCTCTAAGAACTTGTGCGTTTTGATGG<br>ACACGTTTCTGGAGCTTGAACATGACGAAATAGAGTCCGTTTCCAAGCTTTGTC<br>CCGTCAAACCTATAGGGCCTTTGTTCAAAGTTTCTAAAGATCCCAGCTCATCCAT<br>CAGCGGTGACATTATCAAGGCCGATGATTGCAAAGAGTGGTTAGACTCAAAACC<br>ACCGTCCTCCGTTGTATACATCTCTTTTGGCAGCGTAGTCTTCTTGAAGCAAGAG<br>CAAGTAACCGAAATAGCATACGGGCTTTTGAACTCGGAAGTTTCATTCTTATGG<br>GTCCTAAGGCCTCCGGGTAAATCAGGAGGCCACGAGCCACACGTTTTACCAGAA<br>GGATTCTTGGAAAAGGTCGGCGATAAGGGCAAGATTGTGCAGTGGAGTCCACA<br>AGAAGAGGTCTTGGGCCATCCTTCGGTTGCCTGTTTTTTGACGCACTGCGGTTGG<br>AACTCAACCCTGGAGGCTCTCGCTAGTGGGGTGCCGGTGATGGCTTTCCCTCAA<br>TGGGGTGATCAAGTCGTCGATGCAAAATACTTGGTGGACGTTTTCAAAGTTGGG<br>GTTAGAATGTCTAGAGGTGAGGCTGAGAACAGAATCATTCCCAGAGAAGAAGT<br>TGAGAAATGTTTGCGTGAGGCTACGTATGGTCCGAAGGCGGAAGAGATGAAAA<br>AGAACGCATCGACGTGGAAGAAGAAGGCAGAGGAAGCGGCGGCAAATGGTGG<br>TTCCTCCGACCGGAATATCCAAGATTTCCTGGACGAGATCAAGAAGAGATGTTC<br>AATTAAGCACTAG |
| ***Rv*UGT2** | **ATG**GCGAAGGTCGAAAATGATGAACTTCATGTGCTATTTGTTCCATACTTCACA<br>CCAAGCCATATGATTCCTCTAGTTGATGCAGCCAGGGTCTTTGCTGCTCATGGTG<br>TCAAGGTCACCATCATCGCTACCCCAGCAAATGCAGCCCTCTTCAAGTCCTCTGT<br>AGATCGGGACATCGAATTAGGCCAGAAAATCTCCGTTCAGACCATCCCCTTTCC<br>GGCAGCTGCAGTGGGCTTACTTGAGGGAATCGAGAACTTCAATCTTGCAACTTC<br>CATTGAAATGCTTGCCAAACTCATGAGAGCCGTTTTCATTCTCCAAAATCCGATC<br>GAGAAACTTATTCTCGATATTAATCCTCACTGCATCTTCTCCGACAGGTTACTGC<br>CTTGGACTGTCGACGTAGCCGAGAAATTGAAGATTCCGAGATTGTCATTTCATG<br>CTCTGAGTTTCTTTCATCACTGTTTGGGGCACAATTTGGAGATTTATGCACCCCA |

| | |
|---|---|
| | CAAGAACGCAGAATCTGATTCTGGGAGCTTCTTGGTTCCTGATTTACCAGACAA<br>GATTGAGCTGAAAAAGTCCCAGATCGAAGACTATGGTGCCAATGCTTTTGGGAA<br>AGTGATGAACATGGTCAAAGAAGCTGAGCTTCGGAGCTATGCTGCCGTTCATGA<br>TACTGTTCTCGAGCTGGAACCTGCTTATGCTGAGTACTGCGAGAAGGTAAGAGG<br>AAAAAAATCTTGGTCCATTGGCCCCCTTTTTCACTTCTCCAACAGAGAAAAATC<br>AAATCTTGAAGCTGATAAGGCAAGTTGTTTAAATTGGTTAGATTCTCAAGAACA<br>CAACAAAGTTCTTTATATTTGCTTTGGGAGTTTGGTCAACTTCCCAGATCCCCAA<br>CTAAAGGAGATTGCTTTAGCTCTAGAGGCCTCAAGCTGCCCATTTATTTGGGTTG<br>TCAGAACACAAAATGTGCAAGAAAGCTGGTTTCCAGATGGTTTCGAGGAAAAA<br>CTGATCAAGAATGGCAAAGGTTTAATCATCAAAGGCTGGGCTCCTCAGGTGAAA<br>ATCTTGAACCACCCAGCAATCGGTGGTTTCATGAGTCACTGCGGCTGGAACTCG<br>ACTCTGGAATCCATCACCGCCGGCGTTCCGATGATTACTTGGCCGTTGTTCGCCG<br>AGCAATTCTACAACGAGAAGTTCATTGAGGCGATGCAGTTCGGAGTCCGAGTCG<br>GCGCCGAGGTGAGAAACTTGATTCCGCAGATGATCACATCGCCATTGATAGGAA<br>GCAAGCAGATTCAGGAAGCGATTCAGCGTTTGATGTGCGGCTCAGAAGAAAGC<br>GTGCGAAGAAGGGAGATGGTTTTAGAAGCGGCTGCGATTTCAAAAAGGGCTGT<br>TGAAGAAGGCGGGTCATCTCATGAAAATGTCGTGGCTTTGATTGAGGAGATGAA<br>ATCGTTTGCATTTGGTAAGAAGAATTGA |

**Supplementary Table 2** Oligonucleotides used for the construction of 3Ω1/pOPINF/pOPINM expression plasmids destined for transient gene expression in *Nicotiana benthamiana* or *E. coli.*

| Oligonucleotides used for subcloning of gene sequences from cDNA and construction of 3Ω1 | |
|---|---|
| 3Ω1_*Rv*SGD (1)_F | TTTATGAATTTTGCAGCTCGATGGAAAGTAATCAAGGTGAGCCTCTTG |
| 3Ω1_*Rv*SGD (1)_R | GACAACCACAACAAGCACCGTTAGGTTTTTGCCTCTTGACTAACTC |
| 3Ω1_*Rv*SGD2_F | TTTATGAATTTTGCAGCTCGATGGCACTGCAATTACTTCTCTTTGC |
| 3Ω1_*Rv*SGD2_R | GACAACCACAACAAGCACCGCTATTTGAGGAGGAATTTCTTGTACC |
| 3Ω1_*Rv*SGD3_F | TTTATGAATTTTGCAGCTCGATGGCTTCCAAAGATGTCAGTTACTATAAG |
| 3Ω1_*Rv*SGD3_R | GACAACCACAACAAGCACCGTCATGTTTTCTGAAGGAACTTTTTGAACC |
| 3Ω1_*Rv*SGD4_F | TTTATGAATTTTGCAGCTCGATGCTGCCGAGAAAAGAAATTATTGC |
| 3Ω1_*Rv*SGD4_R | GACAACCACAACAAGCACCGTCATGTTTTCTGAAGGAACTTTTTGAACC |
| 3Ω1_*Rv*SGD5_F | TTTATGAATTTTGCAGCTCGATGGCTAAAGAAATCCTGTGCTTTCTTC |
| 3Ω1_*Rv*SGD5_R | GACAACCACAACAAGCACCGCTATTGTTTTCGCAGTAATTTCTGGAACC |
| 3Ω1_*Rv*SGD6_F | TTTATGAATTTTGCAGCTCGATGGTGCCTCGTCCATGCTTC |
| 3Ω1_*Rv*SGD6_R | GACAACCACAACAAGCACCGTCAATAAGAGAACCTGGTGTTTGAAG |
| 3Ω1_*Rv*YOS_F | TTTATGAATTTTGCAGCTCGATGGCTTCAGAGTCACCGGAAGATG |
| 3Ω1_*Rv*YOS_R | GACAACCACAACAAGCACCGTTATGCCGATTTGACAGTGTTTGCCATG |
| 3Ω1_*Rv*YOO _F | TTTATGAATTTTGCAGCTCGATGGAAACAAAAGTCCATATGGTTCTTTC |
| 3Ω1_*Rv*YOO_R | GACAACCACAACAAGCACCGCTATGCAGTAGCCAATCTCTTAAAG |
| 3Ω1_*Rv*DYR1_F | TTTATGAATTTTGCAGCTCGATGGATGCAGCATCTGCAAAAAC |

| | |
|---|---|
| 3Ω1_*Rv*DYR1_R | GACAACCACAACAAGCACCGCTAAGTAGATTTCAATGTATTCTCTATATC |
| 3Ω1_*Rv*DYR2_F | TTTATGAATTTTGCAGCTCGATGGACGCAGCATCTGCAAAAACAG |
| 3Ω1_*Rv*DYR2_R | GACAACCACAACAAGCACCGCTAAGCAGATTTCAATGTATTCTCTACATC |
| 3Ω1_*Rv*CYP71D820_F | TTTATGAATTTTGCAGCTCGATGGACCTTCAGCACTTGCCCTTTAAC |
| 3Ω1_*Rv*CYP71D820_R | GACAACCACAACAAGCACCGTTAACAGGATGTAGCTAGGGATGG |
| 3Ω1_*Rv*CYP72A270_F | TTTATGAATTTTGCAGCTCGATGGACAACATCTTCAACTTAATTGCAG |
| 3Ω1_*Rv*CYP72A270_R | GACAACCACAACAAGCACCGTTATAATTTGTGCAAGATAAGGTGAGCC |
| 3Ω1_*Rv*OMT11_F | TTTATGAATTTTGCAGCTCGATGGATTTGCCATCTTCTGAGATCCG |
| 3Ω1_*Rv*OMT11_R | GACAACCACAACAAGCACCGTTAAGGATACAGTTCAATGGGACTCC |
| Oligonucleotides used for subcloning of gene sequences and construction of pOPINF/pOPINM | |
| pOPINF_*Rv*SGD (1)_F | AAGTTCTGTTTCAGGGTACCATGGAAAGTAATCAAGGTGAGCCTCTTG |
| pOPINF_*Rv*SGD (1)_R | ACTGGTCTAGAAAGCTTTTAGGTTTTTTGCCTCTTGACTAACTC |
| pOPINF_*Rv*DYR1_F | AAGTTCTGTTTCAGGGTACCATGGATGCAGCATCTGCAAAAAC |
| pOPINF_*Rv*DYR1_R | ACTGGTCTAGAAAGCTTCTAAGTAGATTTCAATGTATTCTCTATATC |
| pOPINF_*Rv*DYR2_F | AAGTTCTGTTTCAGGGTACCATGGACGCAGCATCTGCAAAAACAG |
| pOPINF_*Rv*DYR2_R | ACTGGTCTAGAAAGCTTCTAAGCAGATTTCAATGTATTCTCTACATC |
| pOPINF_*Rv*OMT11_F | AAGTTCTGTTTCAGGGTACCATGGATTTGCCATCTTCTGAGATCCG |
| pOPINF_*Rv*OMT11_R | ACTGGTCTAGAAAGCTTTTAAGGATACAGTTCAATGGGACTCC |
| pOPINM_*Rv*UGT1_F | AAGTTCTGTTTCAGGGTACCATGGGTTCCGAGTGCAGAGGTC |
| pOPINM_*Rv*UGT1_R | ATGGTCTAGAAAGCTCTAGTGCTTAATTGAACATCTCTTCTTGATC |
| pOPINM_*Rv*UGT2_F | AAGTTCTGTTTCAGGGTACCATGGCGAAGGTCGAAAATGATGAACTTC |
| pOPINM_*Rv*UGT2_R | ATGGTCTAGAAAGCTTCAATTCTTCTTACCAAATGCAAACGATTTC |
| Oligonucleotides used for subcloning of gene sequences and construction of pESC-Ura | |
| pESC-U-*Rv*CYP71D820 | TCACTAAAGGGCGGCCGCACTAATGGACCTTCAGCACTTG |
| pESC-U-*Rv*CYP71D820 | CGTCATCCTTGTAATCCATCGATACACAGGATGTAGCTAGGGA |

**Supplementary Table 3** Oligonucleotides used for vector-specific primers of 3Ω1/pOPINF/pOPINM /pESC-Ura expression plasmids

| | |
|---|---|
| 3Ω1_seq_F | GATGAAAAAGCCCTAAAATTGGAG |
| 3Ω1_seq_R | ATTATTCACAAATGAGAAACAGAATGG |
| pOPINF_seq_F | TAATACGACTCACTATAGGG |
| pOPINF_seq_R | TAGCCAGAAGTCAGATGCT |
| pOPINM_seq_F | GAAATCATGCCGAACATC |
| pOPINM_seq_R | TAGCCAGAAGTCAGATGCT |
| pESC-Ura-GAL1_F | ATTTTCGGTTTGTATTACTTC |
| pESC-Ura-GAL1_R | GTTCTTAATACTAACATAACT |
| pESC-Ura-GAL10_F | GGTGGTAATGCCATGTAATATG |
| pESC-Ura-GAL10_R | GGCAAGGTAGACAAGCCGACAAC |

**Supplementary Table 4** $^{1}$H (400 MHz) and $^{13}$C (100 MHz) NMR Data assignments for 3-dehydro-*a*-yohimbine (**3**) (in $CD_3OD$, ($\delta$ in ppm, $J$ in Hz)).

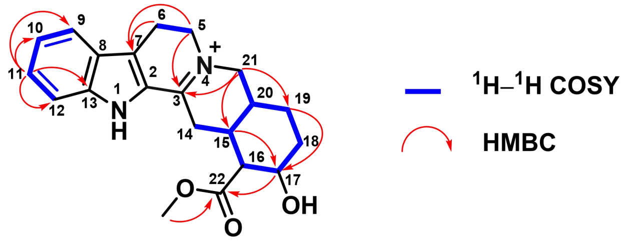

Key $^{1}$H–$^{1}$H COSY and HMBC correlations of **3**

| no. | **3**[a] | |
|---|---|---|
| | $\delta_C$ | $\delta_H$ (mult., J) |
| 2 | 127.7 | - |
| 3 | 166.1 | - |
| 5 | 54.3 | 4.01-4.12[a] |
| 6 | 20.2 | 3.25-3.32[a] |
| 7 | 123.9 | - |
| 8 | 125.4 | - |
| 9 | 114.2 | 7.49 (d, 8.5) |
| 10 | 122.8 | 7.16 (dd, 8.5, 7.2) |
| 11 | 129.7 | 7.42 (dd, 8.3, 7.2) |
| 12 | 122.5 | 7.68 (d, 8.3) |
| 13 | 142.8 | - |
| 14 | 26.5 | 3.25-3.32[a] |
| 15 | 31.6 | 2.77 (m) |
| 16 | 54.1 | 2.64 (dd, 10.5, 4.2) |
| 17 | 66.4 | 4.01-4.12[a] |
| 18 | 34.7 | 2.09 (m); 1.44 (ddd, 23.8, 11.8, 5.1) |
| 19 | 24.9 | 1.60-1.75[a] |
| 20 | 34.3 | 2.27 (m) |
| 21 | 58.5 | 4.16 (dd, 15.7, 3.8); 3.74 (d, 15.7) |
| 22 | 174.5 | - |
| 22-$OCH_3$ | 52.5 | 3.78 (s) |

[a]Overlapped signals were reported without designating multiplicity.

**Supplementary Table 5** Comparison of $^{13}C$ NMR Data of **12** and **5** (100 MHz) in this work and literature[7, 8]

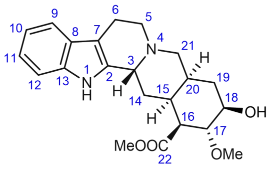

Deserpidic acid methyl ester (**12**)

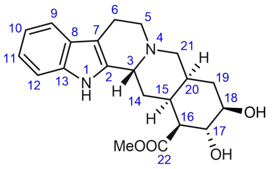

18$\beta$-Hydroxy-3-epi-$\alpha$-yohimbine(**5**)

| no. | This work, $\delta$ [ppm] in $CD_3OD$ | Literature, $\delta$ [ppm] in $CDCl_3$ | This work, $\delta$ [ppm] in $CD_3OD$ | Literature, $\delta$ [ppm] in $CDCl_3$ |
|---|---|---|---|---|
| 2 | 127.4 | 132.1 | 127.9 | 132.0 |
| 3 | 56.8 | 61.2 | 56.6 | 53.8 |
| 5 | 52.2 | 52.1 | 52.3 | 51.0 |
| 6 | 16.8 | 16.9 | 16.9 | 16.6 |
| 7 | 106.9 | 108.3 | 107.0 | 108.3 |
| 8 | 127.8 | 127.8 | 128.1 | 127.6 |
| 9 | 119.1 | 118.3 | 119.0 | 118.0 |
| 10 | 119.1 | 119.7 | 120.7 | 119.6 |
| 11 | 123.5 | 121.7 | 123.3 | 121.7 |
| 12 | 112.5 | 111.1 | 112.5 | 110.9 |
| 13 | 138.5 | 135.7 | 138.4 | 135.7 |
| 14 | 24.3 | 24.5 | 24.5 | 24.1 |
| 15 | 32.0 | 32.4 | 32.3 | 32.3 |
| 16 | 52.5 | 53.9 | 53.2 | 51.4 |
| 17 | 75.4 | 75.3 | 72.1 | 71.4 |
| 18 | 82.1 | 81.6 | 75.0 | 74.0 |
| 19 | 33.2 | 33.0 | 32.6 | 31.1 |
| 20 | 34.0 | 34.7 | 34.3 | 34.0 |
| 21 | 49.9 | 49.5 | 50.1 | 49.2 |
| 22 | 173.7 | 173.7 | 173.8 | 172.8 |
| 17-$OCH_3$ | 52.2 | 51.3 | - | - |
| 22-$OCH_3$ | 52.4 | 51.5 | 52.5 | 52.1 |

**Supplementary Table 6** $^{1}H$ (400 MHz) and $^{13}C$ (100 MHz) NMR Data assignments for reserpic acid methyl ester (**10**) and 11, 18*β*-hydroxy-3-epi-*α*-yohimbine (**6**) (in $CD_3OD$, *δ* in ppm, *J* in Hz).

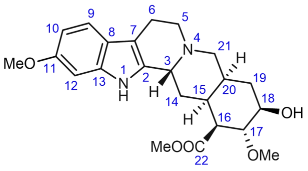

Reserpic acid methyl ester (**10**)

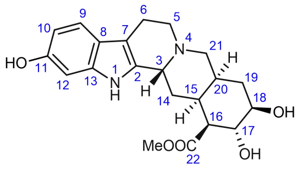

11, 18β-Hydroxy-3-epi-*α*-yohimbine (**6**)

| no. | $\delta_H$ | $\delta_C$ | $\delta_H$ | $\delta_C$ |
|---|---|---|---|---|
| 2 | - | 126.6 | - | 125.1 |
| 3 | - | 56.7 | - | 57.1 |
| 5 | 3.38-3.51[a] | 52.4 | 3.51-3.64 (m) | 52.5 |
| 6 | 3.06 (m); 2.86 (dd, 17.0, 3.6) | 17.0 | 3.06[a]; 2.94 (dd, 17.0, 5.0) | 16.9 |
| 7 | - | 107.0 | - | 106.8 |
| 8 | - | 122.3 | - | 121.6 |
| 9 | 7.30 (d, 8.6) | 119.6 | 7.25 (d, 8.4) | 119.6 |
| 10 | 6.69 (dd, 8.6, 2.0) | 110.7 | 6.61 (dd, 8.4, 2.1) | 111.0 |
| 11 | - | 158.2 | - | 155.2 |
| 12 | 6.86 (d, 2.0) | 95.9 | 6.75 (d, 2.1) | 98.0 |
| 13 | - | 139.2 | - | 139.7 |
| 14 | 2.30 (m); 2.16 (br d, 13.8) | 24.5 | 2.21-2.36[a] | 24.3 |
| 15 | 1.93-2.03[a] | 32.2 | 2.04-2.1[a] | 31.9 |
| 16 | 2.53 (dd, 10.4, 4.4) | 52.3 | 2.56 (dd, 11.0, 4.4) | 53.0 |
| 17 | 3.44 (dd, 11.5, 9.3) | 82.2 | 3.75 (dd, 11.5, 9.3) | 72.0 |
| 18 | 3.38-3.51[a] | 75.5 | 3.39 (ddd, 20.1, 10.1, 5.1) | 74.8 |
| 19 | 1.93-2.03[a]; 1.76 (br d, 10.0) | 33.3 | 1.92 (dd, 24.4, 12.8); 1.84 (dt, 12.8, 3.8) | 32.5 |
| 20 | 1.93-2.03[a] | 34.2 | 2.04-2.11[a] | 34.0 |
| 21 | 3.38-3.51[a]; 2.97 (dd, 12.5) | 49.9 | 3.53 (dd, 12.7, 3.9); 3.11 (d,12.7) | 50.0 |
| 22 | - | 173.8 | - | 173.7 |
| 11-$OCH_3$ | 3.78 (s) | 56.0 | - | - |
| 17-$OCH_3$ | 3.55 (s) | 61.4 | - | - |
| 22-$OCH_3$ | 3.81 (s) | 52.5 | 3.81 (s) | 52.5 |

[a]Overlapped signals were reported without designating multiplicity.

**Supplementary Table 7** Comparison of $^{13}$C NMR Data of **9** (100 MHz) in this work and literature[9].

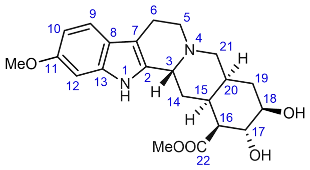

Rauvomitorine G (**9**)

| no. | This work, $\delta$ [ppm], in $CD_3OD$ | Literature, $\delta$ [ppm], in $CD_3OD$ |
|---|---|---|
| 2 | 125.5 | 129.9 |
| 3 | 56.8 | 55.7 |
| 5 | 52.3 | 52.3 |
| 6 | 16.8 | 17.3 |
| 7 | 106.8 | 107.5 |
| 8 | 122.1 | 123.0 |
| 9 | 119.8 | 119.2 |
| 10 | 110.9 | 109.9 |
| 11 | 158.4 | 157.6 |
| 12 | 95.9 | 96.0 |
| 13 | 139.4 | 138.7 |
| 14 | 24.2 | 25.0 |
| 15 | 31.9 | 33.4 |
| 16 | 53.0 | 53.7 |
| 17 | 72.0 | 72.3 |
| 18 | 74.8 | 75.5 |
| 19 | 32.5 | 33.2 |
| 20 | 34.0 | 35.4 |
| 21 | 49.8 | 50.3 |
| 22 | 173.7 | 174.3 |
| 11-$OCH_3$ | 56.0 | 56.1 |
| 17-$OCH_3$ | - | - |
| 22-$OCH_3$ | 52.5 | 52.3 |

### 4. NMR Spectra

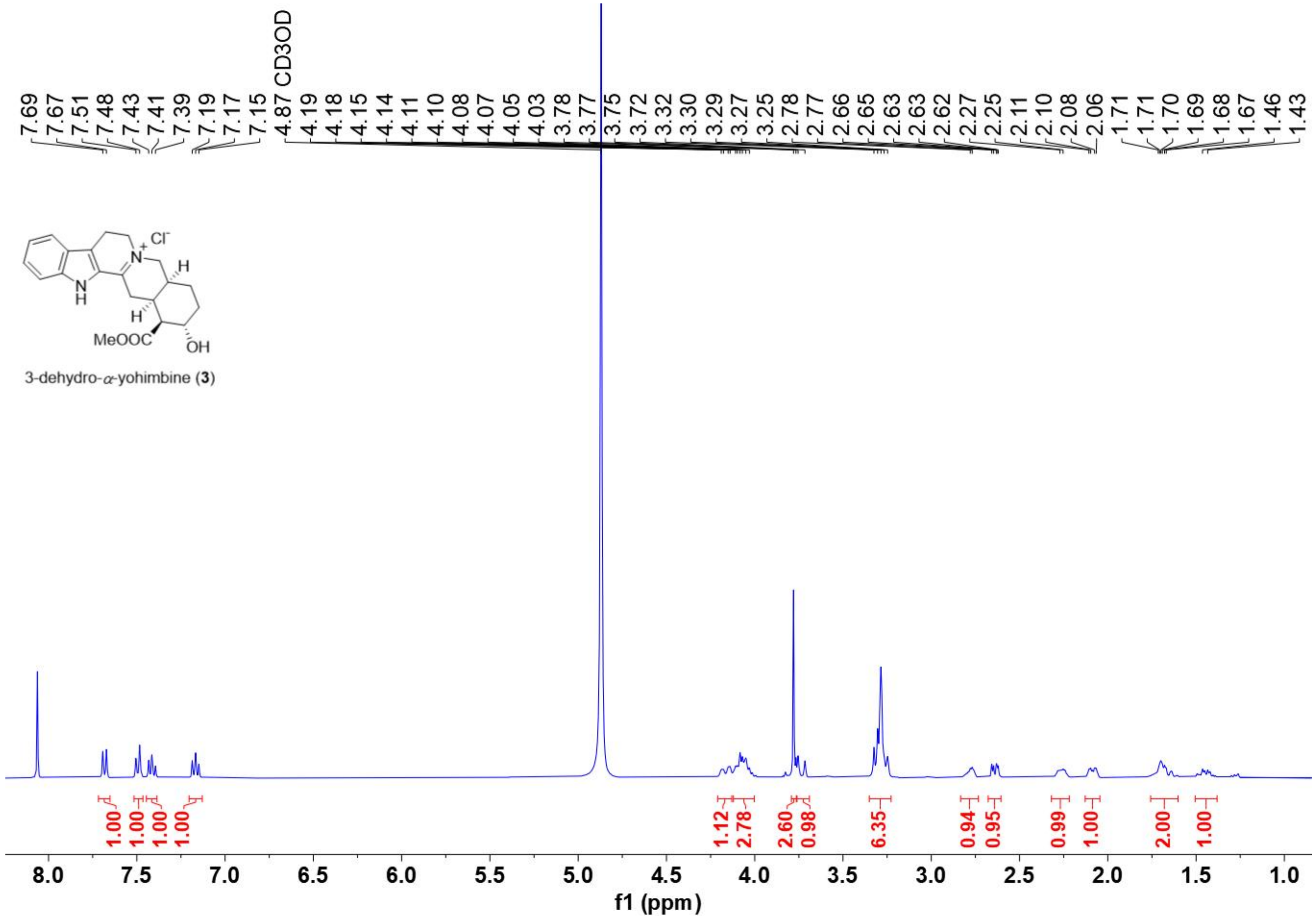

[1]H NMR spectrum of **3** (recorded in $CD_3OD$ at 400 MHz)

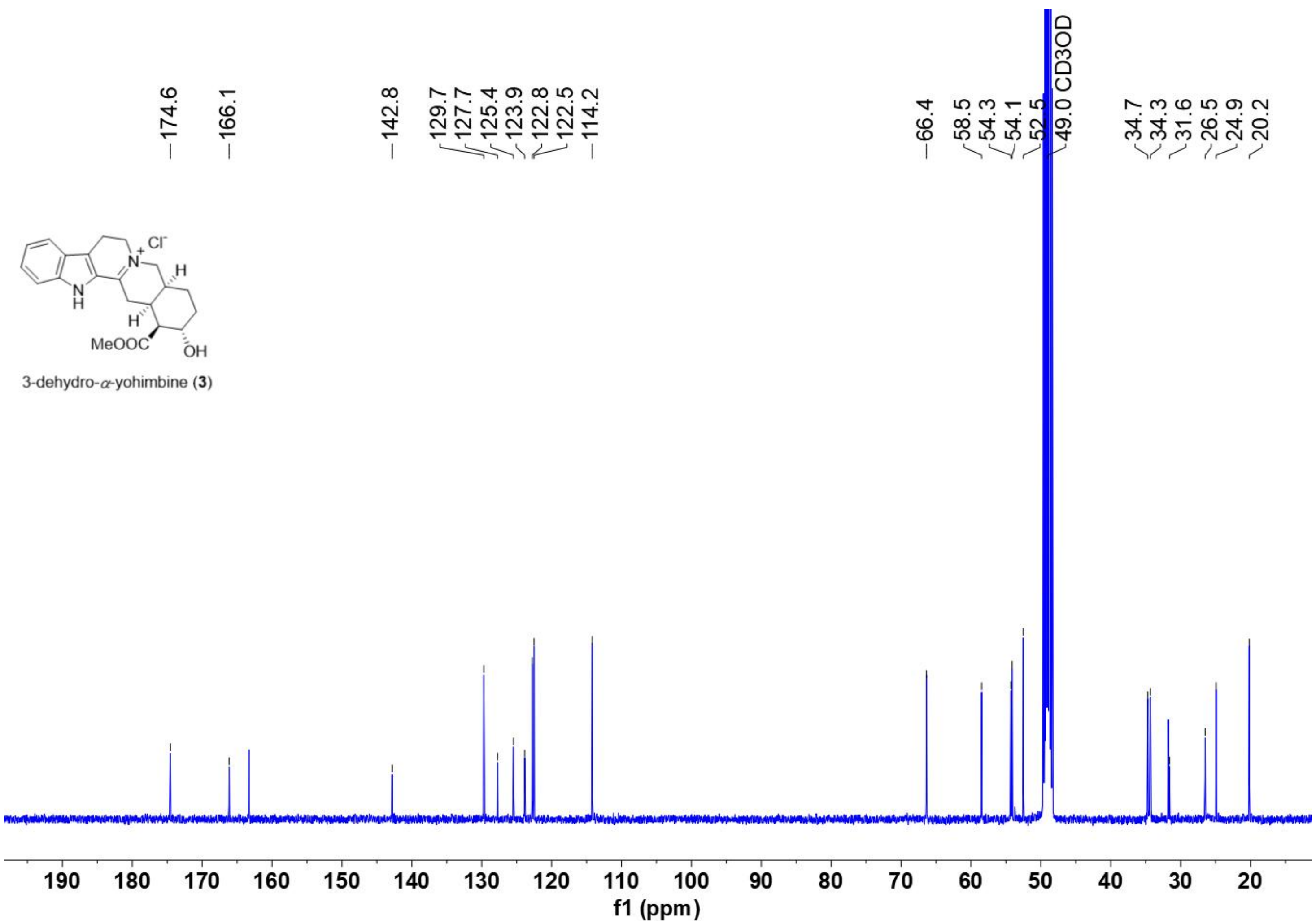

$^{13}C$ NMR spectrum of **3** (recorded in $CD_3OD$ at 100 MHz)

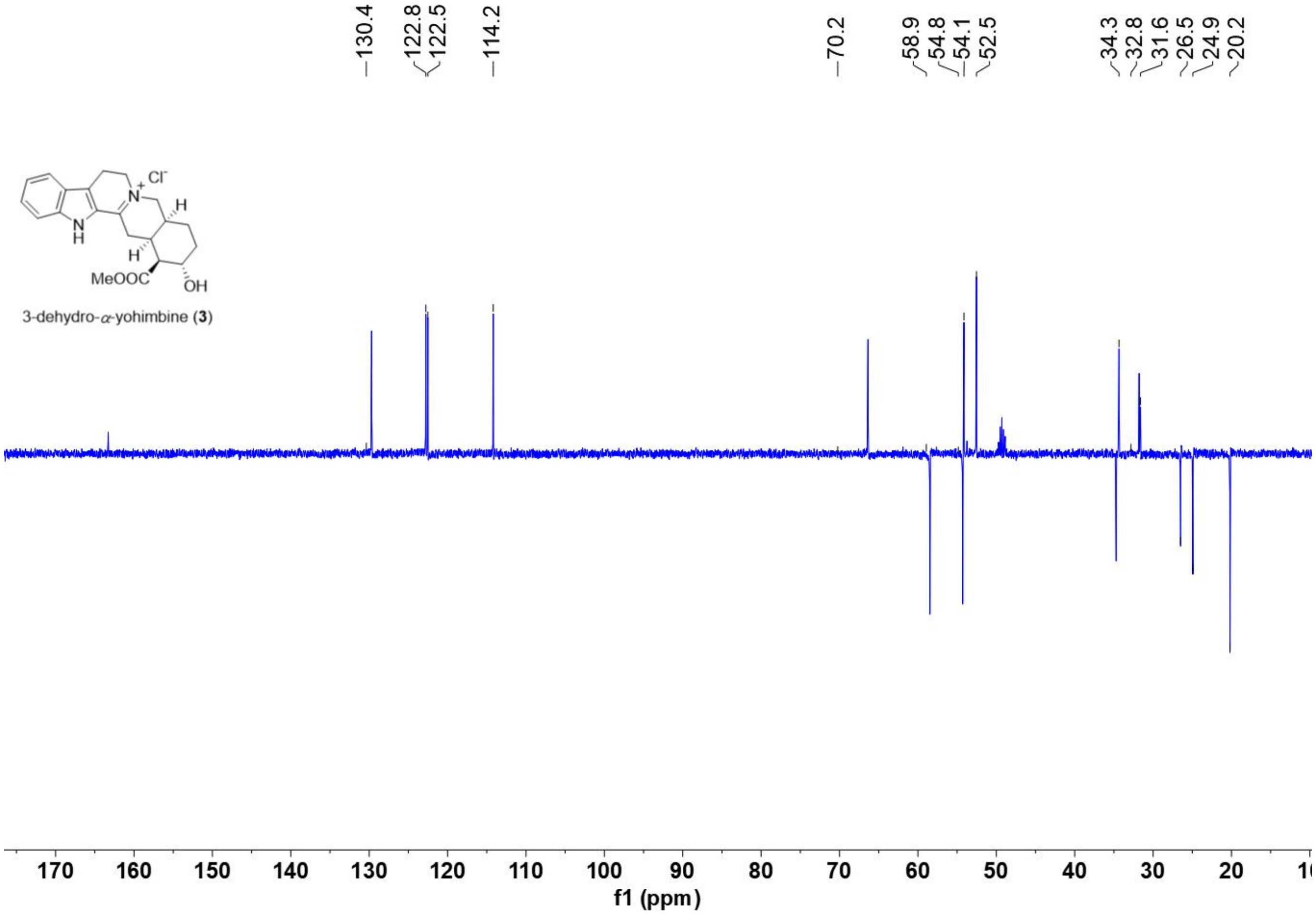

DEPT-135 NMR spectrum of **3** (recorded in $CD_3OD$ at 100 MHz)

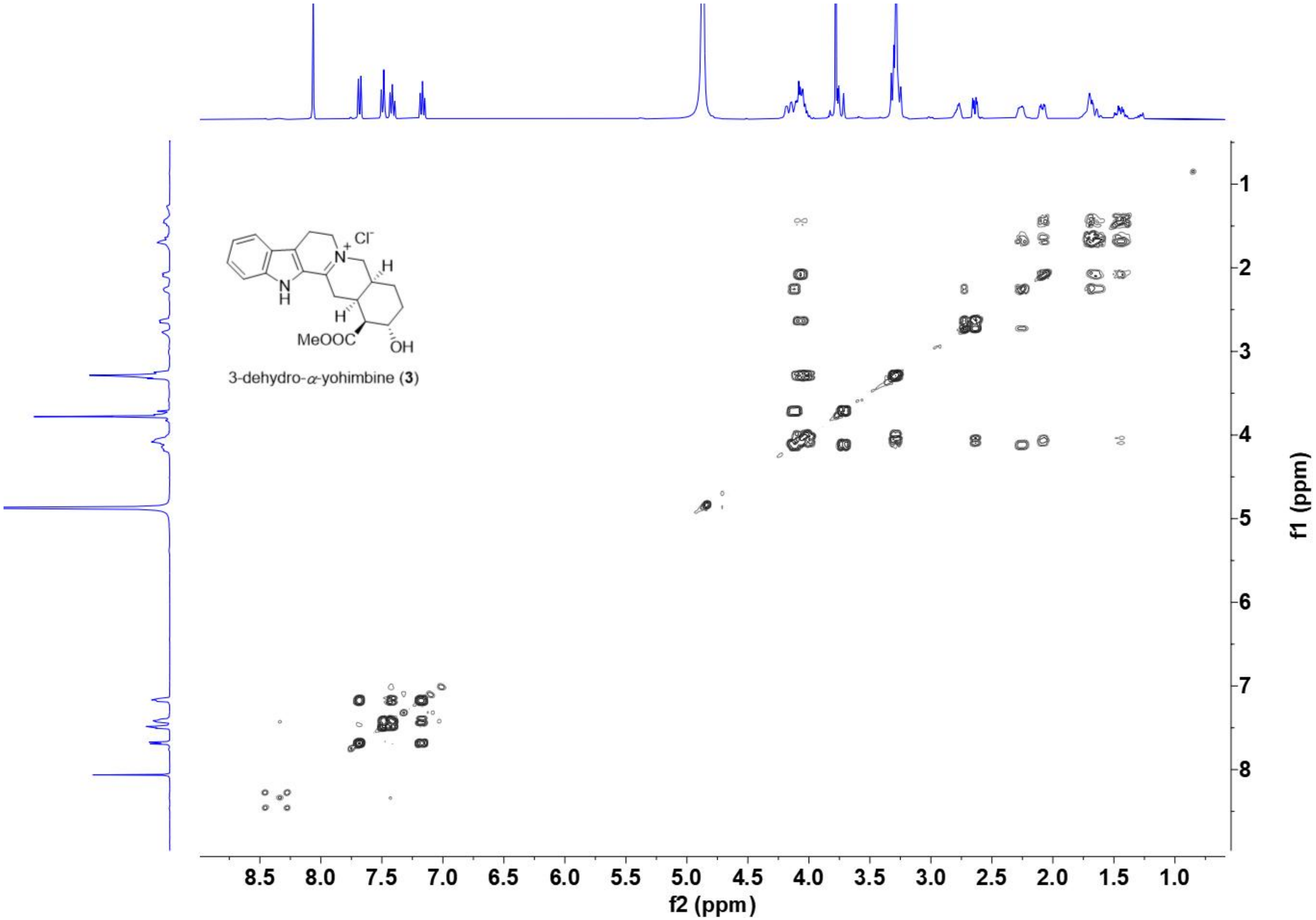

[1]H-[1]H COSY spectrum of **3** (recorded in $CD_3OD$ at 400 MHz)

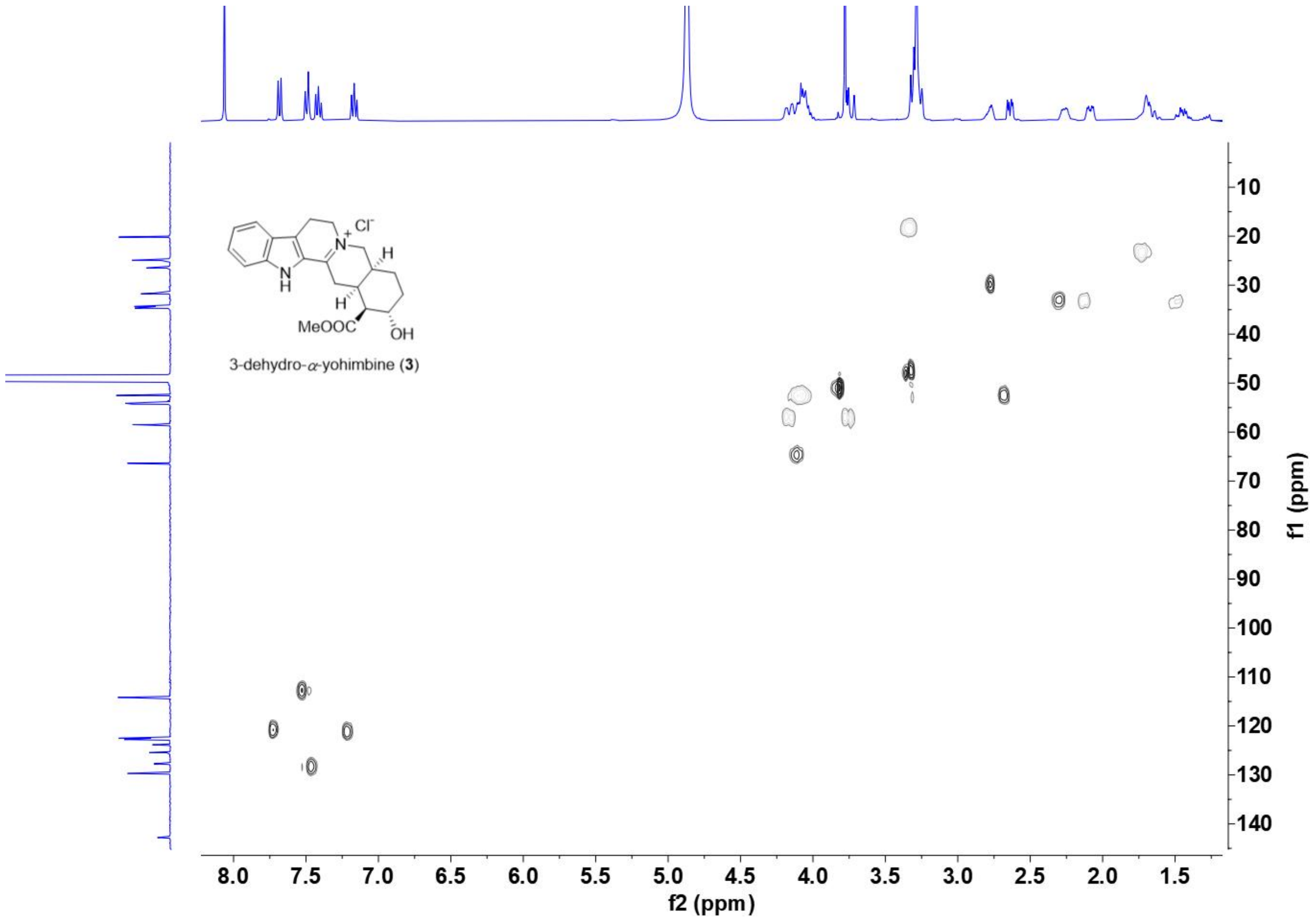

HSQC spectrum of **3** (recorded in $CD_3OD$ at 400 MHz)

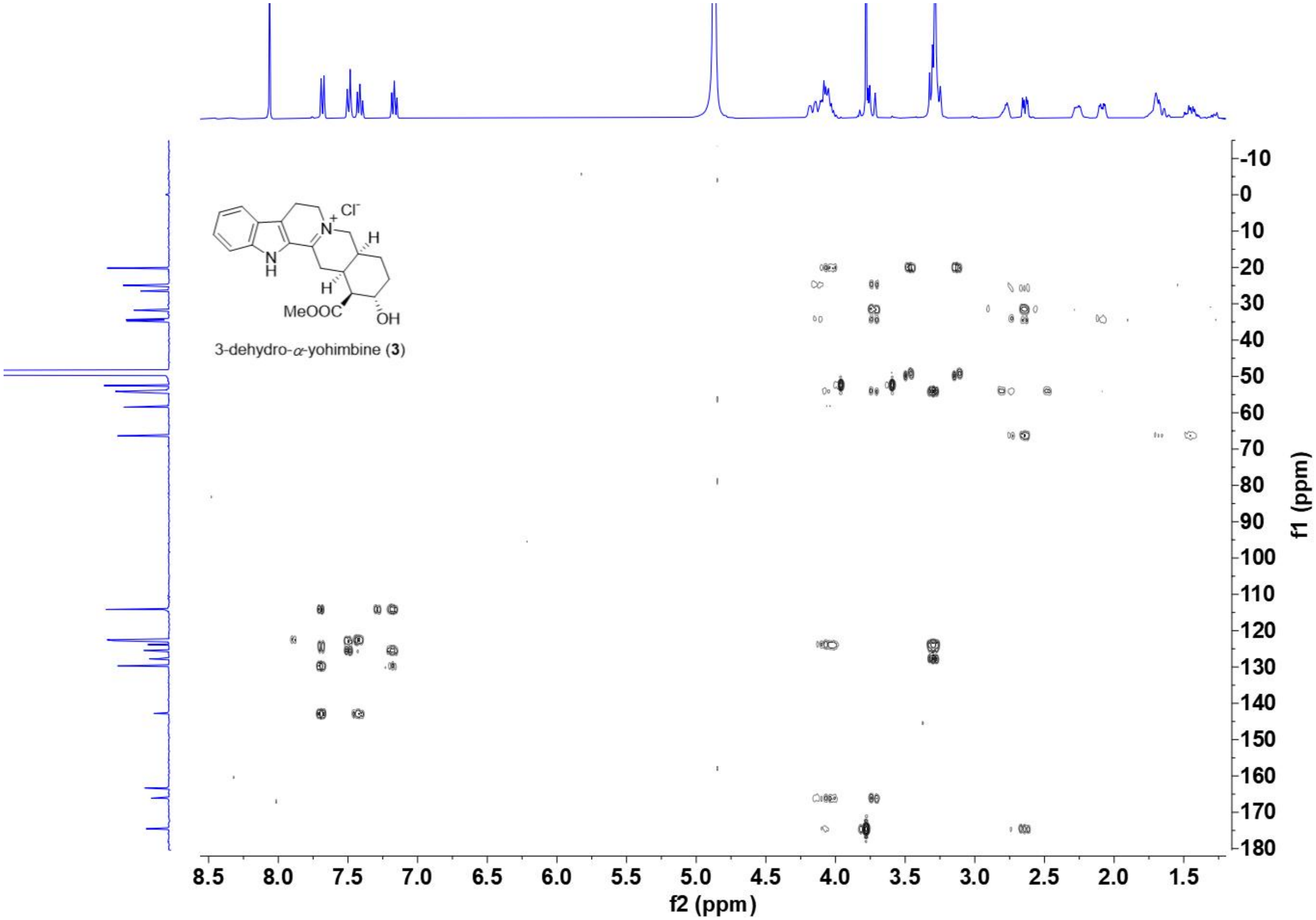

HMBC spectrum of **3** (recorded in $CD_3OD$ at 400 MHz)

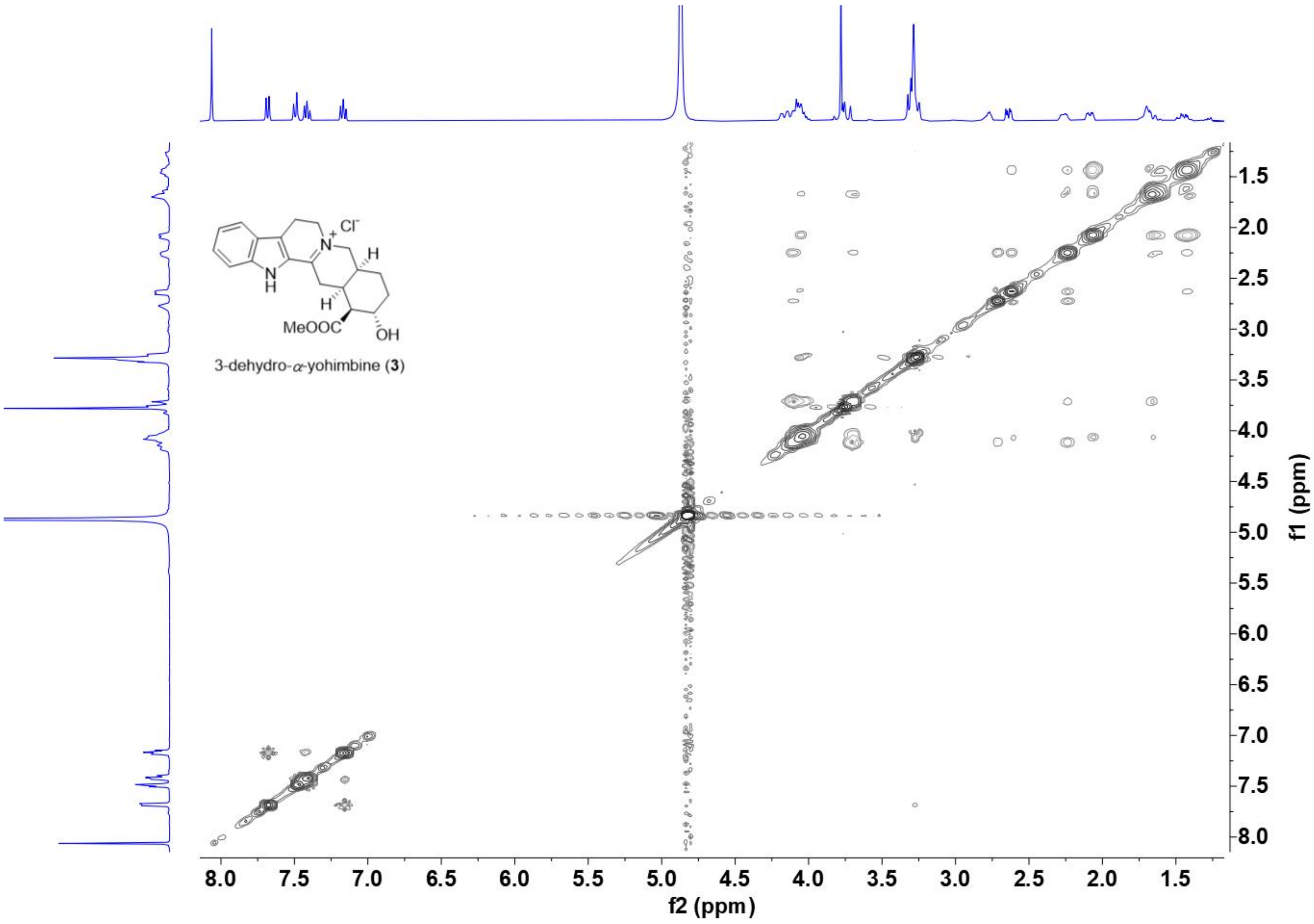

NOSEY spectrum of **3** (recorded in $CD_3OD$ at 400 MHz)

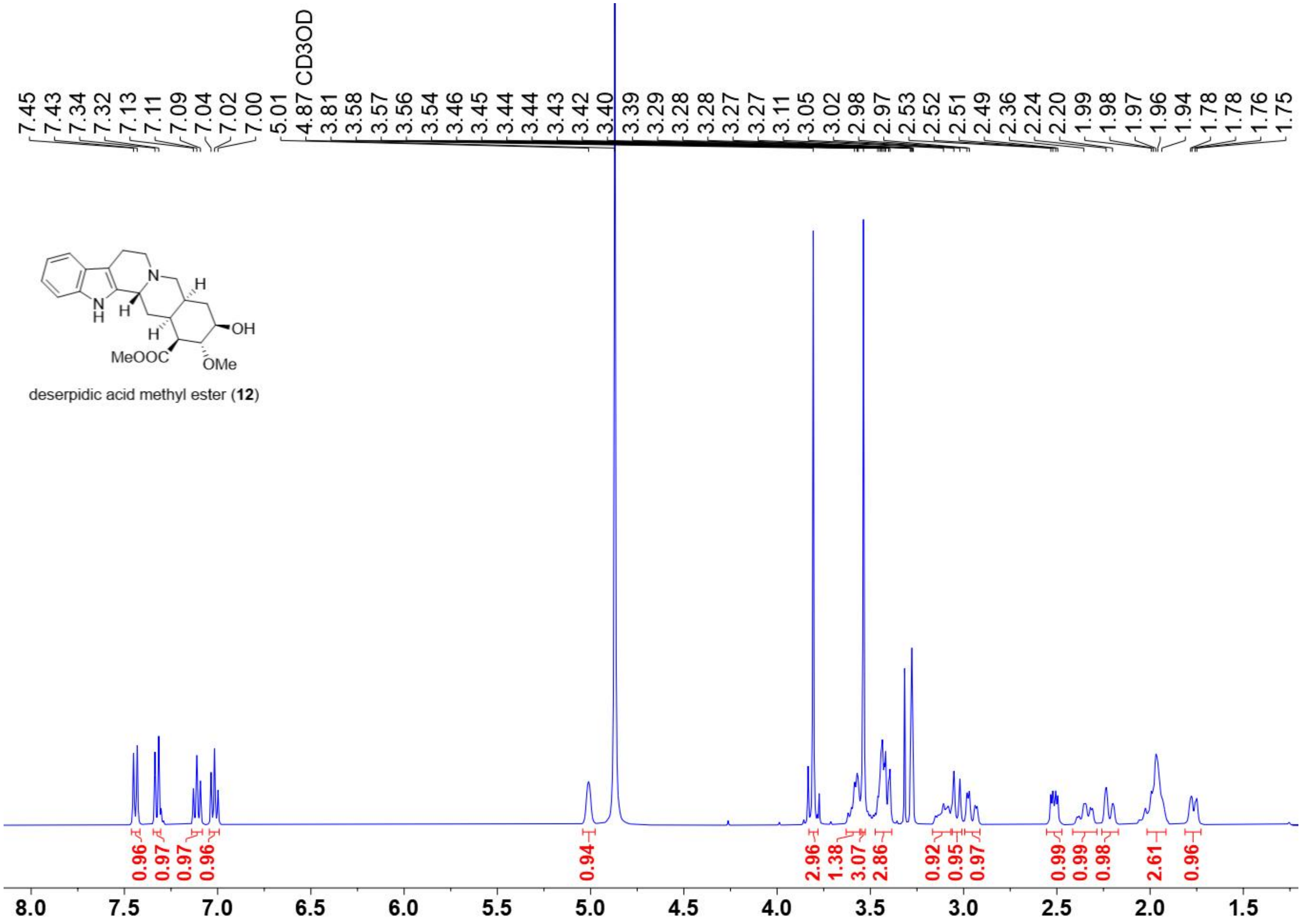

deserpidic acid methyl ester (**12**)

[1]H NMR spectrum of **12** (recorded in $CD_3OD$ at 400 MHz)

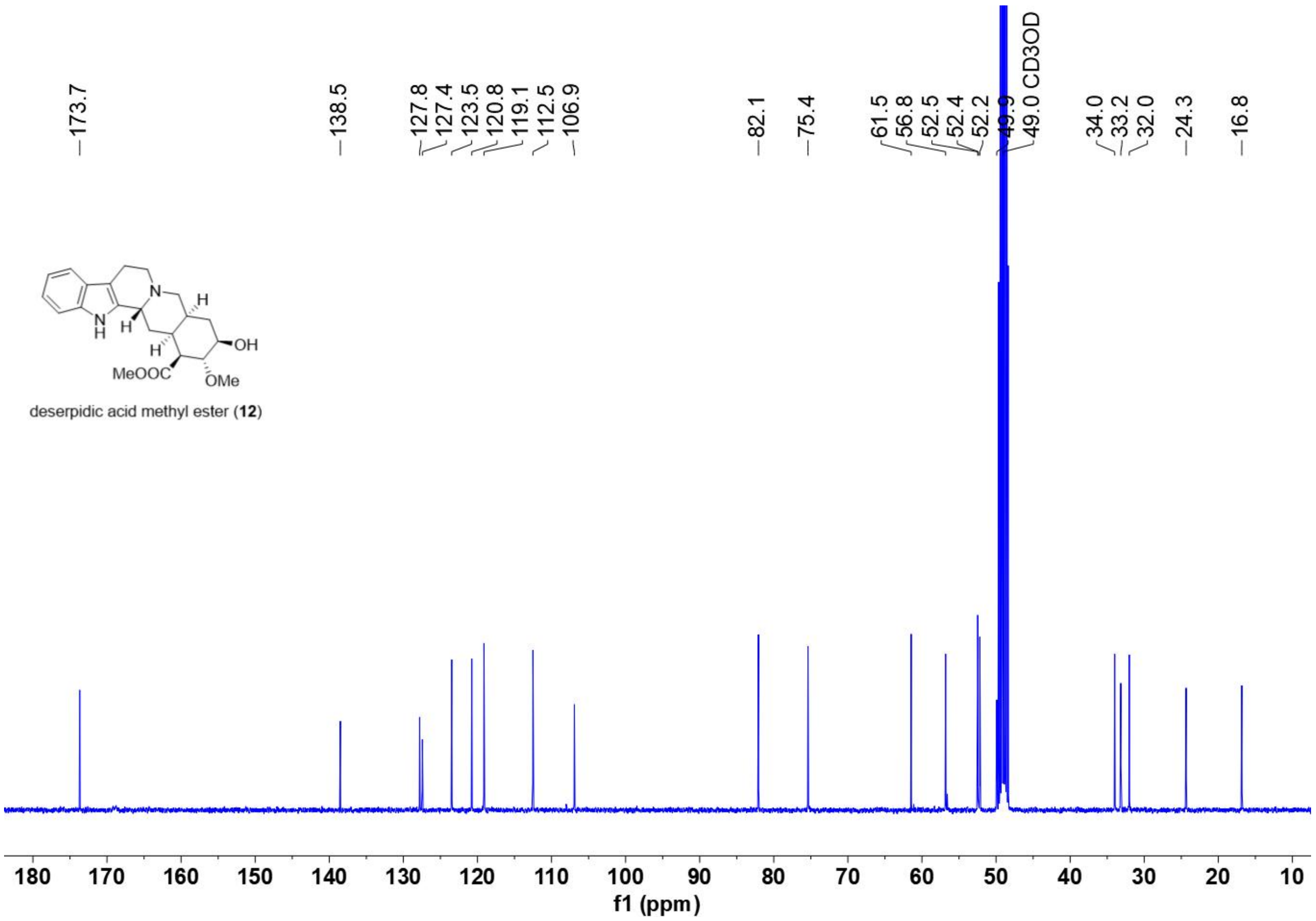

$^{13}C$ NMR spectrum of **12** (recorded in $CD_3OD$ at 100 MHz)

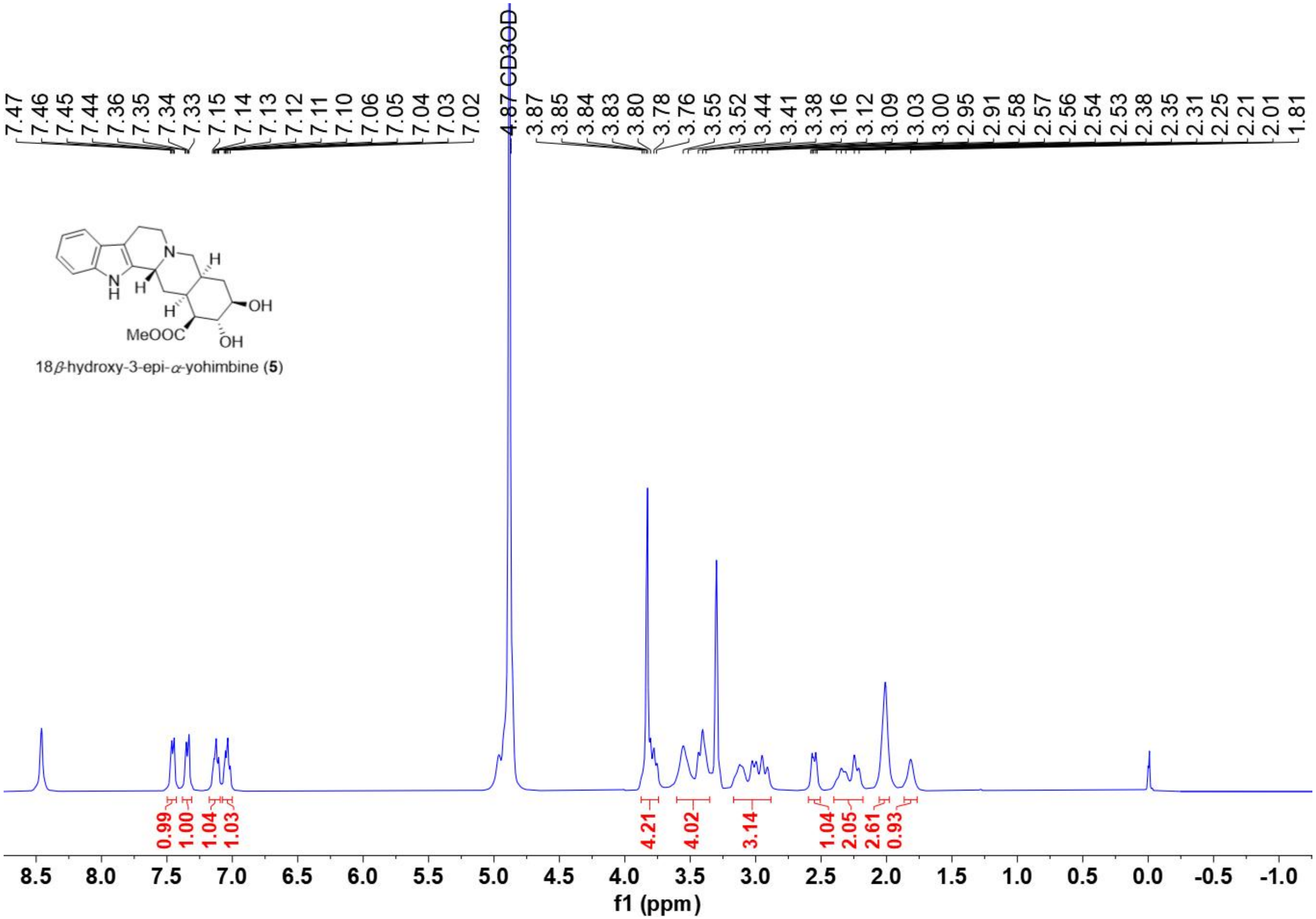

[1]H NMR spectrum of **5** (recorded in $CD_3OD$ at 400 MHz)

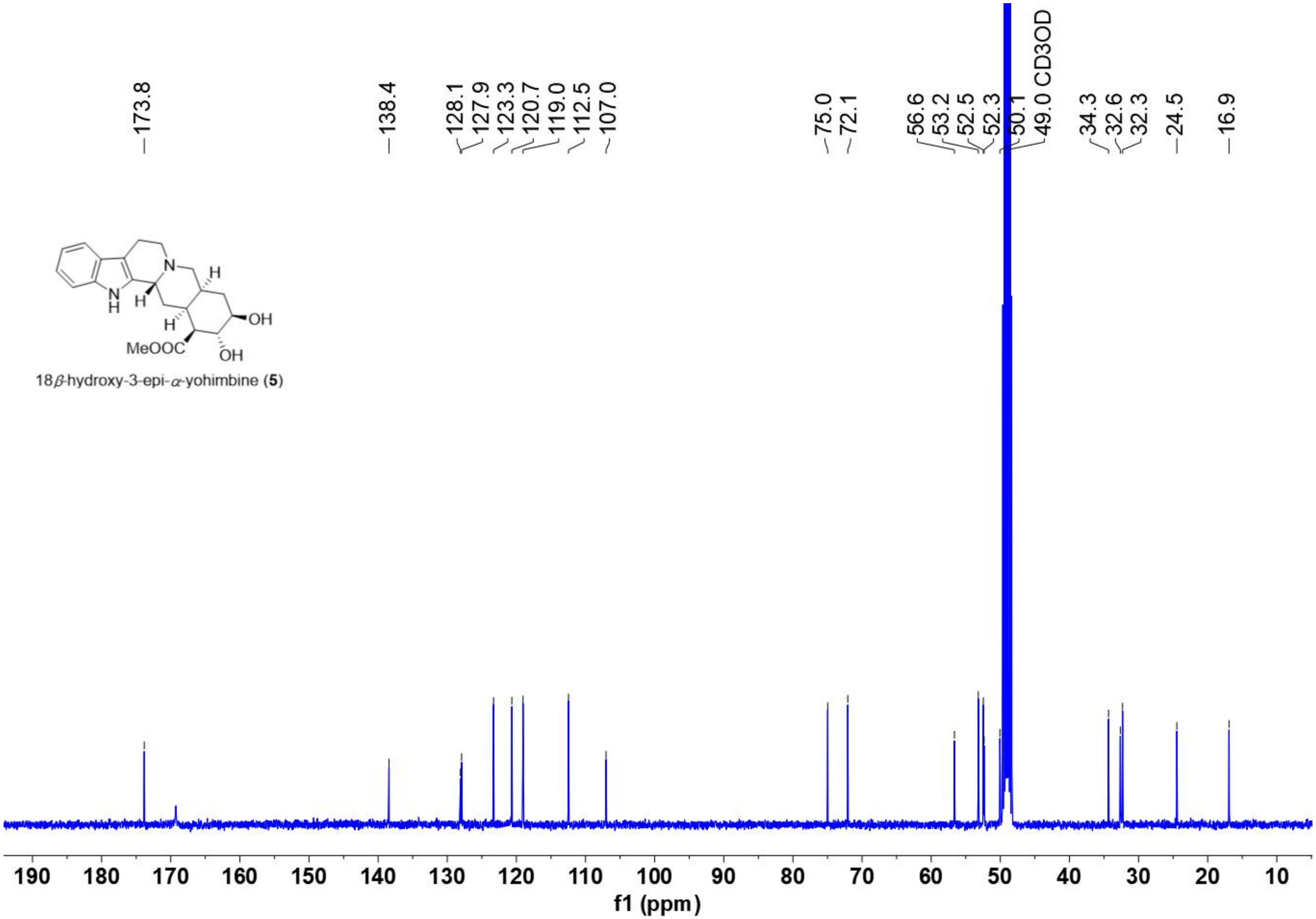

$^{13}$C NMR spectrum of **5** (recorded in $CD_3OD$ at 100 MHz)

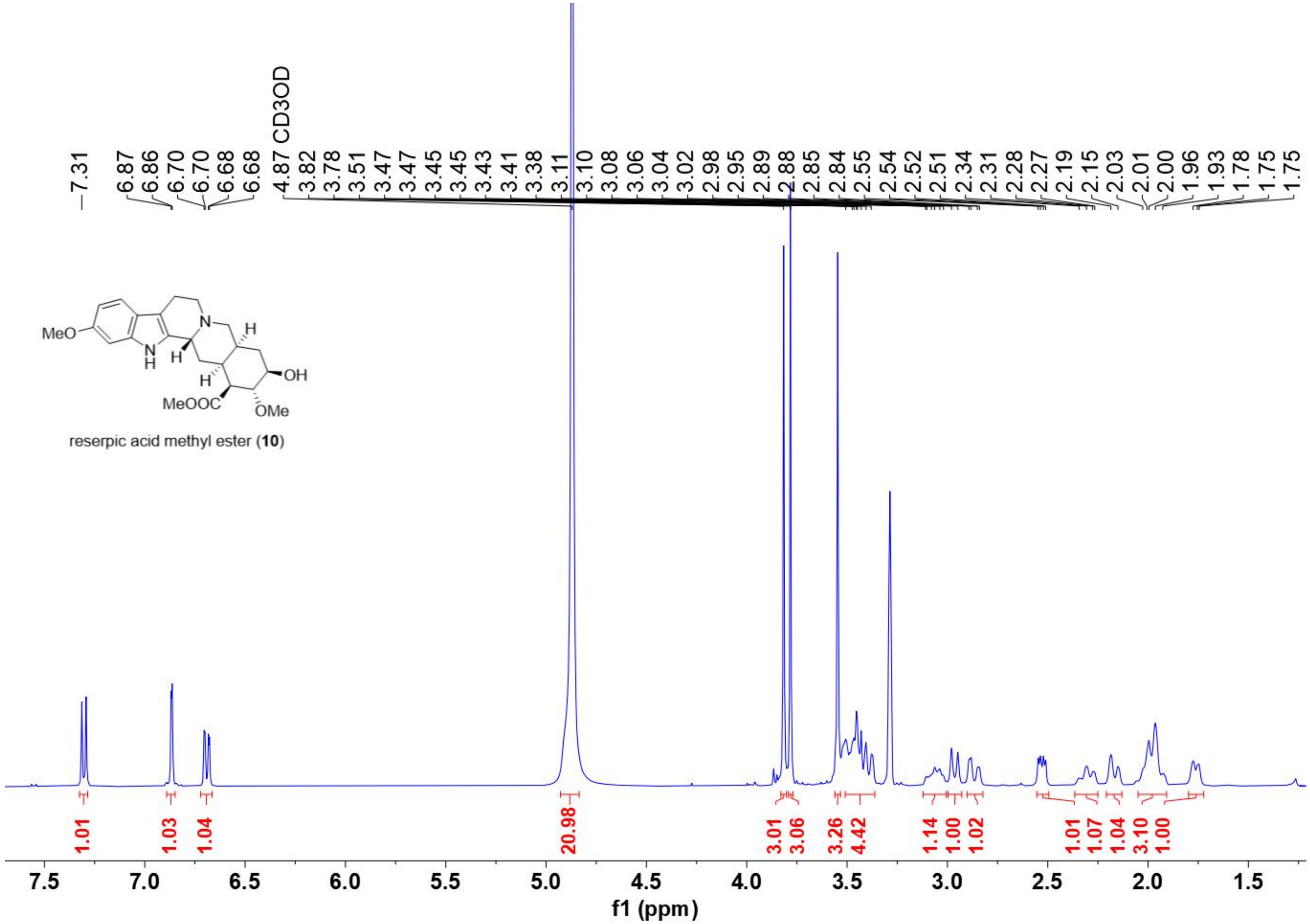

$^1$H NMR spectrum of **10** (recorded in $CD_3OD$ at 400 MHz)

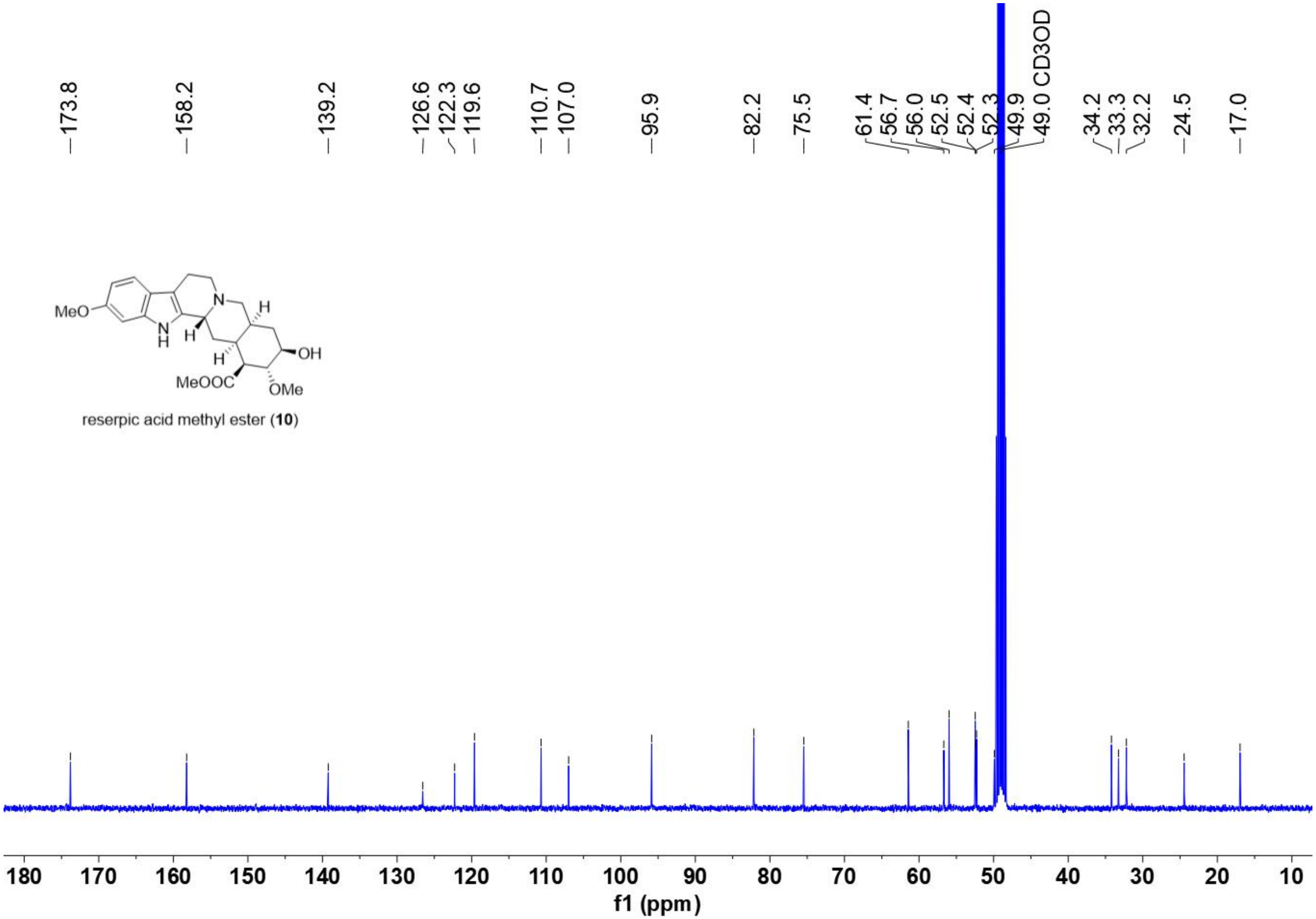

$^{13}C$ NMR spectrum of **10** (recorded in $CD_3OD$ at 100 MHz)

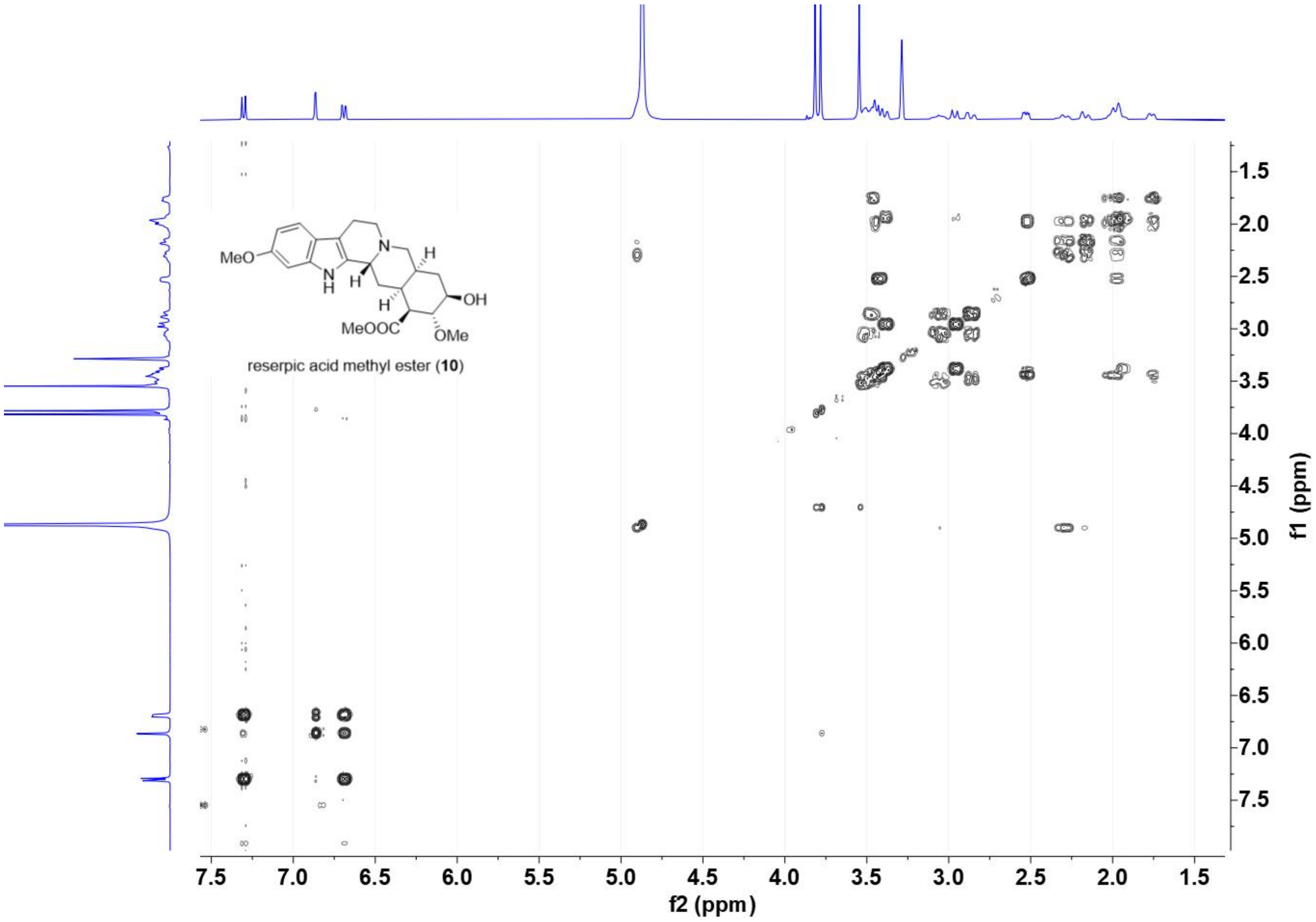

[1]H-[1]H COSY spectrum of **10** (recorded in $CD_3OD$ at 400 MHz)

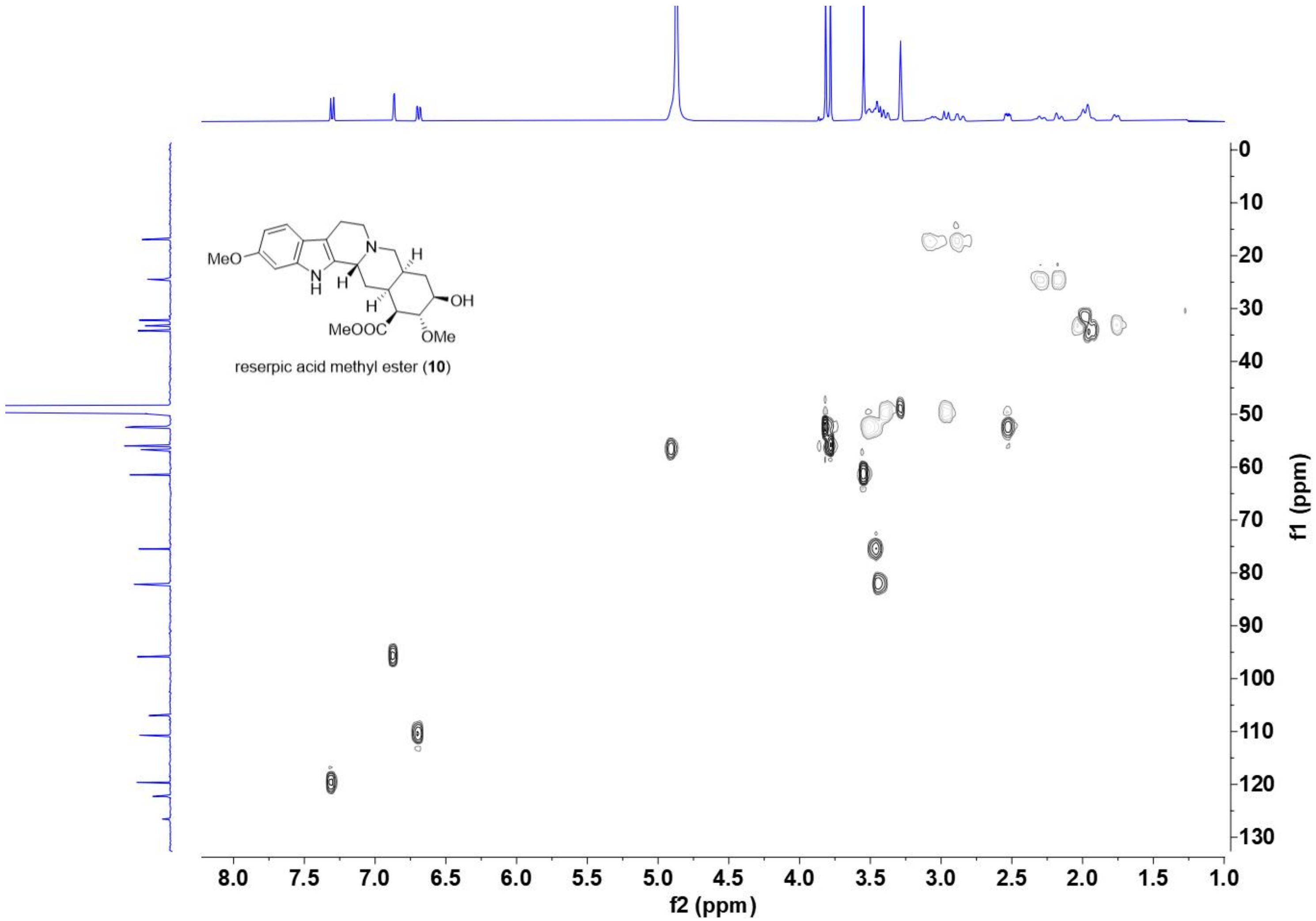

HSQC spectrum of **10** (recorded in $CD_3OD$ at 400 MHz)

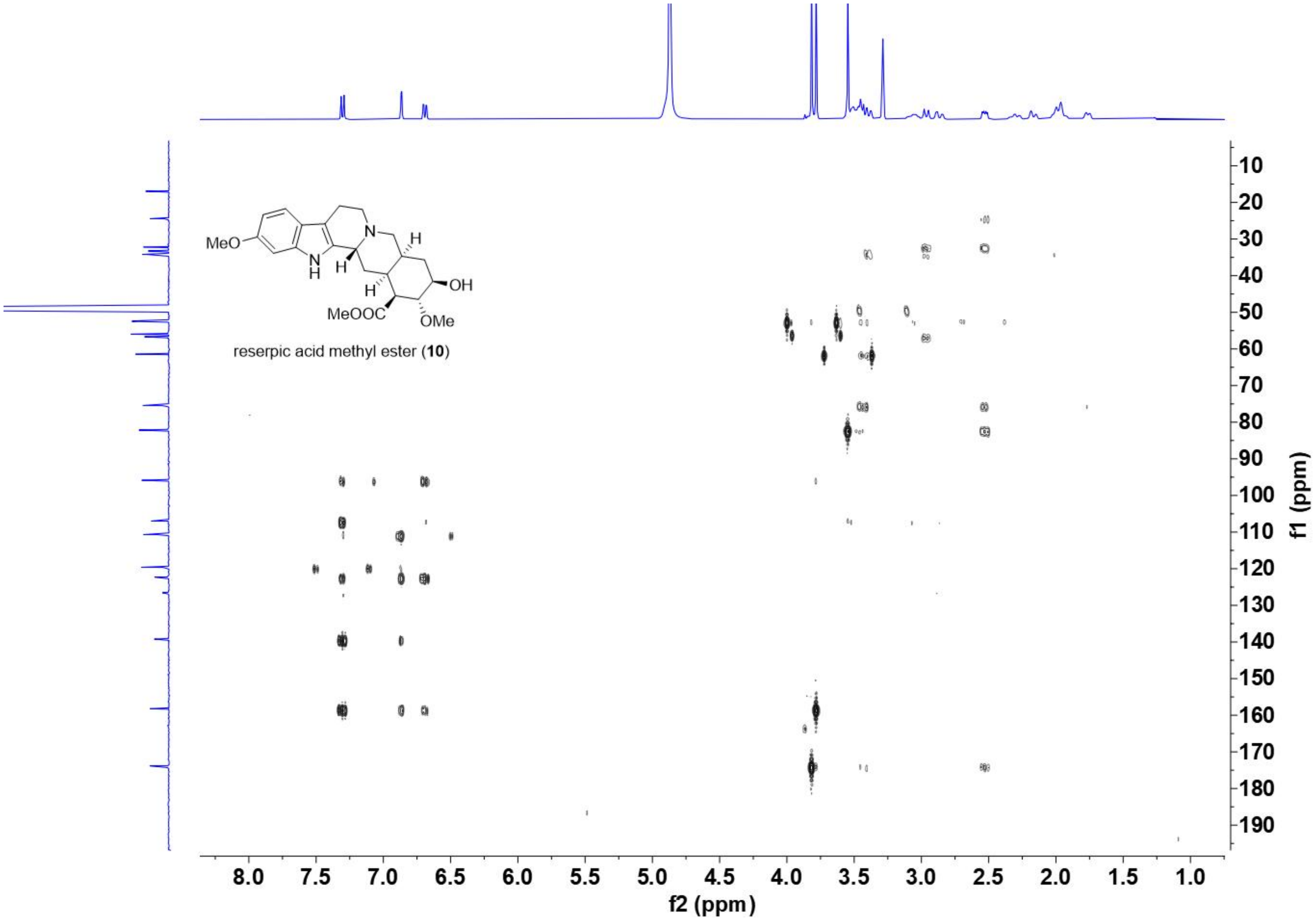

HMBC spectrum of **10** (recorded in $CD_3OD$ at 400 MHz)

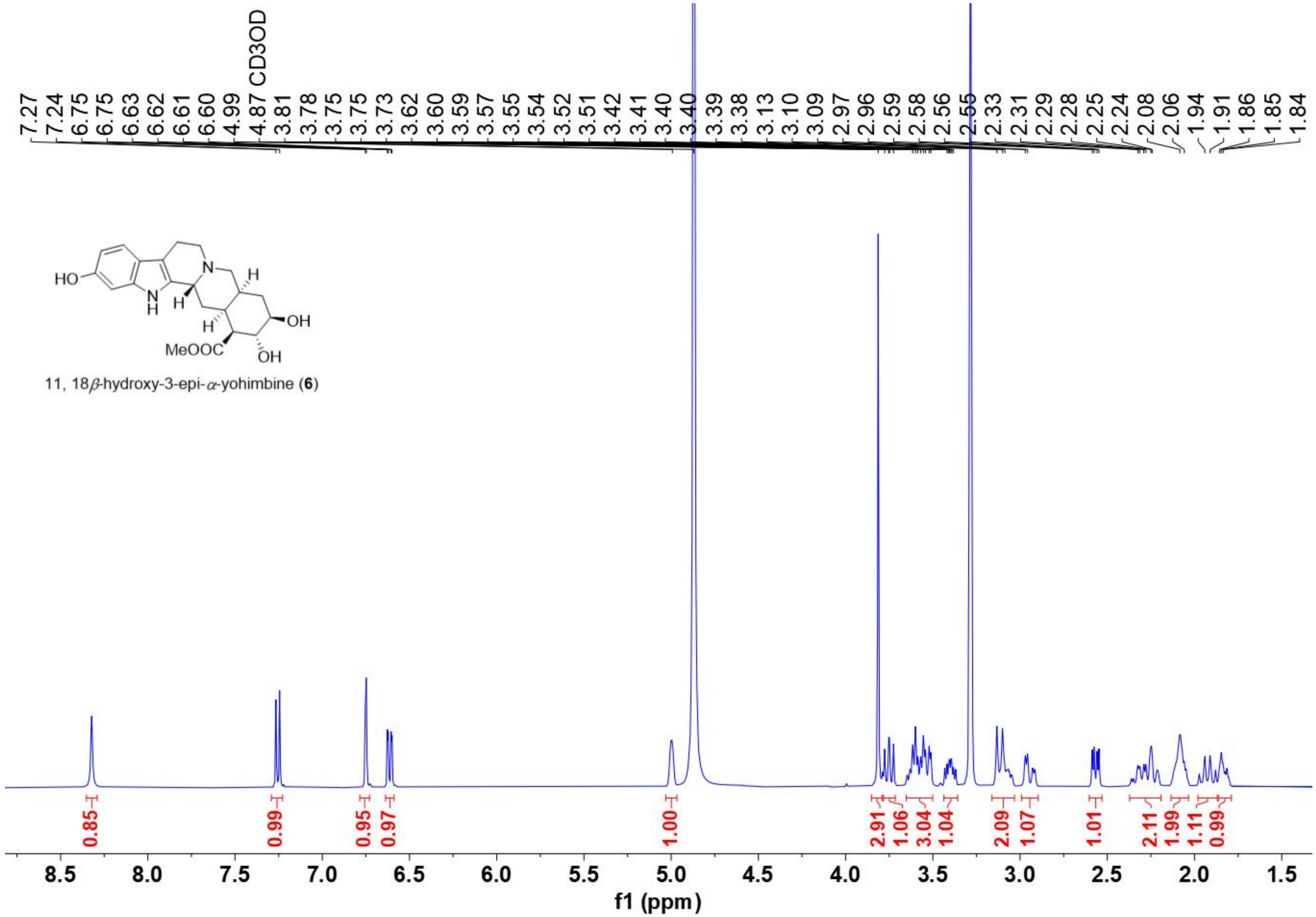

$^{1}$H NMR spectrum of **6** (recorded in $CD_3OD$ at 400 MHz)

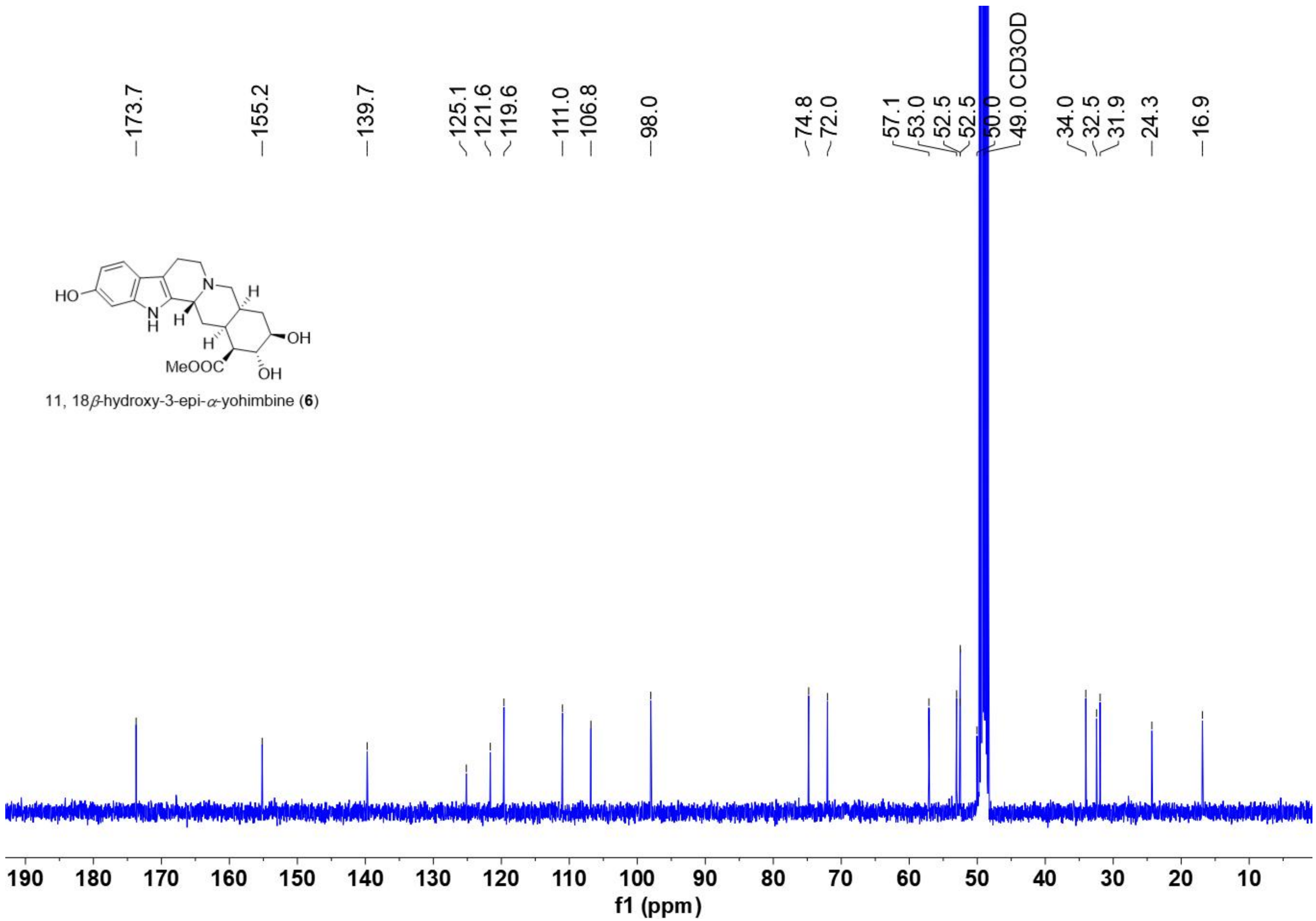

$^{13}C$ NMR spectrum of **6** (recorded in $CD_3OD$ at 100 MHz)

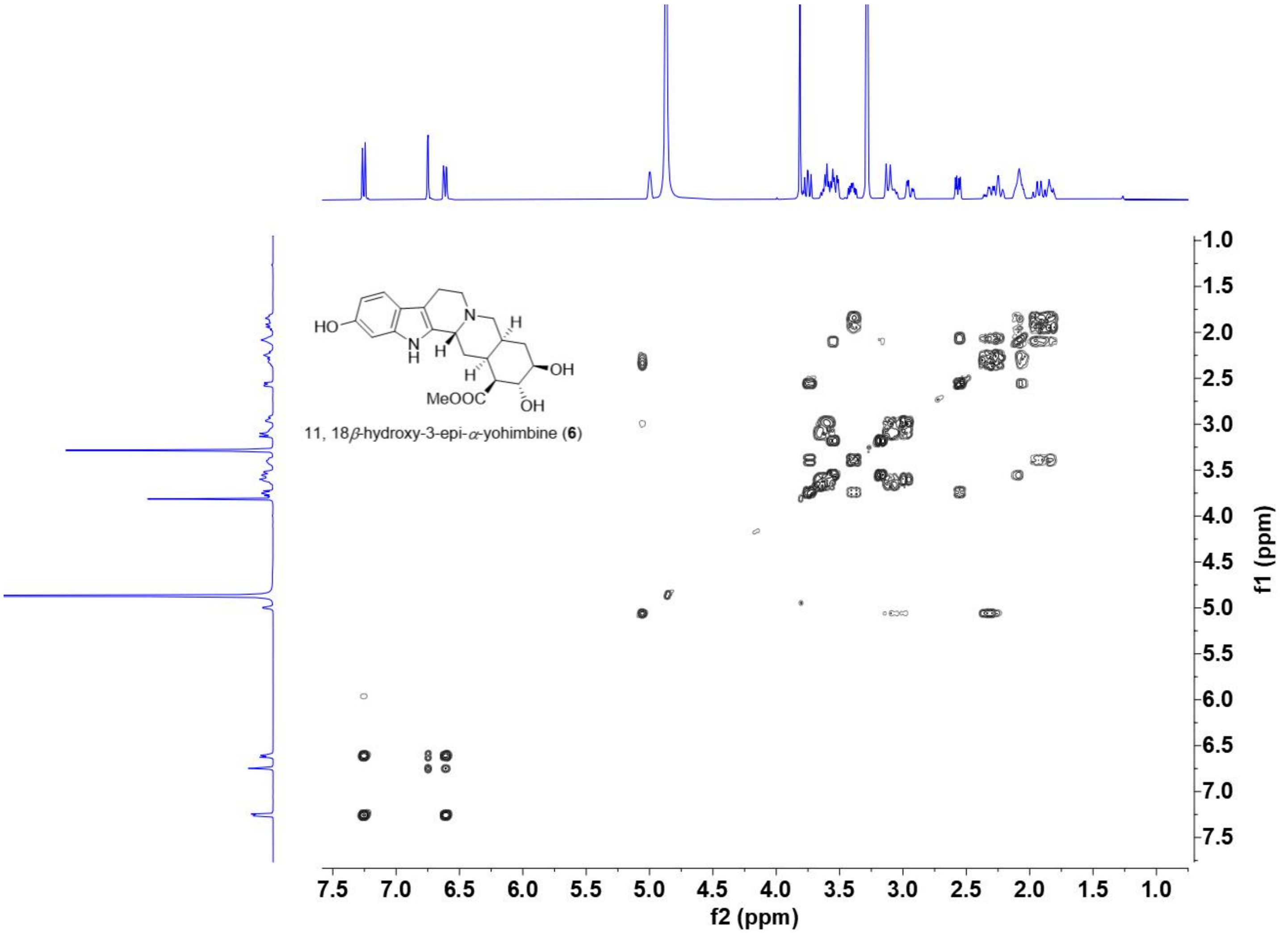

$^1$H-$^1$H COSY spectrum of **6** (recorded in $CD_3OD$ at 400 MHz)

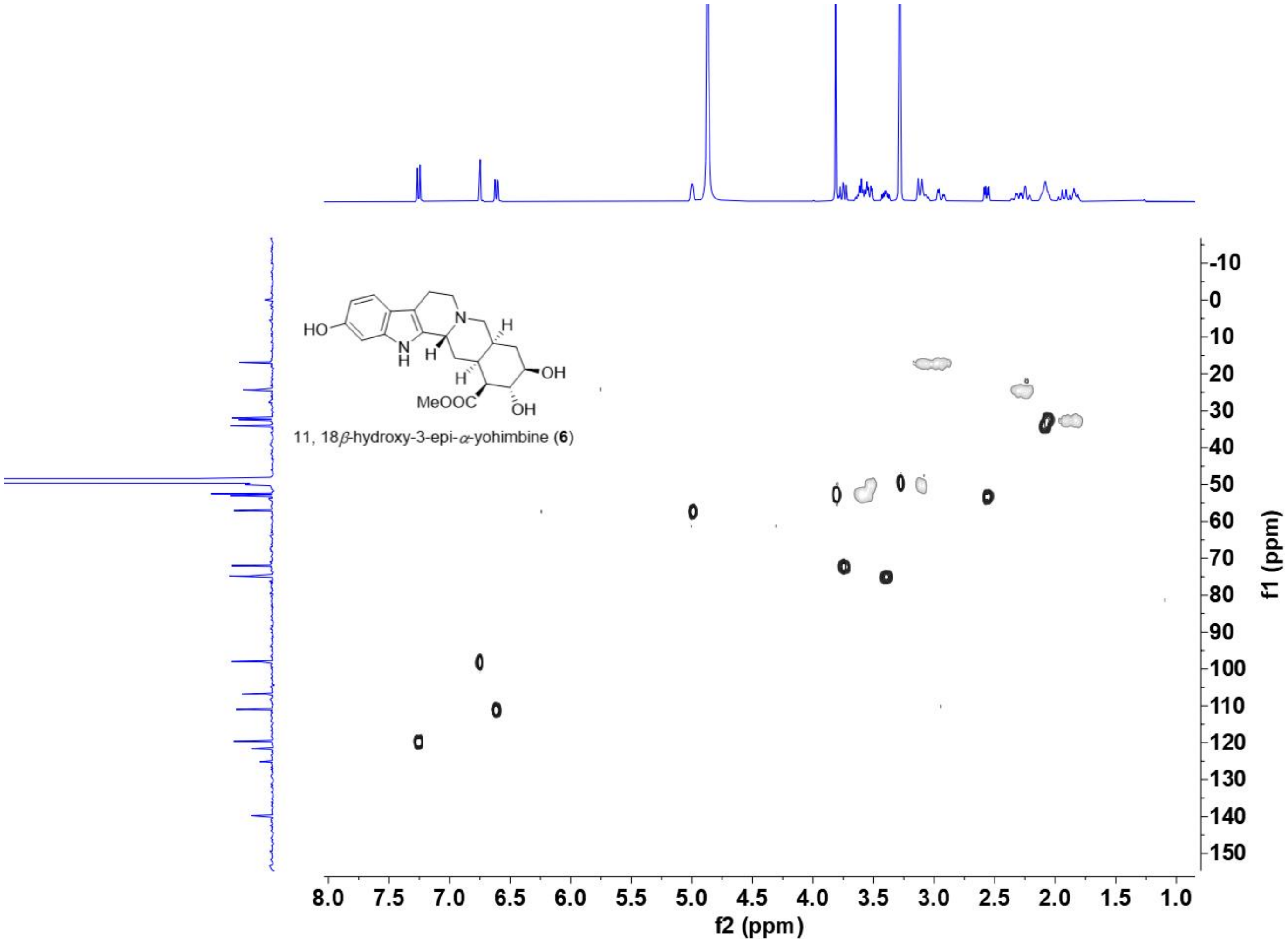

HSQC spectrum of **6** (recorded in $CD_3OD$ at 400 MHz)

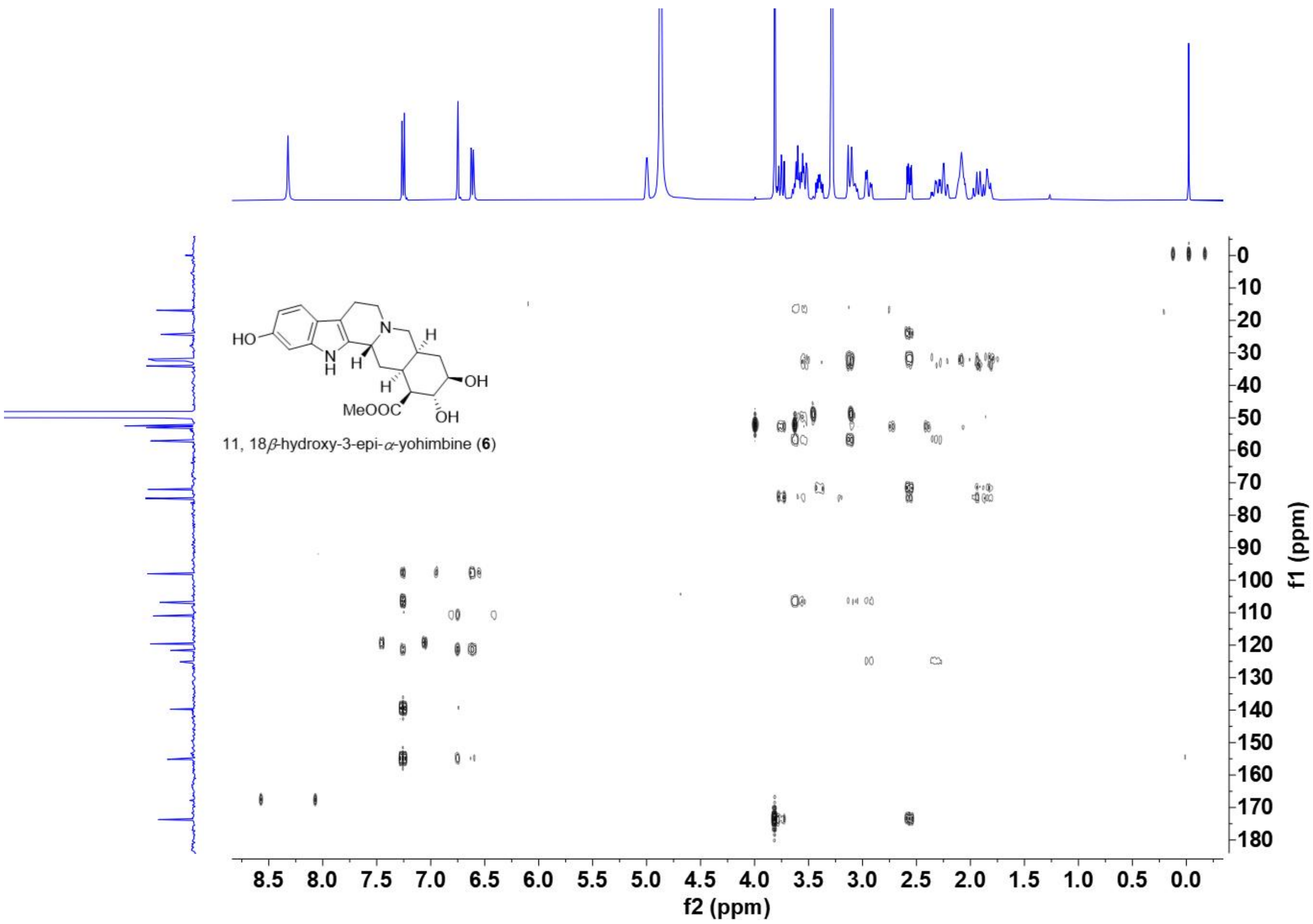

HMBC spectrum of **6** (recorded in $CD_3OD$ at 400 MHz)

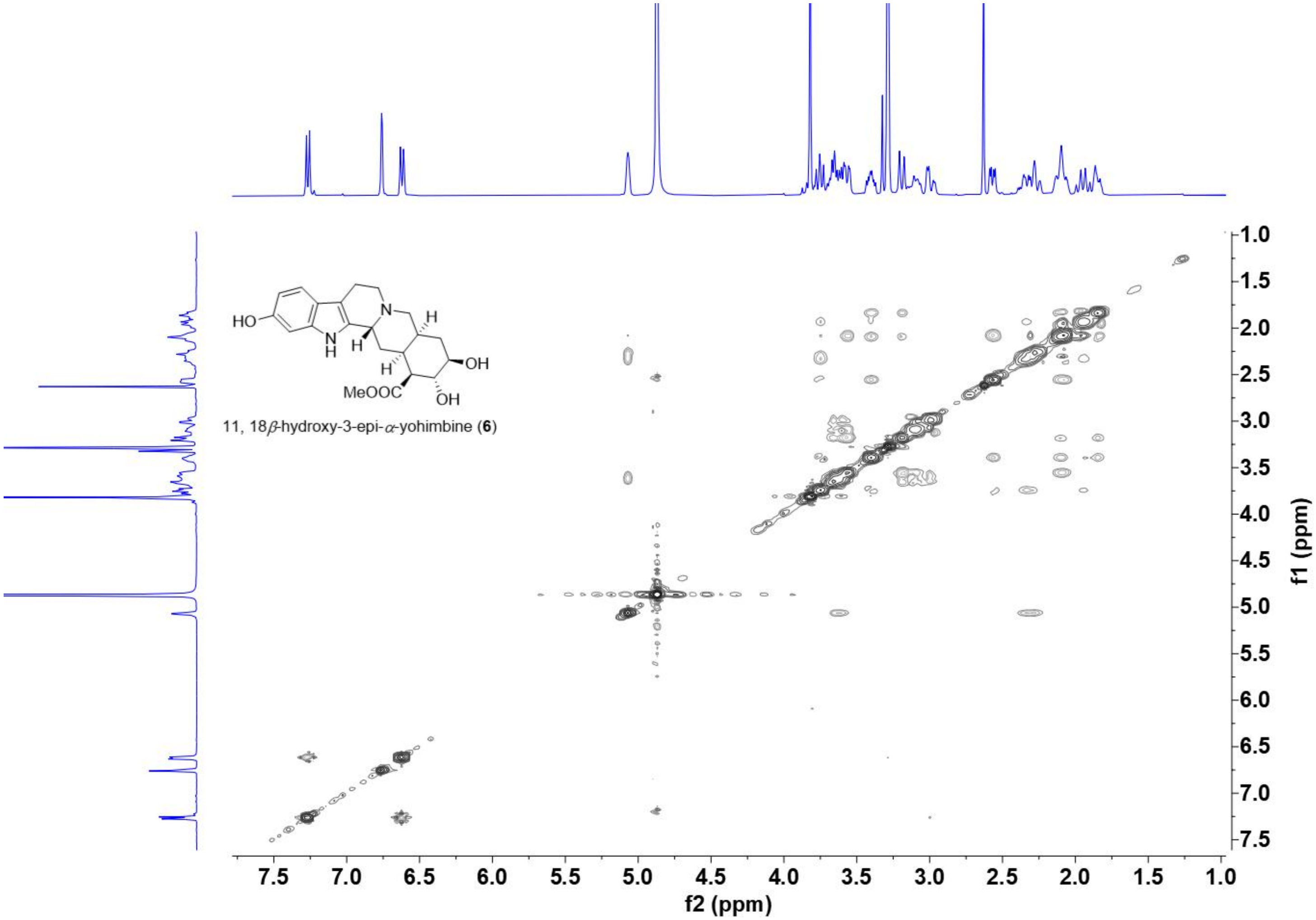

NOSEY spectrum of **6** (recorded in $CD_3OD$ at 400 MHz)

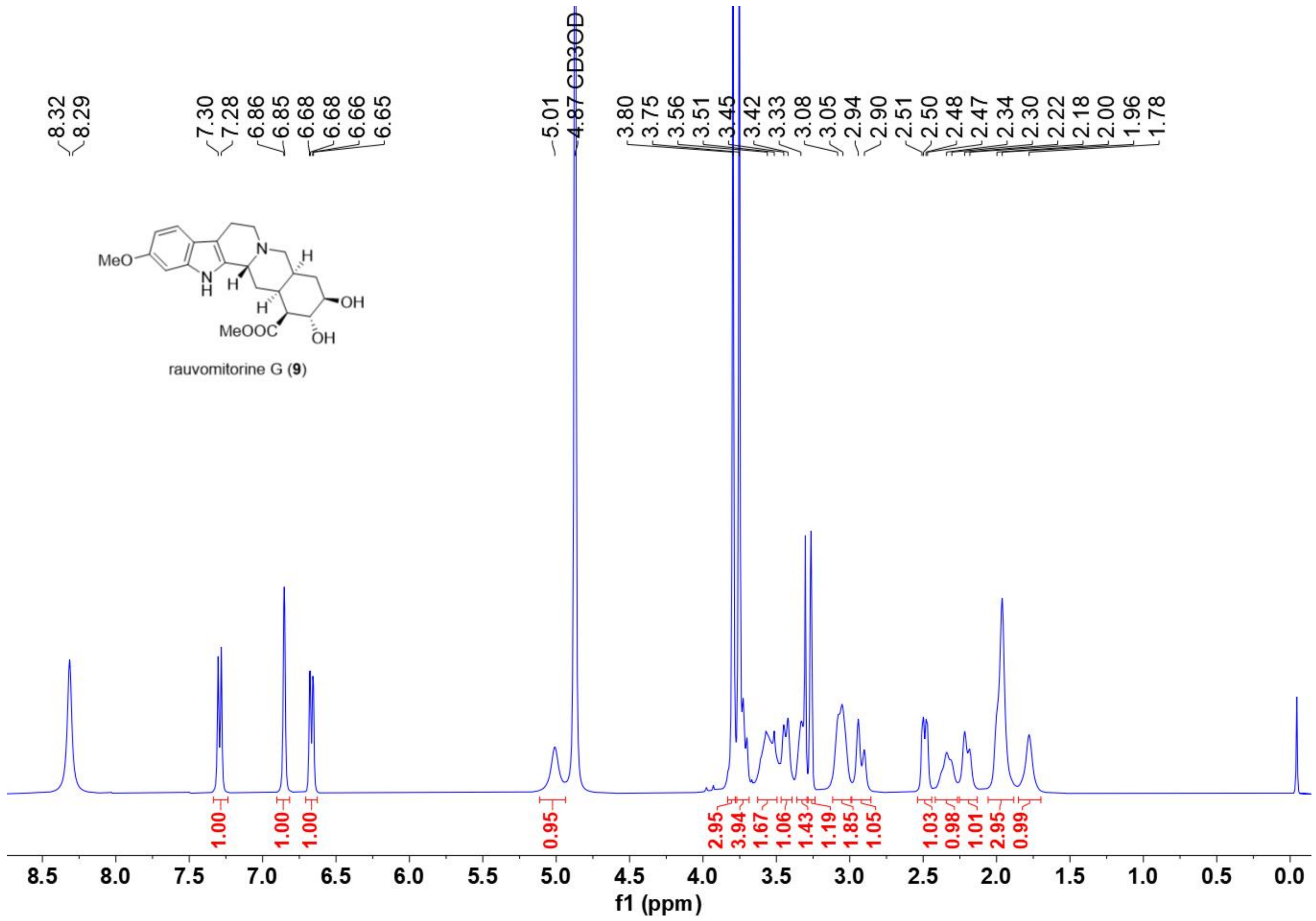

[1]H NMR spectrum of **9** (recorded in $CD_3OD$ at 400 MHz)

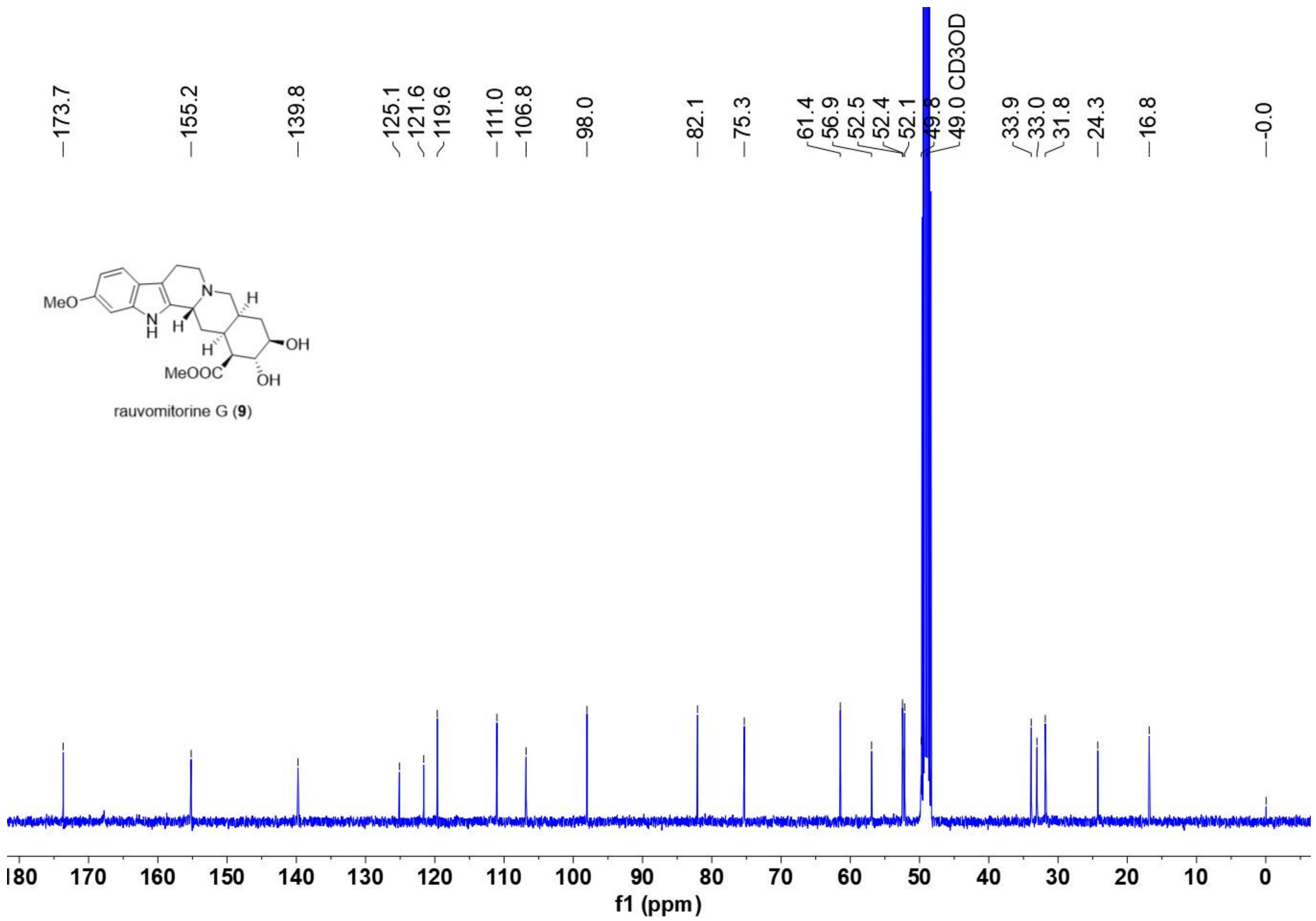

$^{13}C$ NMR spectrum of **9** (recorded in $CD_3OD$ at 100 MHz)
